# DNA-Encoded Library-Guided Discovery of a First-in-Class Molecular Glue of the NOS1AP-SCRIB Complex in Alzheimer’s Disease Patient-Derived Neurons

**DOI:** 10.64898/2026.10.12.758423

**Authors:** Baljit Kaur, Ashraf N. Abdo, Sungwoo Cho, Moustafa T. Gabr

## Abstract

NOS1AP is an adaptor for neuronal nitric oxide synthase that engages several partner proteins and is implicated in Alzheimer’s disease. Efforts to drug NOS1AP have sought to disrupt these; we instead report pharmacological stabilization of one interaction. Affinity selection of a 1.4-billion-member DNA-encoded library identified a benzimidazole-5-carboxamide chemotype, optimized by fragment-directed SAR to NOSTRA with submicromolar NOS1AP binding affinity. Of six interactions, NOSTRA enhanced only SCRIB binding, with a cooperativity factor of 30, predominantly through a 14-fold slowing of dissociation; native mass spectrometry resolved a NOS1AP:NOSTRA:SCRIB ternary complex. In healthy human iPSC-derived cortical neurons, NOSTRA engaged the endogenous complex and activated Rac1 without cytotoxicity; in Alzheimer’s patient-derived neurons, it lowered the Aβ42/Aβ40 ratio and p-tau217, reduced neurofilament light and improved network activity. With an in vitro ADME profile consistent with CNS exposure, NOSTRA is the first molecular glue for a NOS1AP complex, establishing interaction stabilization as a viable NOS1AP pharmacology.

**Table of Contents:** 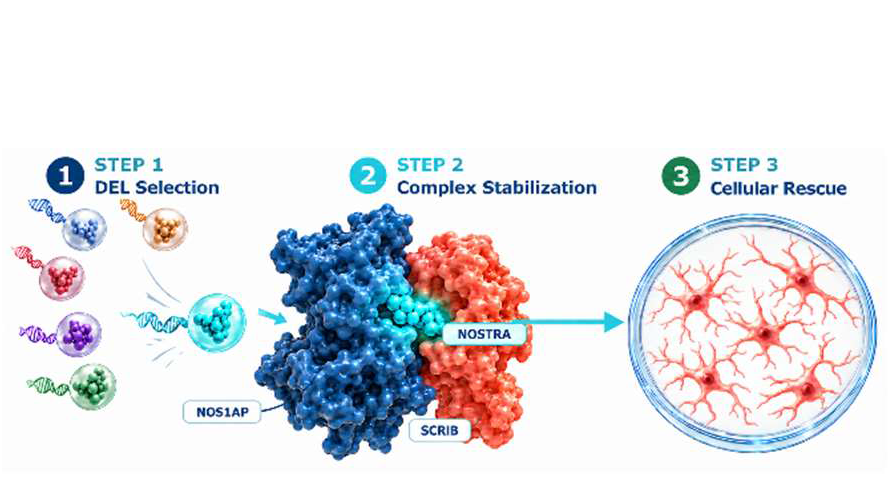

## Introduction

Nitric oxide synthase 1 adaptor protein (NOS1AP), also known as CAPON, coordinates signaling downstream of neuronal nitric oxide synthase (nNOS) and regulates processes spanning synaptic organization, neuronal excitability, and cardiac repolarization.^1,2^ NOS1AP binds nNOS through a C-terminal PDZ-binding motif, competing with postsynaptic density protein-95 (PSD-95) for the same surface and redirecting nitric oxide signaling toward distinct downstream effectors.^3,4^ Genetic and mechanistic studies link NOS1AP dysregulation to QT-interval prolongation and sudden cardiac death,^5,6^ to schizophrenia and depression,^7^ and to Alzheimer’s disease (AD), where nNOS-NOS1AP complex formation increases following amyloid-β exposure and disruption of the interaction restores memory and dendritic morphology in model mice.^8^ NOS1AP has also been shown to promote tau hyperphosphorylation and amyloid-β accumulation.^9^ NOS1AP presents a challenging target for small molecule discovery because it functions as a nonenzymatic scaffolding protein without a defined catalytic pocket, mediating its biological activities primarily through interactions with multiple protein partners.^1–4^

Pharmacological efforts against adaptor proteins have almost uniformly sought to disrupt their interactions, and our own NOS1AP programs have followed that logic.^10–13^ Here, we pursued pharmacological stabilization of a neuroprotective protein complex. Molecular glues, small molecules that increase the affinity between two proteins by binding at or near their shared interface, have expanded what is considered druggable, most visibly in targeted protein degradation.¹⁴ The same cooperative principle applies to native complexes whose assembly is itself the desired outcome, as established by fusicoccin-derived and synthetic 14-3-3 stabilizers.¹⁵,¹⁶ Selective stabilization of a protein-protein interaction offers an alternative to disrupting adaptor-mediated complexes by preferentially reinforcing a specific partnership without broadly interfering with other interactions. NOS1AP associates with nNOS, Dexras1, synapsin, carboxypeptidase E, tau, and SCRIB through distinct binding regions,^1,2^ making it a suitable model for investigating whether small molecules can selectively stabilize one interaction within a network of protein partnerships.

Among these partners, the interaction with the polarity scaffold SCRIB is distinguished by its structural basis and downstream connectivity. Richier et al. showed that NOS1AP associates directly with SCRIB through its N-terminal phosphotyrosine-binding (PTB) domain and the fourth PDZ domain of SCRIB, and that SCRIB links NOS1AP to the β-PIX/GIT1/PAK signaling complex regulating Rac activity and actin-dependent dendritic spine organization; NOS1AP expression influenced Rac activity and dendritic protrusion formation.^17^ This places the NOS1AP-SCRIB complex at a node connecting an adaptor implicated in AD to the cytoskeletal machinery whose dysfunction underlies dendritic spine loss, among the structural correlates most tightly associated with cognitive decline.^18^

We therefore hypothesized that pharmacological stabilization of the NOS1AP-SCRIB interaction could reinforce a synaptic scaffolding network that regulates Rac1-dependent cytoskeletal dynamics and neuronal connectivity, thereby improving neuronal resilience and network function under disease-associated stress. Although the NOS1AP-SCRIB-β-PIX/GIT1/PAK–Rac axis is well established biologically, whether stabilizing this complex is therapeutically beneficial in AD has not been investigated. Rac1 signaling is also context dependent: excessive or mislocalized activity has been implicated in amyloid processing and tau pathology,^19^ whereas appropriately regulated activity supports synaptic maintenance,^20^ and NOS1AP overexpression can impair mature dendritic spine morphology.^21^ In this study, we identify NOSTRA, a small molecule stabilizer of the NOS1AP-SCRIB interaction, and investigate whether selectively reinforcing this partnership can improve neuronal function in AD models without broadly enhancing NOS1AP activity or Rac1 signaling.

Our laboratory has previously generated NOS1AP-directed molecules through distinct discovery strategies.^10–13^ Affinity selection-mass spectrometry identified small molecule NOS1AP binders as well as small molecule inhibitors of NOS1AP-nNOS interaction.^11^ Phage display yielded NOS1AP-targeted cyclic peptides that reduced amyloid-β-induced toxicity and tau phosphorylation in cellular AD models.^12^ Most recently, we reported CAPTOR-1, a small molecule inhibitor of the NOS1AP-tau interaction.^13^ Thus, each prior program was designed to disrupt a NOS1AP interaction based on screening preexisting compound libraries. The central conceptual advance of the present study is to instead pharmacologically stabilize a defined NOS1AP complex. We used DNA-encoded library (DEL) selection as an enabling discovery strategy. Because DEL hits are recovered as DNA conjugates and require de novo resynthesis for validation and optimization,^22^ this platform provides full synthetic control over the resulting series and access to chemical space not represented in conventional screening collections.^23,24^

Selection of a 1.4-billion-member DEL against recombinant NOS1AP identified an enriched benzimidazole-5-carboxamide chemotype, which was resynthesized off-DNA and advanced through fragment-directed structure-activity relationship (SAR) studies to give NOSTRA, a submicromolar NOS1AP binder. Screening NOSTRA across six established NOS1AP interactions showed that it did not inhibit any of them; rather, it selectively enhanced the NOS1AP-SCRIB interaction. We characterize this activity as cooperative complex stabilization by orthogonal biophysics, demonstrate formation of a discrete ternary complex by native mass spectrometry, show that the mechanism operates on the endogenous complex in human iPSC-derived cortical neurons, and evaluate its consequences in neurons derived from AD patients. NOSTRA is, to our knowledge, the first small molecule reported to stabilize a NOS1AP complex.

## Results and Discussion

### Identification of a NOS1AP-Binding Chemotype by DEL Selection

To identify small molecules capable of engaging NOS1AP, we performed an affinity-based DEL selection (Figure 1). Recombinant His-tagged human NOS1AP was immobilized on Ni-NTA magnetic beads and used as bait against a 1.4-billion-member DNA-encoded small-molecule library. Immobilization efficiency and protein integrity were confirmed by SDS-PAGE analysis of input, flow-through, wash, bead-bound, and elution fractions prior to selection.

**Figure 1.**
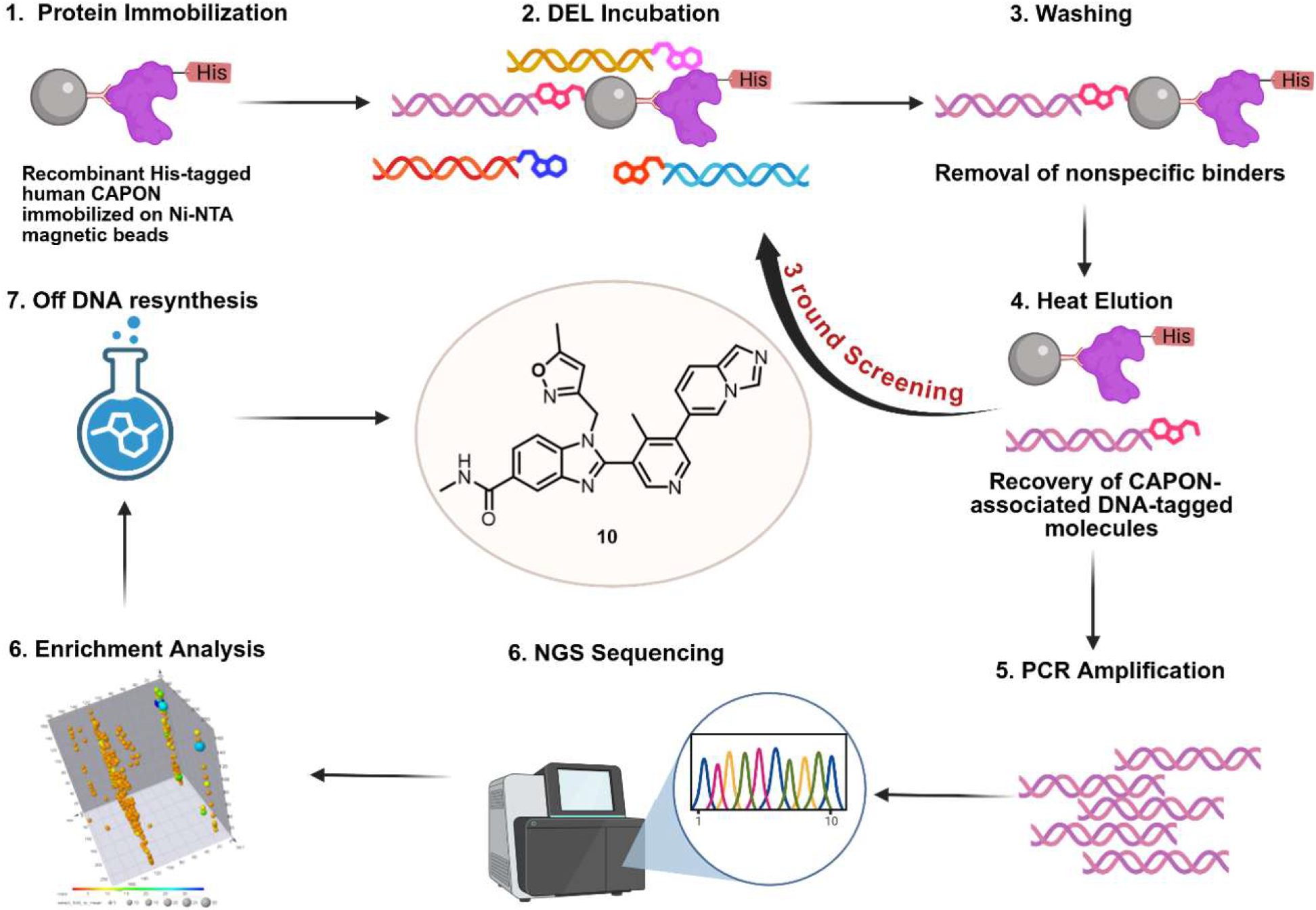
DNA-encoded library selection against NOS1AP. Recombinant His-tagged NOS1AP was immobilized on Ni-NTA magnetic beads and incubated with a 1.4-billion-member DNA-encoded small molecule library. Following incubation, successive stringent washes removed nonspecific binders, and retained DNA-tagged compounds were recovered by heat-mediated elution. Associated DNA barcodes were amplified by PCR and decoded by next-generation sequencing. Enriched library members were identified by comparing sequencing counts from the target selection with those from a no-target control. The corresponding off-DNA compound **10** was synthesized and evaluated by biophysical binding assays as the starting point for medicinal chemistry optimization.

Immobilized NOS1AP was incubated with the encoded library to allow target-associated compounds to bind, followed by stringent washing to remove weakly associated and nonbinding members. Bound compounds were eluted, and their DNA tags were amplified by PCR and analyzed by next-generation sequencing. Enrichment was assessed by comparing sequencing counts from the target selection with those from a no-target control selection performed in parallel.

Sequencing analysis revealed selective enrichment of several building blocks among NOS1AP-associated library members, consistent with productive target engagement across multiple chemotypes. Building block BB3-481 showed the most pronounced enrichment and was prioritized for follow-up on the basis of enrichment magnitude and synthetic accessibility. The corresponding library member was resynthesized off-DNA to afford compound **10** (Figure 1), which was carried forward for biophysical evaluation.

Microscale thermophoresis (MST) confirmed direct, DNA-independent binding of **10** to NOS1AP. The interaction was weak and did not reach saturation across the accessible concentration range, placing the dissociation constant above 50 µM (Supplementary Fig. 90). Weak affinity is characteristic of primary DEL hits against protein-protein interaction surfaces lacking a conventional binding pocket, and compound **10** was judged a chemically tractable starting point on the basis of its defined synthetic origin, modular three-fragment architecture, and amenability to systematic diversification. It was therefore selected as the lead chemotype for SAR studies.

### Synthesis of the DEL Hit and an Unexpected Isoxazole-to-Pyrazole Conversion

Compound **10** was prepared as outlined in Scheme 1. 4-Fluoro-*N*-methyl-3-nitrobenzamide (**1**) was reacted with (5-methylisoxazol-3-yl)methanamine (**2**) in water at 80 °C for 4 h to afford intermediate **3**. Reduction of the nitro group with SnCl_2_ in ethanol gave aniline **4**, which was condensed with 3-bromoisonicotinaldehyde (**5**) in ethanol in the presence of sodium metabisulfite to furnish the aromatized benzimidazole **6**. Suzuki–Miyaura coupling of **6** with imidazo[1,5-a]pyridin-6-ylboronic acid (**9**) in dioxane/water with K_2_CO_3_ at 90 °C delivered the target compound **10**.

**Scheme 1.**
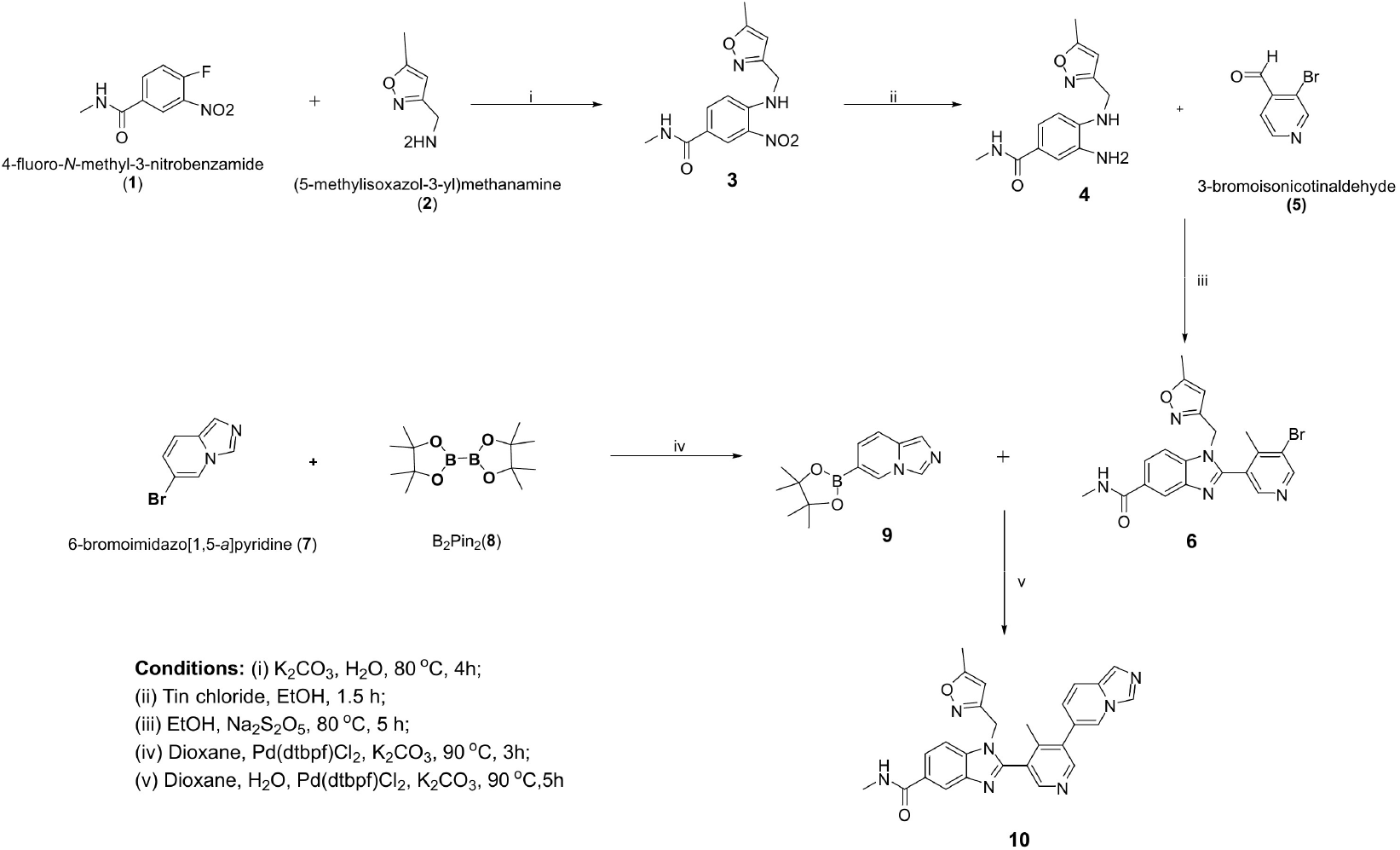
Synthesis of the DEL-derived hit compound **10**.

Scale-up of this route for analogue synthesis was limited by the nitro reduction, at which the isoxazole ring proved labile. Several reducing systems were evaluated. Hydrogenation over Pd/C degraded the isoxazole ring; Fe/AcOH gave sluggish conversion and, on prolonged reaction, multiple by-products; and although SnCl₂/HCl afforded the desired aniline, it required a laborious work-up poorly suited to parallel synthesis.

Treatment with hydrazine in the presence of Pd/C in ethanol instead effected clean reduction within 30 min at 50 °C. Characterization of the product revealed that, in addition to nitro reduction, the isoxazole ring had been converted to a pyrazole (Scheme 2), consistent with reductive N–O bond cleavage and recyclization. Carrying this intermediate through the sequence afforded the pyrazole congener **13**, which bound NOS1AP with a *K*_D_ of 77 µM, comparable to the isoxazole parent **10** within the resolution of the assay, indicating that the heterocycle substitution is approximately affinity-neutral at this position.

We nonetheless adopted the pyrazole as the Fragment C template for the remainder of the campaign, for three reasons. First, the hydrazine/Pd-C reduction was operationally superior to every alternative examined, proceeding to completion in 30 min with a simple filtration work-up and thereby enabling the parallel synthesis required for systematic SAR. Second, **13** displayed improved aqueous solubility relative to **10**. Third, the pyrazole is generally associated with greater metabolic stability than the isoxazole, whose N–O bond is a recognized liability. Affinity gains were therefore pursued through the aromatic regions of the scaffold rather than through this heterocycle.

**Scheme 2.**
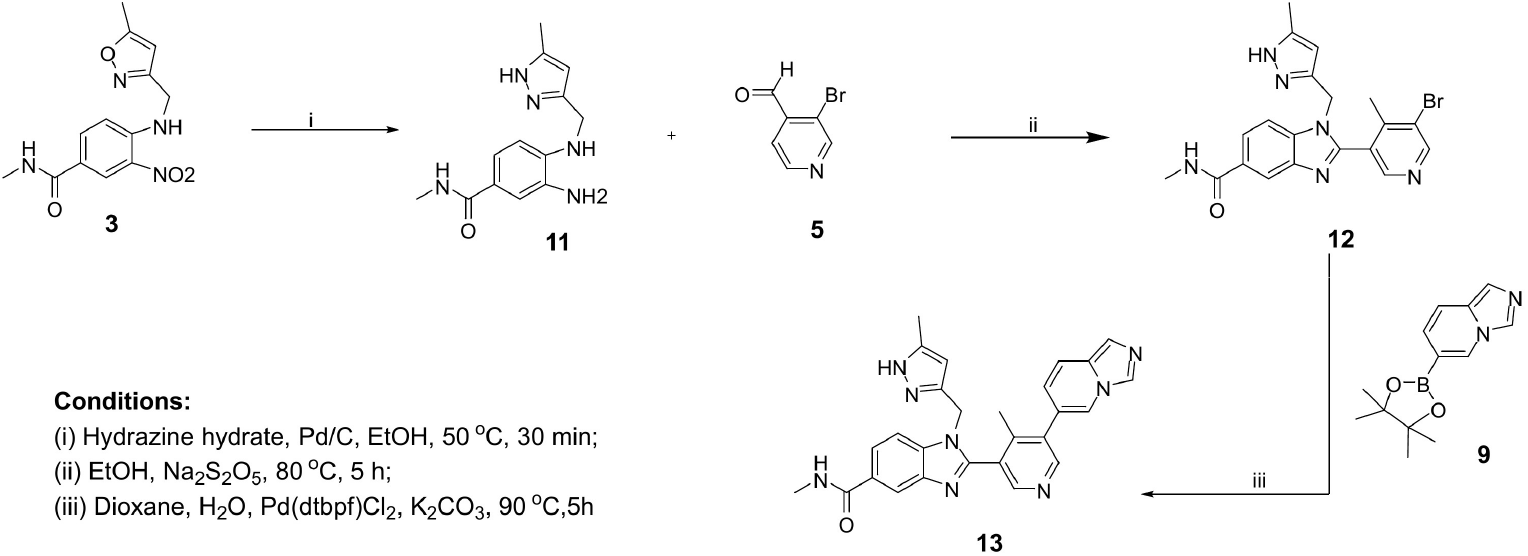
Hydrazine/Pd-C reduction of **3** with concomitant isoxazole-to-pyrazole conversion, affording **13**.

### Fragment-Directed SAR Studies Afford NOSTRA

The scaffold of **13** comprises three modular regions (Figure 2): a conserved *N*-methylbenzo[*d*]imidazole-5-carboxamide core; a C2-linked bicyclic system consisting of a central heteroaryl spacer (Fragment B) and a distal heteroaryl pharmacophore (Fragment A); and an N1-linked 5-methylpyrazolylmethyl terminal cap (Fragment C). We optimized these regions sequentially, fixing the best substituent at each stage before proceeding, so that each round interrogated a single variable against a defined background.

**Figure 2.**
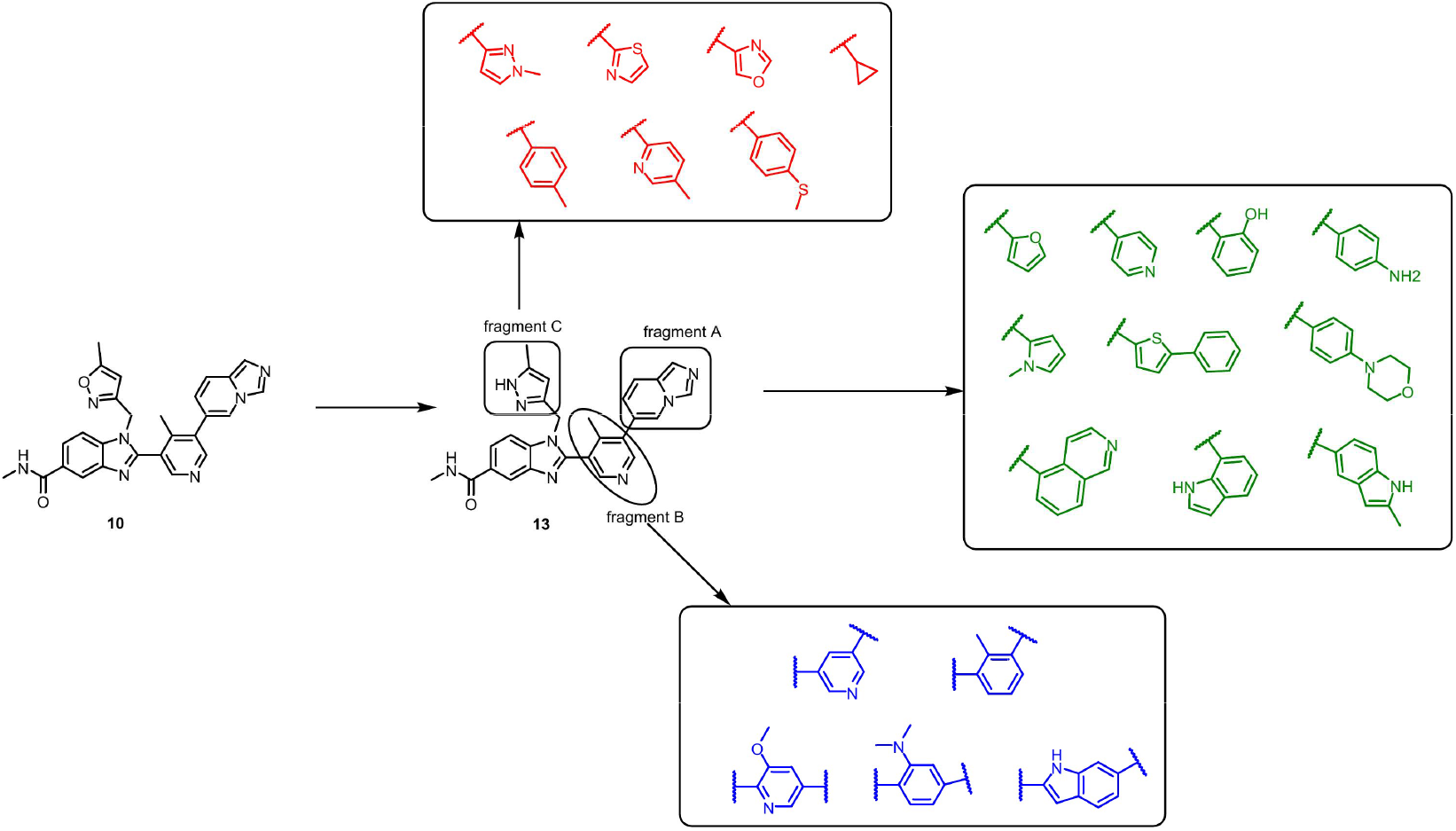
Fragment-directed SAR strategy. The scaffold of **13** is divided into the distal heteroaryl pharmacophore (Fragment A), the central heteroaryl spacer (Fragment B), and the terminal cap (Fragment C), each diversified in a separate round against a fixed background.

### Fragment A

The distal imidazo[1,5-a]pyridine of **13** was replaced by a structurally diverse set of monocyclic and bicyclic aromatic and heteroaromatic substituents (**24**-**33**), selected to vary heteroatom placement, hydrogen-bonding capability, aromatic surface area, and electronic distribution while preserving a planar framework. Compounds **24**-**33** were prepared by Suzuki– Miyaura coupling of intermediate **12** with the corresponding boronic acids **14**-**23** using Pd(dtbpf)Cl_2_ and K_2_CO_3_ in dioxane/H_2_O (Scheme 3).

This region proved highly sensitive to substituent identity (Table 1). Furyl (**24**), pyridyl (**25**), and 4-hydroxyphenyl (**26**) derivatives gave moderate binding (*K*_D_ = 43, 44, and 47 µM), while 4-amino-2-methylphenyl (**27**) and 1-methylpyrrolyl (**28**) were weaker (85 and 80 µM). In contrast, the thiophenyl (**29**) and isoquinolinyl (**31**) analogues improved affinity by approximately an order of magnitude, to 9.4 and 9 µM respectively. The 2-methyl-1H-indolyl analogue **33** retained moderate affinity (48 µM), whereas **30** and **32** gave no measurable binding under the conditions tested. The preference for extended bicyclic systems capable of π-stacking identified thiophenyl and isoquinolinyl as the productive Fragment A motifs.

**Scheme 3.**
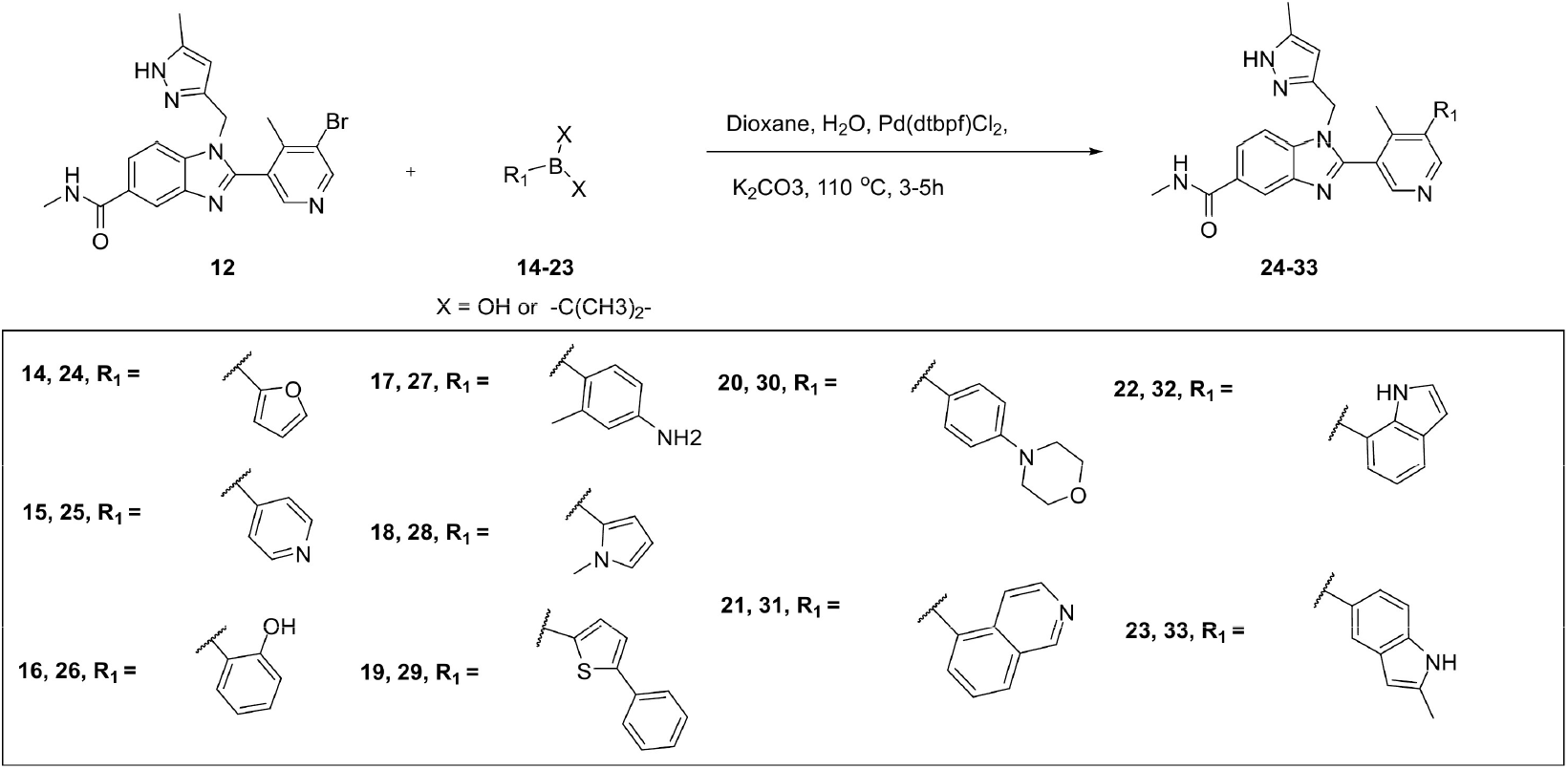
Synthesis of derivatives of **13** (compounds **24**-**33**).

**Table 1.** NOS1AP binding affinities of compound 10 and its derivatives, determined by MST.

| Cpd | $K_D$ ($\mu\text{M}$) | Cpd | $K_D$ ($\mu\text{M}$) | Cpd | $K_D$ ($\mu\text{M}$) |
| --- | --- | --- | --- | --- | --- |
| **10** | $>50^a$ | **30** | ND<sup>b</sup> | **49** | **0.50** |
| **13** | 77 | **31** | 9 | **50** | 83 |
| **24** | 43 | **32** | ND | **51** | 26 |
| **25** | 44 | **33** | 48 | **52** | 65 |
| **26** | 47 | **44** | 50 | **53** | ND |
| **27** | 85 | **45** | ND | **76<sup>c</sup>** | 163 |
| **28** | 80 | **46** | ND | **77<sup>c</sup>** | ND |
| **29** | 9.4 | **47** | ND | **78<sup>c</sup>** | 79 |
|  |  | **48** | ND | **79<sup>c</sup>** | 116 |
|  |  |  |  | **80<sup>c</sup>** | 122 |
|  |  |  |  | **81<sup>c</sup>** | 114 |
|  |  |  |  | **82<sup>c</sup>** | 138 |

Values are means of three independent measurements ( $n = 3$ ). <sup>a</sup>Binding detected but not saturating across the accessible concentration range; $K_D$ could not be accurately determined. <sup>b</sup>ND, not determined, no measurable binding under the assay conditions. <sup>c</sup>Compounds **76-82** are constructed on a 2-(isoquinolin-5-yl)pyridin-4-yl core, which differs in substitution pattern from the 5-(isoquinolin-5-yl)pyridin-3-yl core of **49**; their affinities are not directly comparable with the remainder of the series.

### Fragment B

With Fragment A fixed as either thiophenyl or isoquinolinyl, the central 4-methylpyridyl spacer was varied. Intermediates **39**-**43** were prepared by condensation of **11** with aldehydes **34**-**38** in ethanol with Na_2_S_2_O_5_ at 80 °C (Scheme 4), then coupled with boronic acids **19** and **21** to furnish **44**-**48** and **49**-**53** (Scheme 5).

**Scheme 4.**
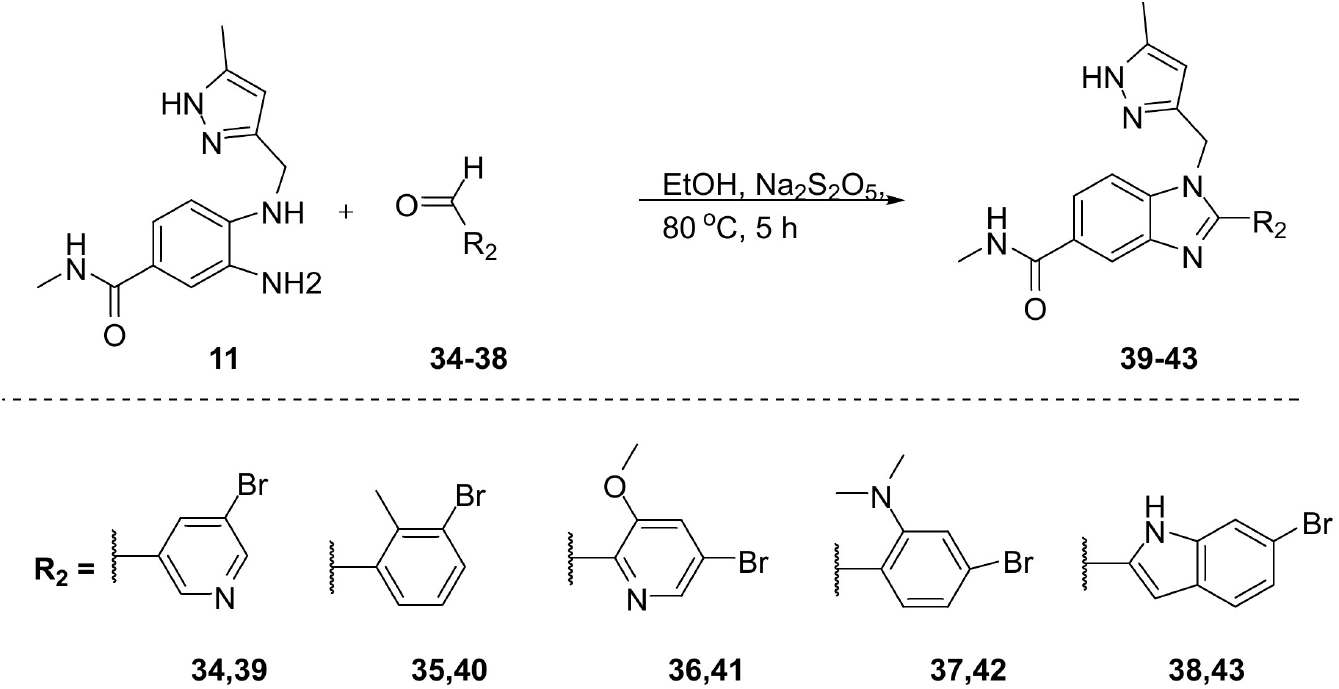
Synthesis of intermediates **39**-**43**.

Binding depended strongly on the combination of the two regions rather than on either independently. Within the isoquinolinyl series, removal of the 4-methyl substituent from the central pyridine gave **49**, the most potent compound of the campaign at *K*_D_ = 0.50 µM. Other modifications of this region were substantially less favorable: the 3-methoxypyridin-2-yl analogue **51** bound 52-fold more weakly (26 µM), the 2-methylphenyl analogue **50** 166-fold more weakly (83 µM), and the *N,N*-dimethylaminophenyl derivative **52** 130-fold more weakly (65 µM), while the indolyl analogue **53** gave no measurable binding. Within the thiophenyl series the corresponding des-methyl analogue **44** was only moderately active (50 µM), and **45**-**48** gave no measurable *K*_D_, indicating that the gains observed with thiophenyl in the first round did not survive recombination.

**Scheme 5.**
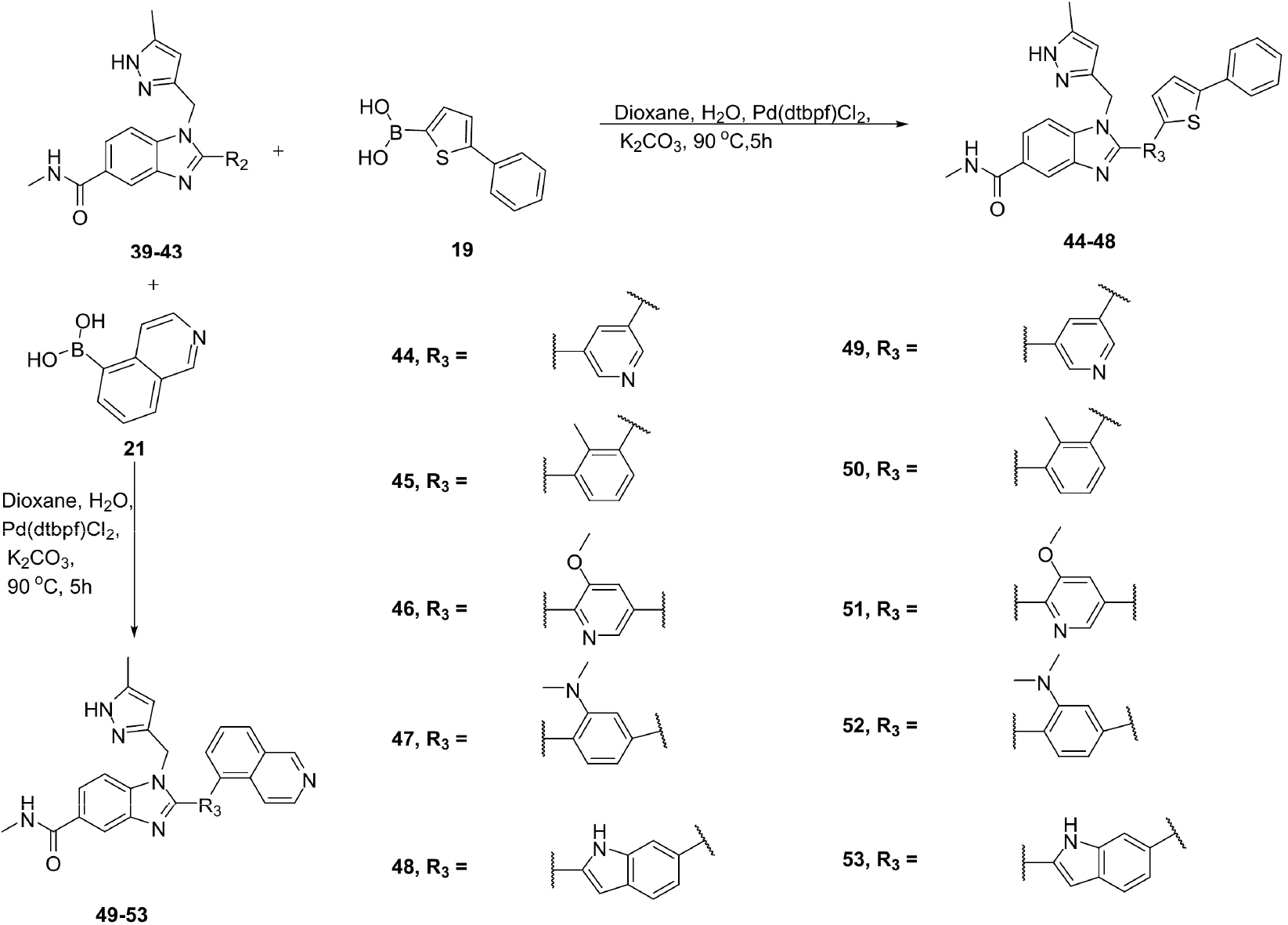
Synthesis of compounds **44**-**53**.

### A single atom increment with context dependence

Compounds **31** and **49** differ by exactly one heavy atom, the 4-methyl group on the central pyridine, and are otherwise identical, making this the only rigorous single-point comparison in the series. Its deletion improves affinity 18-fold (from 9 to 0.50 µM), identifying the position adjacent to the inter-ring bond as sterically intolerant in the isoquinolinyl context. The same deletion has the opposite effect in the thiophenyl series. Compounds **29** and **44** likewise differ only by the 4-methyl group, and there its removal reduces affinity 5-fold. The consequence of a given modification at Fragment B therefore depends on the identity of Fragment A, consistent with the two regions occupying a single contiguous binding surface in which the permitted geometry of the central ring is set by the distal pharmacophore. The remaining comparisons in this round each involve two simultaneous changes, **49** versus **50** alters both a ring heteroatom and a methyl substituent, and **49** versus **51** alters both the ring attachment position and adds a methoxy group, and we therefore do not interpret them as isolating single interactions.

### Fragment C

The terminal 5-methylpyrazolylmethyl cap was replaced by 1-methyl-1H-pyrazolyl, thiazolyl, oxazolyl, cyclopropyl, p-methylbenzyl, 4-(methylthio)benzyl, and 5-methylpyridin-2-yl groups (**76**-**82**), prepared by reaction of **1** with amines **54**-**60** to give intermediates **61**-**67**, reduction to **68**-**74**, and condensation with aldehyde **75** (Scheme 6).

**Scheme 6.**
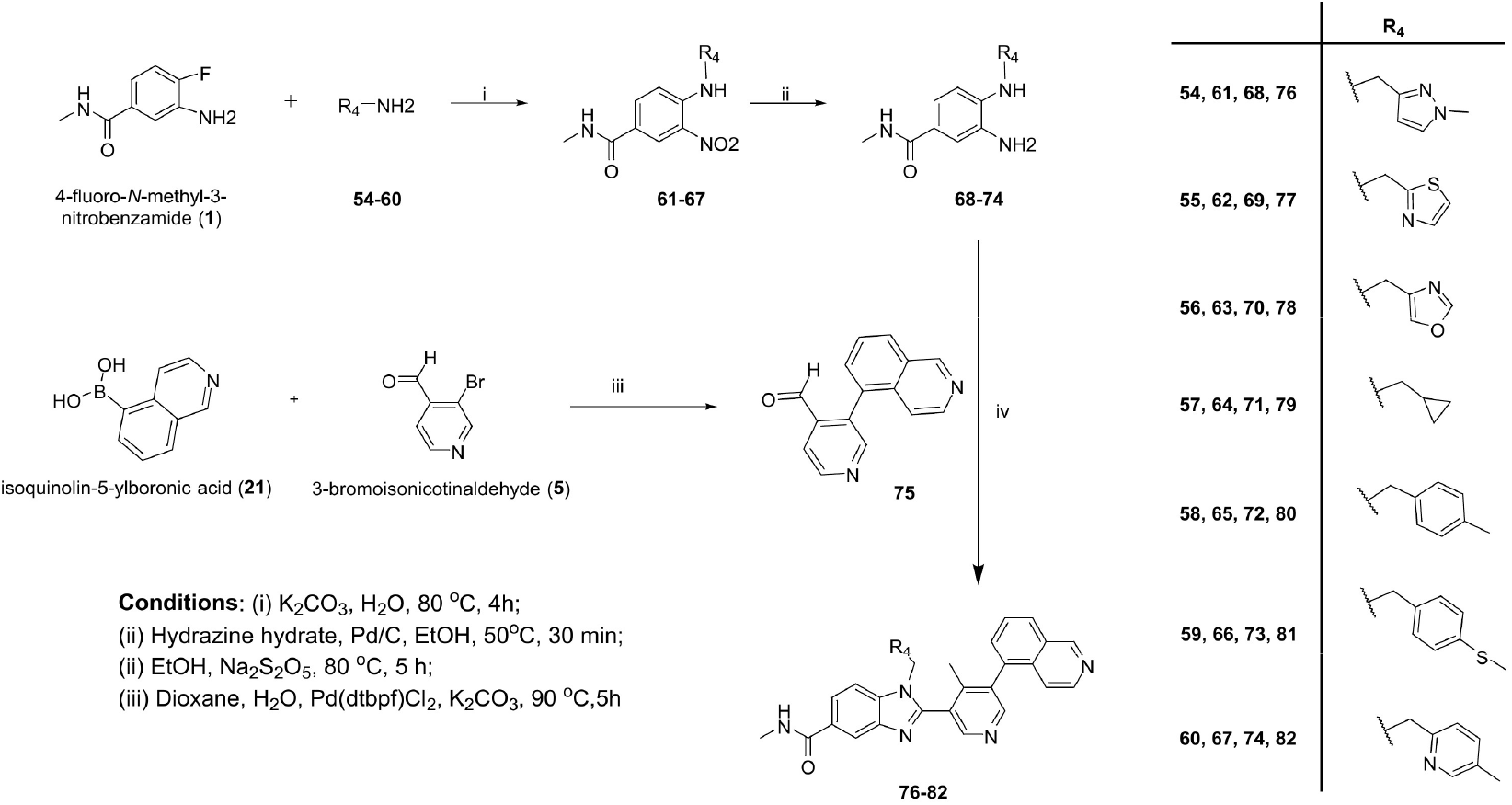
Synthesis of compounds **76**-**82**.

All seven compounds bound NOS1AP weakly, with measurable *K*_D_ values of 79-163 µM and no measurable binding for the thiazolyl derivative **77** (Table 1). This round, however, does not isolate the contribution of the terminal cap. The compounds of this series are built on a central ring bearing a different substitution pattern from that of NOSTRA, 2-(isoquinolin-5-yl)pyridin-4-yl rather than 5-(isoquinolin-5-yl)pyridin-3-yl, so that **49** and **76** are constitutional isomers rather than analogues.

Because every compound in this round carries both the modified cap and the altered core, the two variables are completely confounded across the dataset, and the loss of affinity cannot be apportioned between them. We therefore conclude only that no compound in this round improved on **49**, and we do not infer that the terminal cap is intolerant to substitution. Resolving its contribution would require synthesizing cap variants on the NOSTRA core itself. Across the three rounds, optimization improved NOS1AP binding from a hit too weak for accurate determination (**10**, *K*_D_ > 50 µM) to a submicromolar lead (**49**, *K*_D_ = 0.50 µM). Compound **49** was designated **NOSTRA** and selected for mechanistic characterization.

### NOSTRA Binds NOS1AP with Submicromolar Affinity

NOSTRA (Figure 3a) binding to NOS1AP was characterized by two orthogonal label-free methods. MST afforded a *K*_D_ value of 0.50 ± 0.18 µM, with well-defined upper and lower plateaus across a three-log concentration range (Figure 3b). Surface plasmon resonance (SPR) with NOS1AP immobilized on the sensor surface and NOSTRA injected across a 0.125-8 µM series gave *K*_D_ = 0.93 ± 0.21 µM (n = 3, Figure 3c). Global fitting of the sensorgrams to a 1:1 interaction model returned k_on_ = 3.4 × 10^4^ M^-1^ s^-1^ and k_off_ = 3.1 × 10^-2^ s^-1^, with a saturating response of approximately 23 RU. The two methods agree to within 1.9-fold, establishing NOSTRA as a submicromolar NOS1AP binder and confirming that the affinity gains achieved through fragment-directed optimization are reproduced on an orthogonal platform. An independent SPR steady-state analysis of the equilibrium binding responses returned *K*_D_ = 0.77 ± 0.18 µM, in agreement with the kinetic fit.

**Figure 3.**
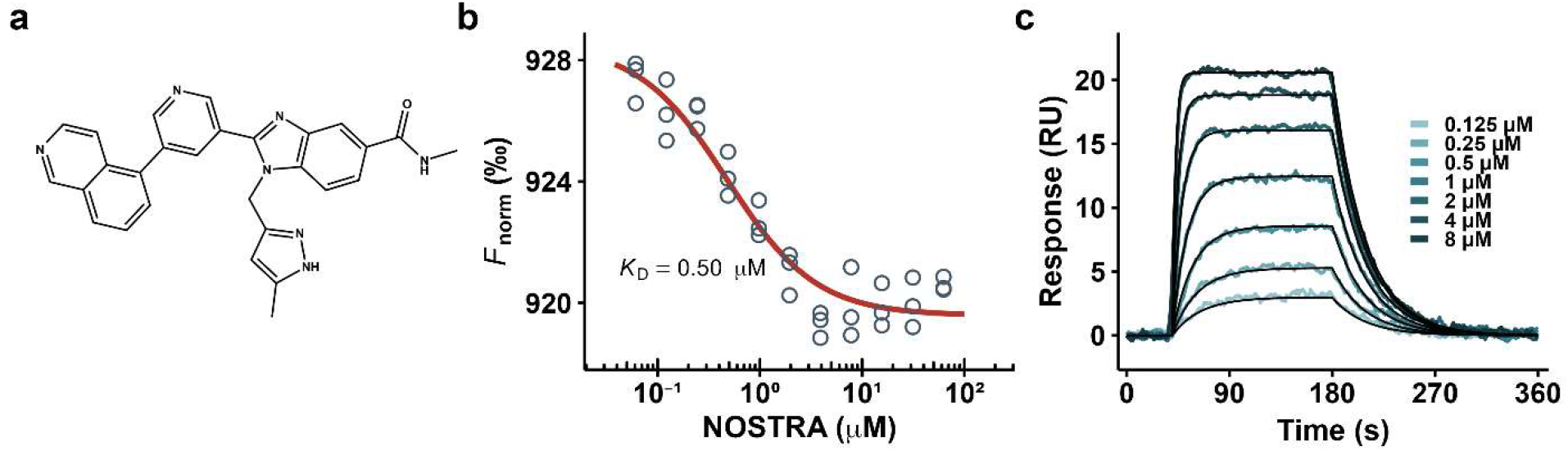
NOSTRA binds NOS1AP with submicromolar affinity. (**a**) Chemical structure of NOSTRA. (**b**) MST dose-response; all individual replicate points from three independent experiments are shown with the fitted 1:1 isotherm. (**c**) SPR sensorgrams for a 0.125-8 µM NOSTRA injection series over immobilized NOS1AP (colored) with the global 1:1 kinetic fit overlaid (black); representative of three replicates.

### Mixed-Solvent Simulations Nominate a Candidate Binding Site on the NOS1AP PTB Domain

The region of NOS1AP responsible for binding this chemotype was unknown: no experimental structure of the protein is available, and no site-directed data constrained the search. We therefore mapped the surface using mixed-solvent molecular dynamics, which ranks candidate ligand-binding hotspots by the occupancy of small organic probes and requires neither a pre-specified site nor an empirical scoring function.

We first asked which regions of NOS1AP could host a small-molecule pocket at all. In the AlphaFold model (AF-O75052-F1), only two segments are confidently folded: a phosphotyrosine-binding (PTB) domain spanning residues 15–176 (162 residues, 32% of the sequence; mean pLDDT 92.7), and a 77-residue segment (288–364; mean pLDDT 94.5) that is 96% helical with no β-strand content, consistent with a single coiled coil (Figure 4a). The remaining 53% of the protein, including both known nNOS-binding elements, is predicted disordered; no residue between 400 and 506 exceeds pLDDT 70. SiteMap detected no pocket in the helical segment. Eight experimental PTB-domain structures identified by sequence search (31-37% identity; Supplementary Table S1) superposed on the model with Cα RMSD 1.68-2.73 Å. Five are peptide complexes; the 24 model residues lying within 5 Å of the bound peptide in at least two of the five define a composite peptide-binding groove, of which 15 are contacted in all five.

**Figure 4.**
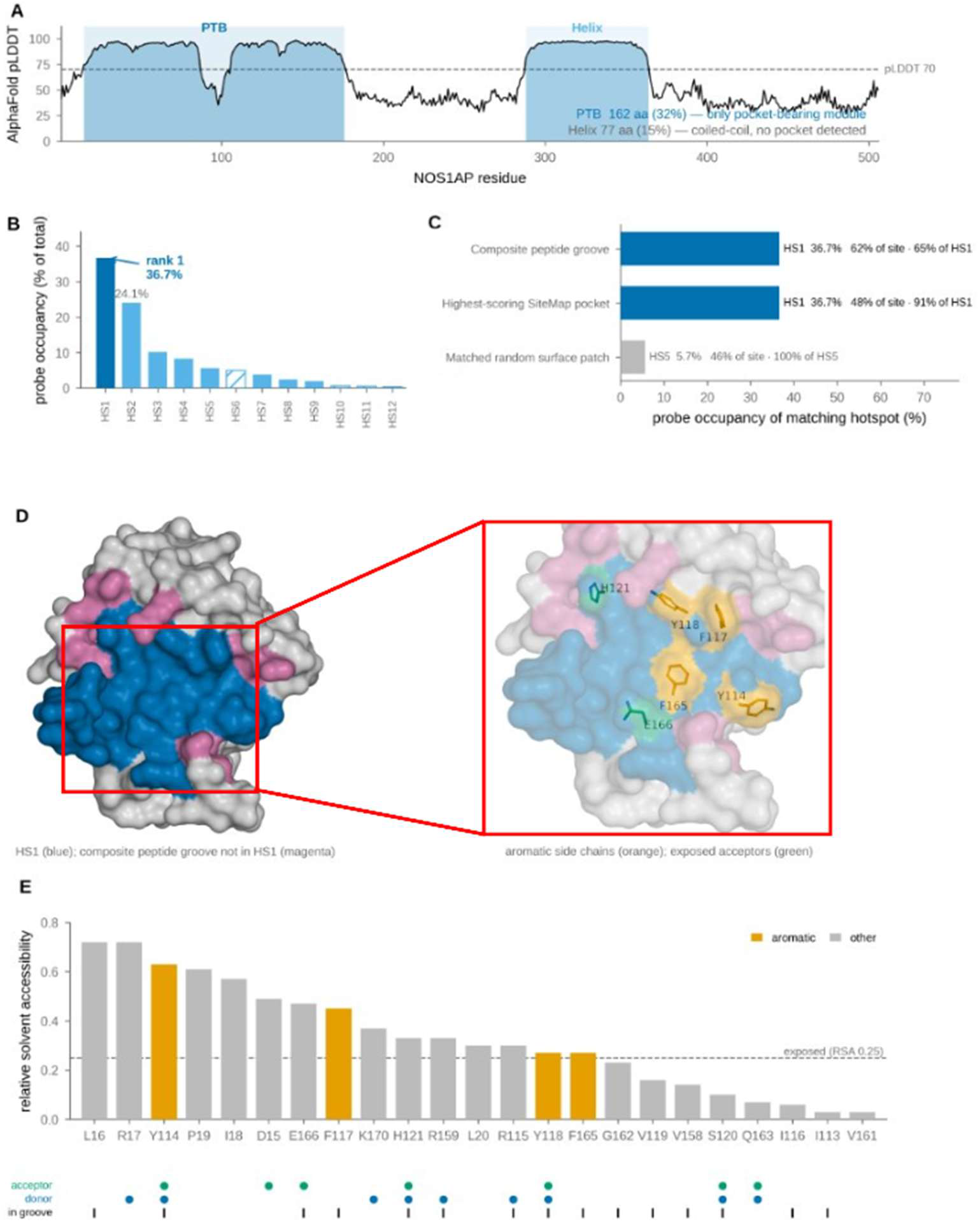
Mixed-solvent simulations nominate a candidate ligand-binding hotspot on the NOS1AP PTB domain. (**a**) Per-residue AlphaFold confidence across NOS1AP; shaded blocks mark the two confidently folded modules. Dashed line, pLDDT = 70. (**b**) Probe occupancy of the twelve hotspots identified by mixed-solvent molecular dynamics (three aromatic probes, ten simulations each; 600 ns simulated, 150 ns analyzed), as a percentage of total. Solid bars, hotspots populated by all three probes; hatched bars, by two. (**c**) Correspondence between independently defined sites and the hotspots; overlap is given in both directions. (**d**) Surface of the PTB domain with HS1 in blue and composite-groove residues not contained in HS1 in magenta; right, close-up of HS1 showing four aromatic side chains (orange) and exposed hydrogen-bond acceptors E166 and H121 (green). (**e**) Composition of HS1: relative solvent accessibility per residue, with side-chain hydrogen-bond acceptor, donor, and composite-groove membership marked below the axis.

We mapped the PTB domain with three aromatic cosolvent probes chosen to resemble the compound scaffold, isoquinoline, benzothiazole, and pyridine, in ten independent simulations per probe (600 ns simulated, of which the 150 ns of unrestrained production sampling was analyzed). The probes were selected from the chemical structures of the series; measured affinities were not used at any point. Probe occupancy was not uniformly distributed. The highest-ranked hotspot, HS1, accounted for 36.7% of total probe occupancy and the two highest-ranked hotspots together for 60.8%, with the remaining ten each below 11% (Figure 4b). The ranking is not driven by the chemistry of any single probe: analyzed separately, each of the three placed HS1 first and HS2 second, with HS1 leading by 1.1-to 2.4-fold (Supplementary Fig. 91).

HS1 overlaps the composite peptide-binding groove. The hotspot comprises 23 residues, D15-L20, I113-H121, and eight residues between V158 and K170, and contains 62% of the homolog-derived composite groove, while 91% of its own residues fall within the highest-scoring SiteMap pocket. A size-, exposure-, and confidence-matched random surface patch instead matched a minor hotspot accounting for 5.7% of occupancy (Figure 4c,d). Within HS1, four aromatic side chains (Y114, F117, Y118, F165) and seven potential hydrogen-bond acceptors are present; four of the acceptors, E166, Y118, Y114, and H121, are solvent-exposed (relative accessibility 0.27-0.63) and lie within the composite groove, separated by 8.8-26.6 Å and so distributed along the groove rather than clustered (Figure 4e).

The SAR of the series is compatible with binding in a confined groove of this kind, though the comparison requires care and we draw it narrowly. The series contains one rigorous single-point increment: deletion of the 4-methyl group adjacent to the inter-ring bond improves affinity 18-fold in the isoquinolinyl context (**31** versus **49**) while reducing it 5-fold in the thiophenyl context (**29** versus **44**). Steep, context-dependent sensitivity to a single methyl group at the junction between two aromatic systems is the behavior expected of a ligand bound in a narrow groove whose permitted inter-ring geometry is constrained by the surrounding surface, and is not easily reconciled with binding at a shallow or solvent-exposed site.

Nor is the affinity of the series explained by bulk physicochemical properties. Regression of p*K*_D_ against calculated logP and topological polar surface area across the 20 compounds with measured affinities accounts for 18% of the variance and does not reach significance (r² = 0.18, p = 0.064). NOSTRA is a pronounced outlier to even this weak trend, binding approximately 70-fold more tightly than its calculated properties predict (residual +1.85 log units, against +0.54 and +0.52 for the next largest). Potency in this series therefore reflects specific molecular recognition rather than lipophilic or polar drift.

The remaining comparisons in the series each involve two or more simultaneous structural changes, and the terminal-cap round is confounded with a change in the central ring, so no matched pair in this dataset isolates a hydrogen-bonding interaction. HS1 presents four solvent-exposed hydrogen-bond acceptors and five exposed donors distributed along the groove, and the series is consistent with occupancy of such a surface, but the SAR cannot currently assign any specific contact. Because the binding data were obtained on full-length NOS1AP and no direct measurement localizes the compounds to this region, we nominate HS1 as a candidate site rather than an established one. It provides the first structural localization hypothesis for this series, narrowing the search from a 506-residue protein to a 23-residue groove. We note that this analysis addresses the binary NOSTRA-NOS1AP interaction and does not define the ternary interface described below.

### A Selectivity Screen across Six NOS1AP Interactions Identifies SCRIB

Having established direct binding, we asked what NOSTRA does to NOS1AP function. Because NOS1AP acts through the complexes it assembles, we screened NOSTRA against six established NOS1AP interactions spanning distinct regions of the protein: SCRIB and RASD1/Dexras1 (PTB-domain interactors), SYN1 (synapsin), NOS1/nNOS (C-terminal PDZ-binding motif), carboxypeptidase E (central region), and MAPT/tau. Each interaction was assayed across a 0-10 µM NOSTRA concentration range in triplicate (Figure 5).

**Figure 5.**
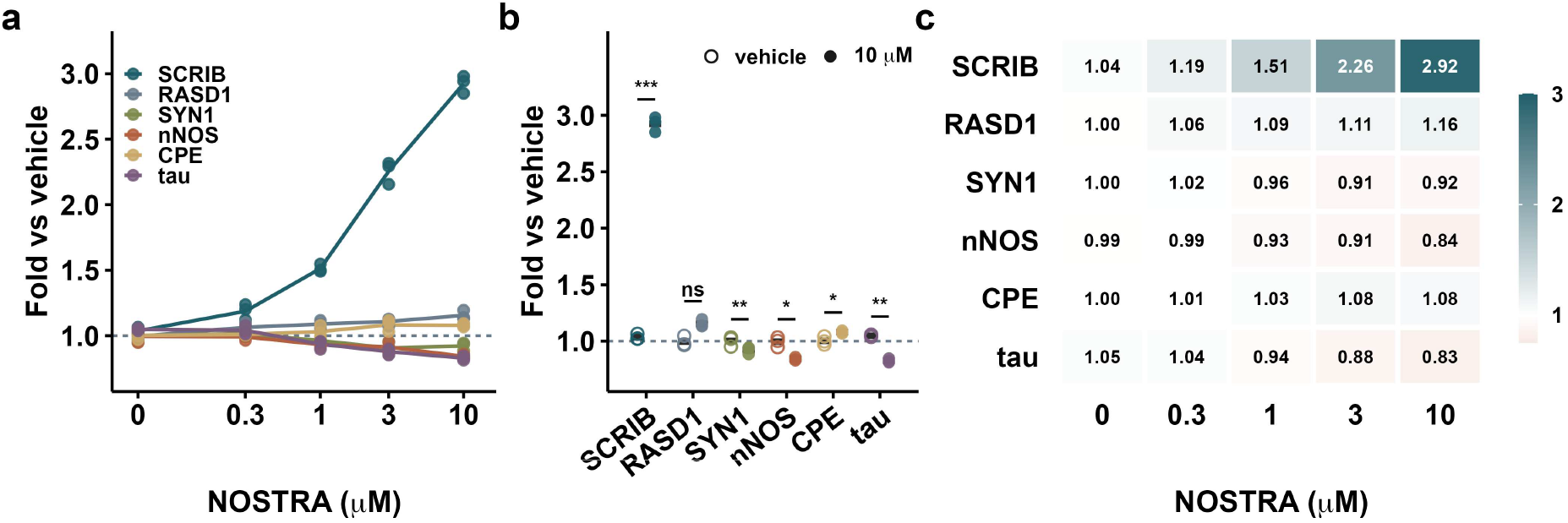
NOSTRA selectively enhances the NOS1AP–SCRIB interaction. (**a**) Concentration response for each of six NOS1AP interactions; all individual replicate values shown (n = 3 independent experiments), lines connect means. (**b**) Vehicle versus 10 µM NOSTRA for each partner; brackets connect the compared groups (paired t-test). (**c**) Heat map of mean fold change across the concentration series. ns, not significant; \**p < 0.05;* \*\**p < 0.01;* \*\*\**p* < 0.001.

NOSTRA did not inhibit any of the six interactions. Instead it produced a concentration-dependent enhancement of a single one: the NOS1AP-SCRIB association increased 1.19-, 1.51-, 2.26-, and 2.92-fold at 0.3, 1, 3, and 10 µM respectively (*p* = 0.0002 at 10 µM versus vehicle). No comparator showed a comparable effect. RASD1 was essentially unchanged (1.16-fold at 10 µM, *p* = 0.051), as were CPE (1.08-fold, *p* = 0.030) and SYN1 (0.92-fold, *p* = 0.0068). Two interactions decreased modestly: nNOS (0.84-fold, *p* = 0.012) and tau (0.83-fold, *p* = 0.0018). At 10 µM, SCRIB differed from every comparator with *p* < 1 × 10^-5^. The small decreases at nNOS and tau are consistent with a redistribution of NOS1AP toward the stabilized SCRIB complex rather than with direct inhibition at those sites, though their magnitude is at the limit of what this assay resolves and we do not interpret them further.

The partner selectivity of this effect is its most informative feature. RASD1/Dexras1 engages the same PTB domain as SCRIB yet was unaffected, so occupancy of the PTB module is not by itself sufficient to enhance an interaction. Whatever NOSTRA does requires complementarity between the compound, NOS1AP, and the specific partner, the defining property of a cooperative glue, and the feature that distinguishes such a compound from one that would merely increase the general avidity of the adaptor. It follows that the HS1 site nominated above, which describes the binary interaction, cannot by itself account for the selectivity we observe.

### NOSTRA Cooperatively Stabilizes the NOS1AP-SCRIB Complex

We next characterized the SCRIB effect using three orthogonal approaches (Figure 6). SPR measurement of the NOS1AP-SCRIB interaction in the presence of increasing NOSTRA showed progressive tightening of the complex (Figure 6 a-c). The binary interaction had *K*_D_ = 26.0 ± 6.0 µM. In the presence of 0.3, 1, 3, and 10 µM NOSTRA this fell to 20.7 ± 5.6, 8.9 ± 3.0, 2.4 ± 0.9, and 1.0 ± 0.7 µM (n = 6 independent experiments; Holm-adjusted *p* = 0.13, 0.0012, 0.0005, and 0.0005 versus vehicle), corresponding to cooperativity factors (α = *K*_D_,binary/ *K*_D_,ternary) of 1.30 ± 0.36, 3.13 ± 1.03, 11.75 ± 3.16, and 30.50 ± 11.14. Cooperativity factors were computed per experiment and then averaged; because a mean of ratios exceeds the ratio of means, α = 30.5 is larger than the 25-fold difference between the mean binary and mean ternary *K*_D_ values. Across this range the dissociation rate constant fell 14.2-fold (3.18 × 10^-1^ to 2.25 × 10^-2^ s^-1^, *p* = 0.0006), whereas the association rate constant increased 1.77-fold (1.22 × 10^4^ to 2.16 × 10^4^ M^-1^ s^-1^, *p* = 0.0041). The stabilization is therefore predominantly a retardation of complex dissociation, accompanied by a smaller acceleration of complex formation. This kinetic asymmetry, an order-of-magnitude effect on complex lifetime against a sub-two-fold effect on association, is the signature expected of a compound occupying an interface within the assembled complex rather than one pre-organizing either partner for binding.

**Figure 6.**
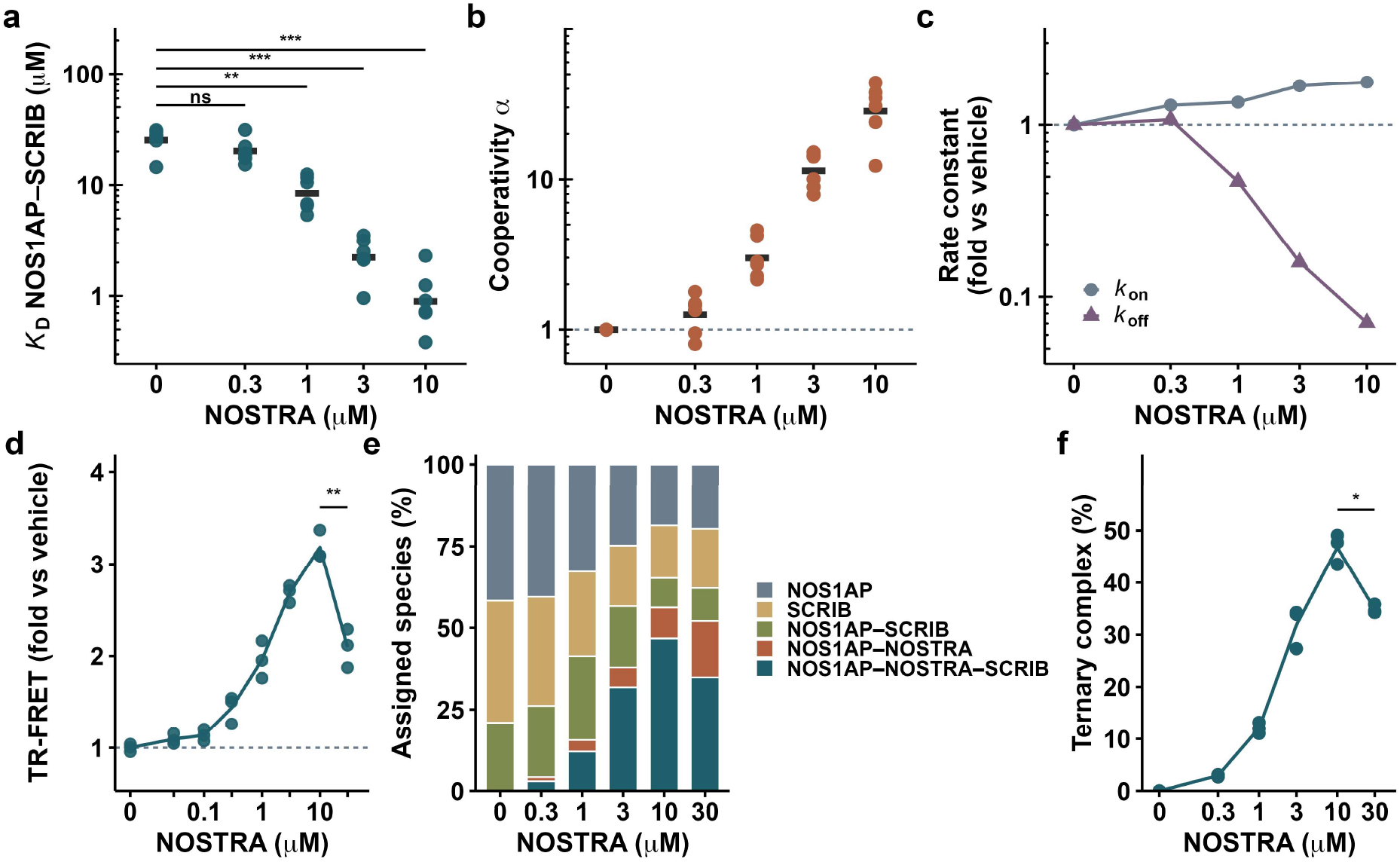
NOSTRA is a cooperative stabilizer of the NOS1AP-SCRIB complex. **(a)** SPR-derived *K*_D_ of the NOS1AP-SCRIB interaction as a function of NOSTRA concentration; all replicate values shown (n = 6 independent experiments), crossbars indicate means, brackets connect the compared groups and report Holm-adjusted paired t tests versus vehicle. **(b)** Cooperativity factor α = *K*_D_,binary/ *K*_D_,ternary (n = 6). **(c)** Kinetic decomposition: fold change in k_on_ and k_off_ relative to vehicle (n = 6). **(d)** TR-FRET proximity signal across 0-30 µM NOSTRA (n = 3); the bracket compares 30 µM with 10 µM and shows the hook effect. **(e)** Native mass spectrometry species distribution (n = 3). **(f)** Ternary complex abundance versus NOSTRA concentration (n = 3); the bracket compares 30 µM with 10 µM. ns, not significant; \**p* < 0.05; \*\**p* < 0.01; \*\*\**p* < 0.001.

A TR-FRET proximity assay (Figure 6d) reproduced the concentration dependence, with signal rising from 1.09-fold at 30 nM to a maximum of 3.18 ± 0.16-fold at 10 µM (*p* = 0.0018 versus vehicle). At 30 µM the signal declined to 2.09 ± 0.21-fold (*p* = 0.0040 versus 10 µM), the hook effect characteristic of a bridging mechanism, in which saturating compound populates binary complexes at the expense of the ternary species. Native mass spectrometry (Figure 6e,f) resolved a discrete NOS1AP:NOSTRA:SCRIB species and allowed the full equilibrium to be quantified. The ternary complex was undetectable in the absence of compound and rose to 2.9%, 12.0%, 31.8%, and 46.7% of assigned species at 0.3, 1, 3, and 10 µM NOSTRA. Over the same range the binary NOS1AP-SCRIB species and both free proteins declined, consistent with recruitment of both partners into the ternary assembly. At 30 µM the ternary species declined to 34.8% (*p* = 0.016 versus 10 µM) while the binary NOS1AP-NOSTRA species continued to rise, independently reproducing the hook effect observed by TR-FRET. Because native mass spectrometry measures the ternary species directly rather than inferring proximity, this observation is not confounded by the signal quenching that can mimic or mask a hook in energy-transfer formats. Therefore, these experiments establish that NOSTRA acts as a molecular glue: it binds NOS1AP directly, forms a physically detectable ternary complex with SCRIB, increases the affinity of that interaction by more than an order of magnitude with positive cooperativity, acts predominantly by slowing complex dissociation, and displays the biphasic concentration dependence diagnostic of a bridging mechanism.

### The Mechanism Operates on the Endogenous Complex in Human iPSC-Derived Cortical Neurons

To determine whether the mechanism established with purified proteins operates in a human neuronal context, we treated iPSC-derived cortical neurons from three independent healthy donors across a 0-10 µM NOSTRA range (Figure 7). Endogenous NOS1AP-SCRIB complex formation, measured by HTRF proximity assay (Figure 7a), increased in a concentration-dependent manner from 1.12-fold at 0.1 µM (*p* = 0.54) to 1.38-fold at 0.3 µM (*p* = 0.014), 1.67-fold at 1 µM (*p* = 0.009), 2.01-fold at 3 µM (*p* = 0.004), and 2.15-fold at 10 µM (*p* < 0.001). The magnitude and the concentration dependence closely track the biophysical measurements, indicating that the glue mechanism operates on the native complex at endogenous expression levels.

**Figure 7.**
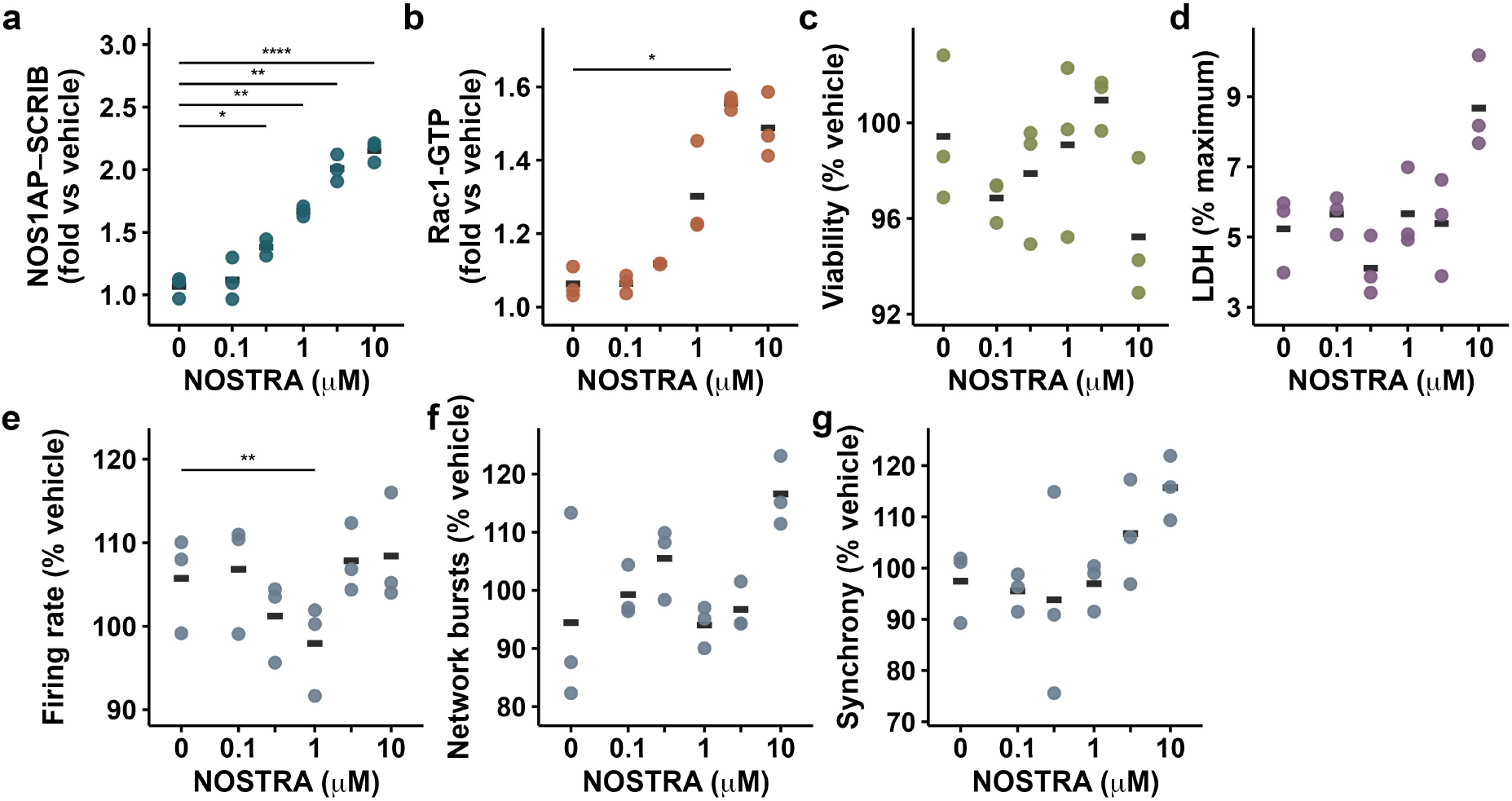
NOSTRA engages the endogenous NOS1AP-SCRIB complex in healthy human iPSC-derived cortical neurons. (**a**) NOS1AP-SCRIB HTRF proximity signal. (**b**) Rac1-GTP. (**c**) CellTiter-Glo viability. (**d**) LDH release. (**e-g**) MEA mean firing rate, network burst frequency, and synchrony, as percentage of vehicle. Each point is the mean of one donor (n = 3 donors, H1– H3); crossbars indicate group means. Brackets connect the compared groups; *p* values are Holm-adjusted within each panel. \**p < 0.05;* \*\**p < 0.01;* \*\*\**p* < 0.001; \*\*\*\**p* < 0.0001.

Consistent with the established role of SCRIB in coupling to the β-PIX/GIT1/PAK module, active Rac1-GTP increased over the same range, reaching 1.56-fold at 3 µM (*p* = 0.014). The value at 10 µM (1.49-fold) did not retain significance after correction (*p* = 0.10), and no change was detected below 1 µM (Figure 7b). The downstream readout therefore follows target engagement with the expected concentration offset, with the maximal effect at 3 µM. Neuronal viability was unaffected across the full range (Figure 7c). LDH release was likewise statistically unchanged (Figure 7d). Microelectrode-array recordings (Figure 7e-g) showed no deterioration of spontaneous network activity. Network burst frequency and synchrony were unchanged at every concentration (*p* ≥ 0.07), and all values remained within 94-117% of vehicle. These data define a working concentration window of approximately 1-3 µM in which target engagement and downstream signaling are robust, network activity is preserved, and no cytotoxicity is detectable. At 10 µM, engagement plateaus, and we regard that concentration as the upper bound of the usable range.

We deliberately do not frame the Rac1 increase as the therapeutic objective. Rac1 biology is strongly context-dependent, with excessive or mislocalized activation implicated in amyloid processing and tau pathology, and the effect observed here is modest, concentration-limited, and plateaus at the same point as target engagement. We interpret it as a pharmacodynamic marker confirming that the stabilized complex remains signaling-competent, not as the mechanism of any benefit. The biphasic profile is reassuring in this respect: a compound driving pathological Rac1 activation would be expected to produce worsening phenotypes at high concentration, whereas what we observe is a plateau in both engagement and effect.

### Activity in AD Patient-Derived Neurons

We then evaluated NOSTRA in iPSC-derived cortical neurons from three AD patients across a 0-10 µM concentration range (Figure 8). Every comparison reported below is against vehicle-treated neurons from the same donor, and the effects therefore describe modification of the AD phenotype rather than normalization toward a healthy reference. At baseline the AD neurons secreted Aβ42 and Aβ40 at 120 and 800 pg/mL (ratio 0.150), p-tau217 at 37.5 pg/mL, p-tau181 at 77.6 pg/mL, total tau at 507 pg/mL and neurofilament light at 140.1 pg/mL, with a mean firing rate of 1.57 Hz, network burst frequency of 0.73 min-1 and synchrony index of 0.34.

**Figure 8.**
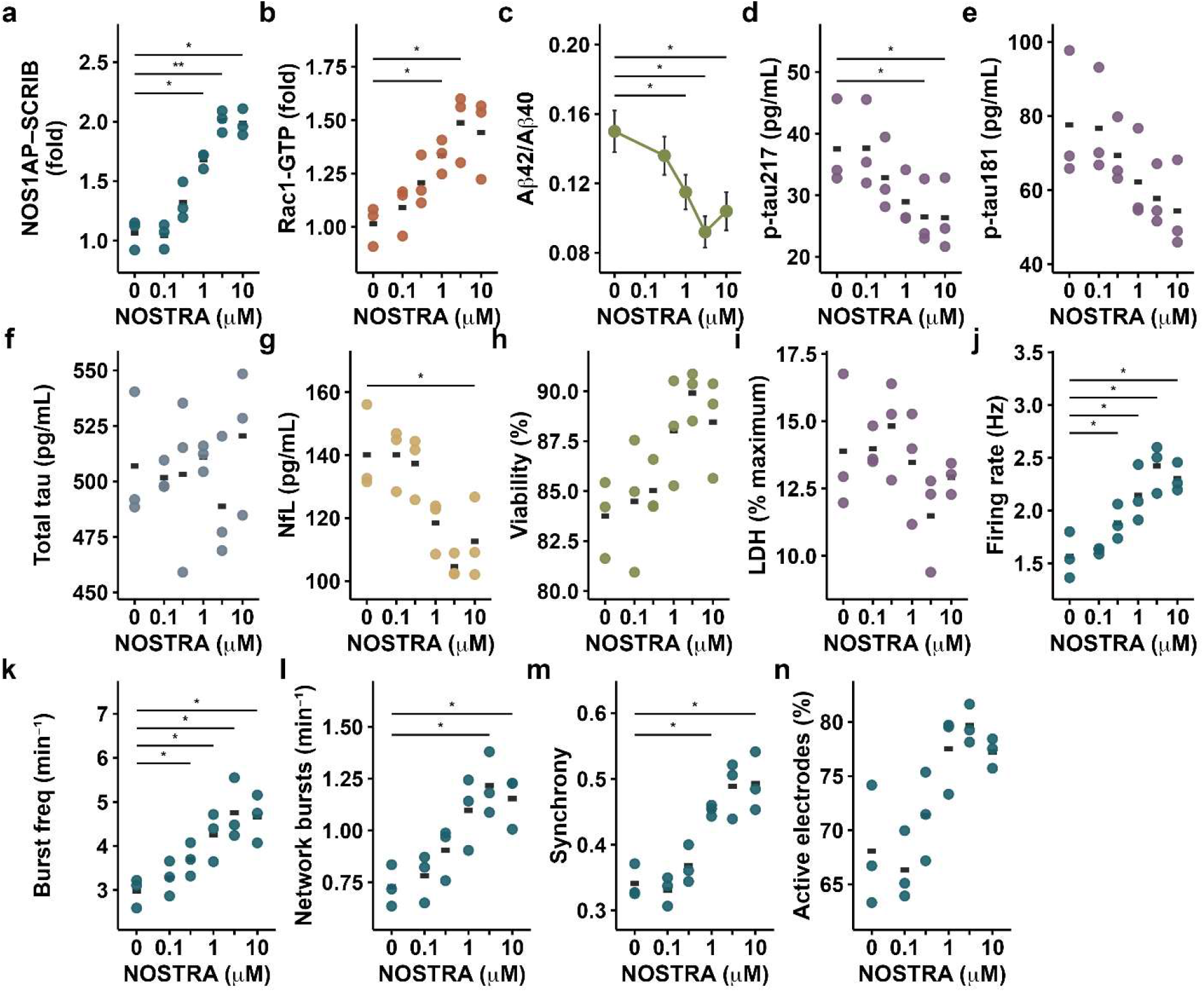
NOSTRA in AD-derived iPSC cortical neurons. **(a)** NOS1AP-SCRIB HTRF proximity. **(b)** Rac1-GTP. **(c)** Secreted Aβ42/Aβ40 ratio. **(d)** p-tau217. **(e)** p-tau181. **(f)** Total tau. **(g)** Neurofilament light. **(h)** CellTiter-Glo viability. **(i)** LDH release. **(j-n)** MEA mean firing rate, burst frequency, network burst frequency, synchrony, and proportion of active electrodes. Each point is the mean of one AD line (n = 3 lines) and crossbars indicate group means; brackets connect the compared groups and report Holm-adjusted paired t tests versus vehicle within each panel. Panel c shows the mean ± SD of the secreted Aβ42/Aβ40 ratio (n = 3 independent experiments per concentration), compared with vehicle by unpaired two-tailed t tests with Holm correction. All comparisons are against vehicle-treated neurons of the same donor. \**p* < 0.05; \*\**p* < 0.01.

NOSTRA increased NOS1AP-SCRIB proximity in AD neurons with the same concentration dependence seen in healthy neurons: 1.68-fold at 1 µM (*p* = 0.012), 2.01-fold at 3 µM (*p* = 0.007), and 1.99-fold at 10 µM (*p* = 0.029). Rac1-GTP rose correspondingly, reaching 1.33-fold at 1 µM and 1.49-fold at 3 µM, both significant after correction (*p* = 0.036); the value at 10 µM (1.44-fold) fell just short (*p* = 0.056). Disease state therefore does not impair target engagement, and the downstream signaling response tracks it with the same 1-3 µM optimum observed in healthy neurons.

Phosphorylated tau species decreased with treatment. p-tau217 fell from 37.5 pg/mL in vehicle to 28.9 at 1 µM, 26.5 at 3 µM, and 26.4 at 10 µM, a 29% reduction at the point of maximal effect; the 3 and 10 µM comparisons were significant after correction for multiple concentrations (*p* = 0.038 for both). p-tau181 declined in parallel, from 77.6 pg/mL to 57.7 at 3 µM and 54.4 at 10 µM, but did not survive correction (p = 0.12 at 10 µM). Total tau was unchanged at every concentration (p = 1.00), establishing that the phospho-tau changes reflect altered phosphorylation state rather than altered tau abundance. NOSTRA also lowered secreted Aβ42 in a concentration-dependent manner, from 120 ± 11 pg/mL in vehicle to 108 ± 11, 91 ± 9 and 72 ± 7 pg/mL at 0.3, 1 and 3 µM, a 40% reduction at 3 µM, while Aβ40 was essentially unchanged over the same range (800 ± 54 to 780 ± 46 pg/mL, a 2.5% decrease). The Aβ42/Aβ40 ratio consequently fell from 0.150 ± 0.012 to 0.092 ± 0.009 at 3 µM and returned partway toward baseline at 10 µM (0.104 ± 0.011). A selective reduction in Aβ42 without a corresponding change in Aβ40 is the profile associated with modulation rather than inhibition of γ-secretase processing. Amyloid values are reported as mean ± SD of three independent experiments; the reductions in the Aβ42/Aβ40 ratio at 1, 3 and 10 µM were significant after Holm correction (*p* = 0.036, 0.010 and 0.024), whereas 0.3 µM was not (*p* = 0.21). Both tau and amyloid endpoints respond over the same 1-3 µM window that produces target engagement, with the effect plateauing or reversing at 10 µM.

Neurofilament light release, a marker of axonal damage, fell from 140.1 pg/mL to 118.4 at 1 µM, 104.5 at 3 µM, and 112.6 at 10 µM. The reduction at 10 µM survived correction (*p* = 0.043); the 3 µM value, the largest numerical effect at 25%, did not (p = 0.10). Viability improved modestly (83.8% to 89.9% at 3 µM) and LDH release decreased (13.9% to 11.5%), neither reaching significance. Mean firing rate improved significantly at every concentration from 0.3 to 10 µM (*p* = 0.013-0.031), rising from 1.57 Hz in vehicle to 2.42 Hz at 3 µM; burst frequency likewise improved across 0.3–10 µM (2.97 to 4.76 min⁻¹, *p* = 0.031–0.049); network burst frequency improved at 3 µM (0.73 to 1.22 min⁻¹, *p* = 0.018) and 10 µM (*p* = 0.041); and synchrony improved at 1 µM (*p* = 0.044) and 10 µM (0.34 to 0.49, *p* = 0.038). The proportion of active electrodes rose from 68.1% to 79.7% at 3 µM but did not reach significance (*p* = 0.47).

The concordance across the different readouts is the central observation. Target engagement, downstream Rac1 signaling, amyloid and tau pathology, axonal injury, and network function all respond over the same 1-3 µM concentration range and all plateau or decline at 10 µM, the behavior expected of a cooperative glue operating through a defined ternary complex and difficult to account for by a non-specific mechanism. Among the molecular-pathology endpoints, p-tau217 reaches significance after correction, while p-tau181 moves in the same direction without doing so at the present sample size. The improvements are consistent with reinforcing a scaffolding complex that supports network integrity under disease-associated stress. This also means the therapeutic hypothesis does not require NOS1AP-SCRIB disassembly to be part of AD pathogenesis, a claim for which there is currently no evidence.

### In Vitro ADME Profile

We profiled NOSTRA in a standard in vitro panel to establish whether the chemotype is suitable for further development toward a central nervous system (CNS) indication (Table 2). NOSTRA showed satisfactory aqueous solubility (kinetic 78 µM, thermodynamic 53 µM at pH 7.4), moderate lipophilicity (logD_7.4_ = 2.21), and weak basicity (p*K*_a_ 7.2), properties within the ranges typically associated with CNS-penetrant compounds. Metabolic stability was acceptable in both species: human liver microsomal t₁/₂ 74 min with intrinsic clearance of 8.2 µL/min/10^6^ hepatocytes, and mouse t₁/₂ 50 min with Cl_int_ 14.1 µL/min/10^6^. The compound was stable in human plasma (95% remaining at 120 min), showed no marked red-cell partitioning (blood:plasma 0.90), and had moderately high plasma protein binding (*f*_u,p_ 0.078 human, 0.067 mouse) with a measurable unbound brain fraction (*f*_u,brain_ 0.037). Permeability was high and efflux liability low: PAMPA-BBB *P*_e_ = 8.7 × 10^-6^ cm/s, MDCK-MDR1 A→B *P*_app_ = 16.2 × 10^-6^ cm/s with an efflux ratio of 1.36, and a BCRP efflux ratio of 1.21.

**Table 2.** In vitro ADME and safety profile of NOSTRA.

| Parameter | Value |
| --- | --- |
| Kinetic solubility, pH 7.4 | 78 $\mu\text{M}$ |
| Thermodynamic solubility, pH 7.4 | 53 $\mu\text{M}$ |
| $\log D_{7.4}$ | 2.21 |
| $pK_a$ | 7.2 |
| **Human liver microsome $t_{1/2}$** | 74 min |
| **Mouse liver microsome $t_{1/2}$** | 50 min |
| **Human hepatocyte $Cl_{int}$** | 8.2 $\mu\text{L}/\text{min}/10^6$ cells |
| **Mouse hepatocyte $Cl_{int}$** | 14.1 $\mu\text{L}/\text{min}/10^6$ cells |
| **Human plasma, 120 min** | 95% remaining |
| **Human $f_{u,p}$** | 0.078 |
| **Mouse $f_{u,p}$** | 0.067 |
| **$f_{u,brain}$** | 0.037 |
| **Blood:plasma ratio** | 0.90 |
| **PAMPA-BBB $P_e$** | $8.7 \times 10^{-6}$ cm/s |
| **MDCK-MDR1 A→B $P_{app}$** | $16.2 \times 10^{-6}$ cm/s |
| **MDCK-MDR1 efflux ratio** | 1.36 |
| **BCRP efflux ratio** | 1.21 |
| **CYP1A2 / 2C9 / 2C19 / 2D6**<br>**/ 3A4 $IC_{50}$** | 29 / 19 / 24 / 36 / 31 $\mu\text{M}$ |
| **CYP3A4 time-dependent**<br>**inhibition** | 1.10-fold shift |
| **hERG $IC_{50}$** | 41 $\mu\text{M}$ |
| **HepG2 $CC_{50}$** | 98 $\mu\text{M}$ |
| **Primary hepatocyte $CC_{50}$** | 80 $\mu\text{M}$ |
| **Glutathione trapping** | Negative |

Values are means of n=3.

The safety and drug-drug interaction profile was clean at the concentrations relevant to the cellular activity described above. Inhibition of the five major cytochrome P450 isoforms was weak (IC_50_ = 19-36 µM), with no time-dependent inhibition of CYP3A4 (1.10-fold shift). hERG inhibition was weak (IC_50_ = 41 µM), giving a margin of more than 40-fold over the 1 µM working concentration, and cytotoxicity was low in both HepG2 cells (CC_50_ = 98 µM) and primary hepatocytes (CC_50_ = 80 µM). Glutathione trapping was negative, indicating no detectable reactive metabolite formation. This profile indicates that NOSTRA is a suitable chemical probe for cellular and ex vivo work and a promising starting point for optimization toward in vivo studies.

## CONCLUSIONS

Adaptor proteins have been approached almost exclusively through interaction disruption, and our own prior NOS1AP programs followed that convention. The work reported here establishes that the opposite pharmacology is achievable against the same protein. NOSTRA satisfies the full evidentiary standard for a molecular glue: direct binding to one partner at submicromolar affinity by two orthogonal methods, concentration-dependent enhancement of a specific interaction, a cooperativity factor of 30.5, a physically resolved ternary complex by native mass spectrometry, and a hook effect reproduced in two independent assay formats. To our knowledge it is the first small molecule reported to stabilize any NOS1AP complex. In human iPSC-derived cortical neurons it engages the endogenous complex and, in neurons from AD patients, improves microelectrode-array network activity and reduces neurofilament light release, within a concentration window in which viability is preserved.

NOSTRA provides the first chemical tool for interrogating NOS1AP-SCRIB stabilization and establishes interaction stabilization as a viable pharmacology for this adaptor. More broadly, the architecture of NOS1AP, several mutually exclusive partners engaging distinct structural regions, is a configuration shared by many scaffolding proteins, and one for which partner-selective stabilization offers something that inhibition structurally cannot: the ability to bias a protein toward one of its functions rather than removing it from all of them. We expect adaptor and scaffolding proteins to prove a productive, and currently underexploited, substrate for stabilizer pharmacology.

## Experimental section

### DNA-encoded library selection

A 1.4-billion-member DNA-encoded chemical library (AlphaMa SDEL Pubkit) was obtained from AlphaMa Biotechnology (Suzhou, China). According to the manufacturer, each kit supports a single selection condition, using either the protein target or blank beads for a no-target control (NTC). For the NOS1AP selection, 10 μg of target protein was captured on 20 μL of HisPur™ Ni-NTA magnetic beads and exposed to the DNA-encoded library. Three rounds of selection were conducted for each condition. Each round comprised protein immobilization, library panning, washing, and heat-mediated elution, following the manufacturer’s protocol. Eluates collected after the third round were analyzed by next-generation sequencing to identify library members retained in each condition. FASTQ files were processed using AlphaMa’s proprietary decoding software, which is not publicly disclosed, to assign chemical identities and estimate their copy numbers in each selection. Library members and their associated building-block combinations were evaluated by comparing their representation in the NOS1AP selection with that in the matched NTC. Candidates showing enrichment in the presence of NOS1AP and interpretable relationships between structure and enrichment were selected for off-DNA synthesis and subsequent biophysical testing.

### Chemistry

All reagents, solvents, and other chemicals were purchased from commercial suppliers and used as received unless otherwise specified. ^1^H NMR spectra were recorded on a Bruker 500 MHz spectrometer. Chemical shifts (δ) are reported in parts per million (ppm) relative to the residual solvent signal. Signal multiplicities are denoted as s (singlet), d (doublet), dd (doublet of doublets), t (triplet), q (quartet), td (triplet of doublets), dt (doublet of triplets), ddd (doublet of doublet of doublets), br (broad), and m (multiplet).

Crude products were purified by flash column chromatography using a CombiFlash system fitted with prepacked RediSep RF silica cartridges. Final pyrazole-containing compounds were purified by preparative HPLC employing a gradient of 5-95% acetonitrile in water containing 0.01% formic acid as the mobile phase. Reaction progress was monitored by thin-layer chromatography (TLC) on silica gel 60 F_254_ plates. Product purity was analyzed with HPLC using the above mentioned method. All compounds are >95% pure by HPLC analysis.

### Synthetic Procedures and Characterization of the Compounds

#### General Procedure for intermediates 3 and 61-67

To a solution of 4-fluoro-*N*-methyl-3-nitrobenzamide (**1**, 5.05 mmol, 1g) in water (20 mL) was added potassium carbonate (10.09 mmol), followed by addition of amine **2/54-60** (5.05 mmol). The reaction mixture was stirred at 80 °C for 3-4 h, and the progress of the reaction was monitored by TLC. After completion, the reaction mixture was cooled to room temperature, and the resulting yellow precipitate was collected by filtration. The solid was washed with water followed by a small amount of acetone and dried under reduced pressure to afford intermediates in quantitative yields.

#### General Procedure for intermediate 4

Intermediate **3** (500 mg, 1.72 mmol) was dissolved in ethanol (100 mL), and the reaction mixture was cooled in an ice bath. Aqueous HCl (1 N, 5 mL) was added, followed by the portionwise addition of tin(II) chloride (SnCl₂, 10.32 mmol) to the stirred solution. The reaction mixture was allowed to cool between additions to ensure complete dissolution of tin(II) chloride. After completion of the addition, the reaction was stirred at room temperature for 1.5 h. The reaction mixture was then basified to pH 10–12 by the addition of 10% aqueous NaOH and diluted with ethyl acetate (EtOAc). The resulting suspension was filtered through Celite™, and the layers were separated. The organic layer was washed with water and brine, dried over anhydrous Na₂SO₄, filtered, and concentrated under reduced pressure to afford the crude product, which was used directly in the subsequent step without further purification.

#### General Procedure for intermediate 6, 12, 39-43, 76-82

Intermediate **4/11/68-74** (200 mg, 0.76 mmol) was dissolved in ethanol, and aldehyde (**5/34-38/75**, 0.76 mmol) was added followed by addition of 0.76 mmol of sodium metabisulfite. The reaction mixture was stirred at 80 °C for 5h. Upon completion (monitored by TLC), the reaction mixture was filtered, and solvent was evaporated and triturated using diethyl ether and methanol to get desired compounds in quantitative yield. In case of compounds **76-82**, semi-prep HPLC was used to purify compounds using ACN:H_2_O with 0.1% formic acid as mobile phase.

#### General Procedure for intermediate 9

Under a nitrogen atmosphere, 6-bromoimidazo[1,5-*a*]pyridine (**7**, 500 mg, 2.53 mmol), bis(pinacolato)diboron (**8**, 5.07 mmol), potassium acetate (KOAc, 7.61 mmol), and 1,4-dioxane (25 mL) were added sequentially. The reaction mixture was purged with nitrogen for 15 min, followed by the addition of Pd(dtbpf)Cl₂ (10 mol%). The reaction mixture was heated to reflux for 3 h. After cooling to room temperature, the reaction mixture was filtered through Celite™, and the solvent was removed under reduced pressure. The crude product was purified by flash column chromatography using a CombiFlash system with hexane containing 0.1% triethylamine (TEA) as the eluent to afford the desired compounds in 40-50% yield.

#### General Procedure for 10, 13, 24-33, 44-53, 75

Intermediate **5**/**6/12/39-43** (100 mg, 0.227 mmol) was subjected to Suzuki–Miyaura cross-coupling with imidazo[1,5-*a*]pyridin-6-ylboronic acid (**9/5/14-23**, 0.22 mmol) in 1,4-dioxane (25 mL). Aqueous potassium carbonate solution (K₂CO₃, 0.22 mmol in 1.5 mL water) was added, and the reaction mixture was purged with nitrogen. Pd(dtbpf)Cl₂ (10 mol%) was added, and the reaction mixture was stirred at 90 °C for 3 h. Upon completion (as monitored by TLC), the reaction mixture was filtered through Celite™, and the filtrate was concentrated under reduced pressure. The crude product was purified by flash column chromatography using a Combi Flash system with dichloromethane/methanol as the eluent to afford the pure target compounds.

#### General Procedure for 11 and 68-74

Intermediate **3/61-67** (500 mg, 0.05 mol) was dissolved in ethanol (100 mL) in a three-necked flask. The reaction mixture was degassed by alternating vacuum and nitrogen for 15 min. Palladium on carbon (10% Pd/C, 125 mg) was added, followed by the dropwise addition of hydrazine monohydrate (N₂H₄·H₂O, 3 mL). The reaction mixture was stirred at 50 °C for 30 min, during which the yellow color gradually disappeared to afford a colorless solution. Upon completion of the reaction (as monitored by TLC), the mixture was filtered through Celite™ to remove the catalyst. The filtrate was concentrated under reduced pressure, and the resulting residue was washed with diethyl ether, followed by a small amount of methanol, to afford the desired products as a solid in 55-80% yield.

#### 2-(5-(imidazo[1,5-a]pyridin-6-yl)-4-methylpyridin-3-yl)-N-methyl-1-((5-methylisoxazol-3-yl)methyl)-1H-benzo[d]imidazole-5-carboxamide (10)

Light yellow solid; yield 54%; ^1^H NMR (500 MHz, DMSO) δ 8.65 (d, *J* = 7.6 Hz, 2H), 8.55 – 8.40 (m, 3H), 8.25 (d, *J* = 3.7 Hz, 1H), 7.85 (d, *J* = 7.5 Hz, 1H), 7.68 (dd, *J* = 9.0, 4.7 Hz, 2H), 7.44 (d, *J* = 3.5 Hz, 1H), 6.91 (d, *J* = 9.3 Hz, 1H), 6.04 (d, *J* = 3.8 Hz, 1H), 5.46 (d, *J* = 4.4 Hz, 2H), 2.82 (s, 3H), 2.31 (s, 3H), 2.12 (s, 3H); ^13^C NMR (126 MHz, DMSO) δ 171.12, 167.37, 160.18, 151.89, 151.25, 149.90, 146.17, 142.67, 137.18, 134.55, 129.72, 129.43, 129.24, 126.90, 123.21, 122.98, 122.23, 121.99, 119.95, 118.91, 118.04, 111.22, 101.38, 26.87, 17.68, 12.21; HRMS: calcd. for C_27_H_23_N_7_O_2_ [M+H] m/z 478.1991, found 478.1982.

#### 2-(5-(imidazo[1,5-a]pyridin-6-yl)-4-methylpyridin-3-yl)-N-methyl-1-((5-methyl-1H-pyrazol-3-yl)methyl)-1H-benzo[d]imidazole-5-carboxamide (13)

Light brown solid; yield 58%; ^1^H NMR (500 MHz, DMSO) δ 12.38 (s, 1H), 8.67 (d, *J* = 17.2 Hz, 2H), 8.52 (s, 1H), 8.43 (d, *J* = 5.0 Hz, 2H), 8.22 (s, 1H), 7.82 (d, *J* = 8.5 Hz, 1H), 7.68 (t, *J* = 9.6 Hz, 2H), 7.44 (s, 1H), 6.91 (d, *J* = 9.3 Hz, 1H), 5.74 (s, 1H), 5.25 (s, 2H), 2.82 (d, *J* = 4.6 Hz, 3H), 2.10 (d, *J* = 12.0 Hz, 6H); ^13^C NMR (126 MHz, DMSO) δ 167.43, 151.87, 151.04, 150.03, 147.46, 146.17, 142.70, 137.27, 134.46, 129.39, 129.36, 129.24, 127.40, 123.20, 122.59, 122.32, 122.04, 119.96, 118.72, 118.00, 111.48, 103.03, 26.85, 17.64; HRMS: calcd. for C_27_H_25_N_8_O [M+H] m/z 477.2151, found 477.2142.

#### 2-(5-(furan-2-yl)-4-methylpyridin-3-yl)-N-methyl-1-((5-methyl-1H-pyrazol-3-yl)methyl)-1H-benzo[d]imidazole-5-carboxamide (24)

white solid; yield 60%; ^1^H NMR (500 MHz, DMSO) δ 12.35 (s, 1H), 8.95 (s, 1H), 8.58 (s, 1H), 8.43 (q, J = 4.5 Hz, 1H), 8.22 (d, J = 1.5 Hz, 1H), 7.93 (d, J = 1.8 Hz, 1H), 7.82 (dd, J = 8.5, 1.6 Hz, 1H), 7.70 (d, J = 8.5 Hz, 1H), 6.98 (d, J = 3.4 Hz, 1H), 6.72 (dd, J = 3.4, 1.8 Hz, 1H), 5.70 (s, 1H), 5.20 (s, 2H), 2.82 (d, J = 4.4 Hz, 3H), 2.24 (s, 3H), 2.09 (s, 3H); ^13^C NMR (126 MHz, DMSO) δ 166.34, 150.72, 148.81, 148.19, 147.96, 143.42, 142.67, 141.57, 136.17, 128.29, 125.90, 121.52, 117.63, 111.43, 110.83, 110.40, 101.86, 25.76, 17.19; HRMS: calcd. for C_24_H_23_N_6_O_2_ [M+H] m/z 427.1882, found 427.1873.

#### N-methyl-1-((5-methyl-1H-pyrazol-3-yl)methyl)-2-(4-methyl-[3,4’-bipyridin]-5-yl)-1H-benzo[d]imidazole-5-carboxamide (25)

white solid; yield 59%; ^1^H NMR (500 MHz, DMSO) δ 12.39 (s, 1H), 8.79 – 8.67 (m, 3H), 8.60 (s, 1H), 8.43 (q, *J* = 4.5 Hz, 1H), 8.22 (d, *J* = 1.6 Hz, 1H), 7.82 (dd, *J* = 8.5, 1.6 Hz, 1H), 7.70 (d, *J* = 8.5 Hz, 1H), 7.59 – 7.52 (m, 2H), 5.74 (s, 1H), 5.25 (s, 2H), 2.82 (d, *J* = 4.4 Hz, 3H), 2.11 (s, 3H), 2.04 (s, 3H); ^13^C NMR (126 MHz, DMSO) δ 166.32, 162.56, 150.64, 149.59, 149.23, 144.16, 144.11, 141.58, 136.17, 134.48, 128.28, 126.37, 123.85, 121.52, 117.63, 110.38, 101.98, 25.76, 16.43; HRMS: calcd. for C_25_H_24_N_7_O [M+H] m/z 438.2042, found 438.2054.

#### 2-(5-(2-hydroxyphenyl)-4-methylpyridin-3-yl)-N-methyl-1-((5-methyl-1H-pyrazol-3-yl)methyl)-1H-benzo[d]imidazole-5-carboxamide (26)

white solid; yield 62%; ^1^H NMR (500 MHz, DMSO) δ 12.38 (s, 1H), 9.71 – 9.60 (m, 1H), 8.61 (t, *J* = 2.4 Hz, 1H), 8.44 (dt, *J* = 19.0, 3.5 Hz, 2H), 8.21 (s, 1H), 7.80 (d, *J* = 7.9 Hz, 1H), 7.69 – 7.60 (m, 1H), 7.27 (q, *J* = 4.8 Hz, 1H), 7.24 – 7.15 (m, 1H), 7.03 – 6.97 (m, 1H), 6.94 (dt, *J* = 8.9, 4.2 Hz, 1H), 5.71 (s, 1H), 5.21 (s, 2H), 2.81 (t, *J* = 4.5 Hz, 3H), 2.11 (s, 3H), 1.96 (d, *J* = 3.1 Hz, 3H); ^13^C NMR (126 MHz, DMSO) δ 167.45, 155.25, 152.22, 151.51, 149.13, 147.40, 146.40, 142.71, 139.73, 137.20, 135.71, 131.39, 130.07, 129.30, 126.67, 124.50, 122.49, 119.71, 118.66, 116.15, 111.50, 102.97, 26.85, 17.38; HRMS: calcd. for C_26_H_25_N_6_O_2_ [M+H] m/z 453.2039, found 453.2055.

#### 2-(5-(4-amino-2-methylphenyl)-4-methylpyridin-3-yl)-N-methyl-1-((5-methyl-1H-pyrazol-3-yl)methyl)-1H-benzo[d]imidazole-5-carboxamide (27)

white solid; yield 59%; ^1^H NMR (500 MHz, DMSO) δ 12.34 (s, 1H), 8.58 (s, 1H), 8.42 (d, *J* = 4.6 Hz, 1H), 8.37 (s, 1H), 8.20 (d, *J* = 1.4 Hz, 1H), 7.81 (dd, *J* = 8.5, 1.6 Hz, 1H), 7.69 (d, *J* = 8.5 Hz, 1H), 6.82 (d, *J* = 8.1 Hz, 1H), 6.57 – 6.45 (m, 2H), 5.68 (s, 1H), 5.21 (s, 2H), 5.13 (s, 2H), 2.81 (d, *J* = 4.4 Hz, 3H), 2.10 (s, 3H), 1.92 (s, 3H), 1.82 (s, 3H); ^13^C NMR (126 MHz, DMSO) δ 166.35, 151.13, 150.54, 147.97, 147.60, 145.18, 141.56, 137.29, 136.19, 135.36, 129.51, 128.18, 125.79, 123.22, 121.40, 117.55, 114.54, 110.97, 110.28, 101.74, 25.75, 19.09, 16.09; HRMS: calcd. for C_27_H_28_N_7_O [M+H] m/z 466.2364, found 466.2355.

#### N-methyl-1-((5-methyl-1H-pyrazol-3-yl)methyl)-2-(4-methyl-5-(1-methyl-1H-pyrrol-2-yl)pyridin-3-yl)-1H-benzo[d]imidazole-5-carboxamide (28)

white solid; yield 61%; ^1^H NMR (500 MHz, DMSO) δ 12.35 (s, 1H), 8.66 (s, 1H), 8.55 (s, 1H), 8.43 (d, *J* = 4.7 Hz, 1H), 8.21 (s, 1H), 7.82 (dd, *J* = 8.6, 1.6 Hz, 1H), 7.71 (d, *J* = 8.5 Hz, 1H), 6.93 (t, *J* = 2.3 Hz, 1H), 6.15 (d, *J* = 2.2 Hz, 2H), 5.72 (s, 1H), 5.23 (s, 2H), 3.44 (s, 3H), 2.82 (d, *J* = 4.4 Hz, 3H), 2.10 (s, 3H), 1.94 (s, 3H); ^13^C NMR (126 MHz, DMSO) δ 166.34, 151.12, 150.78, 148.52, 146.61, 141.56, 136.19, 128.89, 128.24, 127.31, 126.12, 122.95, 121.48, 117.59, 110.32, 109.38, 106.84, 101.85, 33.34, 25.76, 16.42; HRMS: calcd. for C_25_H_26_N_7_O [M+H] m/z 440.2199, found 440.2217.

#### N-methyl-1-((5-methyl-1H-pyrazol-3-yl)methyl)-2-(4-methyl-5-(5-phenylthiophen-2-yl)pyridin-3-yl)-1H-benzo[d]imidazole-5-carboxamide (29)

Light brown solid; yield 69%; ^1^H NMR (500 MHz, DMSO) δ 12.45 (s, 1H), 8.78 (s, 1H), 8.65 (s, 1H), 8.44 (d, *J* = 4.6 Hz, 1H), 8.23 (s, 1H), 8.17 (s, 1H), 7.83 (d, *J* = 8.5 Hz, 1H), 7.72 (dd, *J* = 13.1, 8.1 Hz, 3H), 7.64 (d, *J* = 3.7 Hz, 1H), 7.46 (t, *J* = 7.6 Hz, 2H), 7.41 (d, *J* = 3.8 Hz, 1H), 7.37 (d, *J* = 7.4 Hz, 1H), 5.73 (s, 1H), 5.25 (s, 2H), 2.82 (d, *J* = 4.3 Hz, 3H), 2.23 (s, 3H), 2.11 (s, 3H); ^13^C NMR (126 MHz, DMSO) δ 166.34, 150.71, 149.95, 148.67, 144.50, 144.15, 141.58, 136.20, 136.10, 132.65, 129.48, 128.64, 128.29, 127.45, 124.83, 123.81, 121.54, 117.64, 110.39, 101.98, 25.77, 17.06; HRMS: calcd. For C_30_H_27_N_6_OS [M+H] m/z 519.1967, found 519.1972.

#### N-methyl-1-((5-methyl-1H-pyrazol-3-yl)methyl)-2-(4-methyl-5-(4-morpholinophenyl)pyridin-3-yl)-1H-benzo[d]imidazole-5-carboxamide (30)

White solid; yield 72%; ^1^H NMR (500 MHz, DMSO) δ 12.37 (s, 1H), 8.58 (d, *J* = 3.3 Hz, 1H), 8.52 (t, *J* = 2.3 Hz, 1H), 8.42 (t, *J* = 4.4 Hz, 1H), 8.21 (s, 1H), 7.81 (d, *J* = 8.6 Hz, 1H), 7.68 (dd, *J* = 9.0, 3.0 Hz, 1H), 7.40 – 7.23 (m, 2H), 7.07 (dt, *J* = 9.0, 2.1 Hz, 2H), 5.72 (s, 1H), 5.22 (s, 2H), 3.76 (q, *J* = 4.3 Hz, 4H), 3.19 (q, *J* = 4.2 Hz, 4H), 2.82 (d, *J* = 4.2 Hz, 3H), 2.07 (d, *J* = 37.1 Hz, 6H); ^13^C NMR (126 MHz, DMSO) δ 167.44, 152.24, 151.05, 148.73, 145.00, 142.68, 139.68, 137.87, 137.24, 130.64, 129.29, 127.67, 127.25, 122.51, 118.66, 115.25, 111.44, 102.98, 66.58, 26.85, 17.74; HRMS: calcd. for C_30_H_32_N_7_O_2_ [M+H] m/z 522.2617, found 522.2620.

#### 2-(5-(isoquinolin-5-yl)-4-methylpyridin-3-yl)-N-methyl-1-((5-methyl-1H-pyrazol-3-yl)methyl)-1H-benzo[d]imidazole-5-carboxamide (31)

white solid; yield 52%; ^1^H NMR (500 MHz, DMSO) δ 12.38 (s, 1H), 9.45 (s, 1H), 8.81 (s, 1H), 8.64 – 8.49 (m, 2H), 8.43 (d, *J* = 4.6 Hz, 1H), 8.33 – 8.18 (m, 2H), 7.90 – 7.78 (m, 3H), 7.74 (d, *J* = 8.5 Hz, 1H), 7.23 (d, *J* = 5.9 Hz, 1H), 5.78 (s, 1H), 5.33 (d, *J* = 9.5 Hz, 2H), 2.82 (d, *J* = 4.4 Hz, 3H), 2.11 (s, 3H), 1.76 (s, 3H); HRMS: calcd. for C_29_H_26_N_7_O [M+H] m/z 488.2199, found 488.2188.

#### 2-(5-(1H-indol-7-yl)-4-methylpyridin-3-yl)-N-methyl-1-((5-methyl-1H-pyrazol-3-yl)methyl)-1H-benzo[d]imidazole-5-carboxamide (32)

Light brown solid; yield 45%; ^1^H NMR (500 MHz, CDCl_3_) δ 10.03 (s, 1H), 8.56 (d, *J* = 19.3 Hz, 2H), 8.04 (s, 1H), 7.76 (dd, *J* = 35.6, 8.4 Hz, 2H), 7.47 (d, *J* = 8.3 Hz, 1H), 7.20 (t, *J* = 7.8 Hz, 1H), 7.04 (d, *J* = 7.2 Hz, 1H), 6.81 (s, 1H), 6.61 (s, 1H), 5.81 (s, 1H), 5.32 – 5.21 (m, 2H), 3.01 (t, *J* = 3.7 Hz, 3H), 2.10 (d, *J* = 48.0 Hz, 6H); ^13^C NMR (126 MHz, CDCl_3_) δ 168.48, 136.42, 134.63, 130.46, 128.32, 125.54, 123.54, 122.18, 121.19, 120.08, 119.69, 117.90, 110.78, 103.30, 102.53, 42.70, 26.96, 17.70, 11.02; HRMS: calcd. for C_28_H_26_N_7_O [M+H] m/z 476.2199, found 476.2216.

#### N-methyl-1-((5-methyl-1H-pyrazol-3-yl)methyl)-2-(4-methyl-5-(2-methyl-1H-indol-5-yl)pyridin-3-yl)-1H-benzo[d]imidazole-5-carboxamide (33)

Brown solid; yield 41%; ^1^H NMR (500 MHz, DMSO) δ 12.41 (s, 1H), 10.78 (s, 1H), 8.71 (t, *J* = 2.5 Hz, 1H), 8.58 (t, *J* = 2.5 Hz, 1H), 8.42 (t, *J* = 4.5 Hz, 1H), 8.22 (d, *J* = 3.3 Hz, 1H), 7.80 (dd, *J* = 8.5, 3.0 Hz, 1H), 7.64 (dd, *J* = 8.7, 3.0 Hz, 1H), 7.48 (dd, *J* = 8.3, 3.1 Hz, 1H), 7.06 (d, *J* = 3.1 Hz, 1H), 6.92 (dd, *J* = 7.5, 3.1 Hz, 1H), 6.23 (s, 1H), 5.80 (d, *J* = 3.3 Hz, 1H), 5.29 (d, *J* = 3.1 Hz, 2H), 2.82 (d, *J* = 4.1 Hz, 3H), 2.36 (d, *J* = 3.2 Hz, 3H), 2.13 (d, *J* = 3.0 Hz, 3H), 1.91 (t, *J* = 2.5 Hz, 3H); ^13^C NMR (126 MHz, DMSO) δ 167.47, 152.30, 151.44, 149.54, 147.58, 146.33, 142.77, 139.74, 137.27, 137.24, 135.59, 134.88, 129.50, 129.29, 127.20, 122.47, 121.83, 120.42, 119.47, 119.27, 118.68, 111.53, 103.07, 100.15, 42.70, 26.85, 17.32, 13.86, 10.78; HRMS: calcd. for C_29_H_28_N_7_O [M+H] m/z 490.2355, found 490.2367.

#### N-methyl-1-((5-methyl-1H-pyrazol-3-yl)methyl)-2-(5-(5-phenylthiophen-2-yl)pyridin-3-yl)-1H-benzo[d]imidazole-5-carboxamide (44)

white solid; yield 60%; ^1^H NMR (500 MHz, DMSO) δ 12.56 (s, 1H), 9.12 (d, *J* = 3.7 Hz, 1H), 9.02 (s, 1H), 8.66 (d, *J* = 3.5 Hz, 1H), 8.46 (t, *J* = 4.5 Hz, 1H), 8.26 (d, *J* = 3.1 Hz, 1H), 7.88 – 7.78 (m, 2H), 7.74 (d, *J* = 7.8 Hz, 3H), 7.65 (d, *J* = 4.0 Hz, 1H), 7.51 – 7.43 (m, 2H), 7.37 (d, *J* = 6.6 Hz, 1H), 6.00 (s, 1H), 5.48 (d, *J* = 3.2 Hz, 2H), 2.83 (t, *J* = 4.0 Hz, 3H), 2.19 (d, *J* = 3.1 Hz, 3H); ^13^C NMR (126 MHz, DMSO) δ 167.37, 151.86, 148.84, 147.93, 147.26, 144.94, 142.59, 140.01, 138.47, 138.36, 133.66, 133.03, 129.83, 129.74, 129.60, 128.67, 127.63, 126.87, 125.91, 125.73, 122.88, 118.82, 111.47, 103.32, 43.13, 26.86, 10.84; HRMS: calcd. for C_29_H_25_N_6_OS [M+H] m/z 505.1811, found 505.1823.

#### N-methyl-1-((5-methyl-1H-pyrazol-3-yl)methyl)-2-(2-methyl-3-(5-phenylthiophen-2-yl)phenyl)-1H-benzo[d]imidazole-5-carboxamide (45)

white solid; yield 57%; ^1^H NMR (500 MHz, DMSO) δ 12.40 (s, 1H), 8.48 – 8.40 (m, 1H), 8.20 (s, 1H), 7.79 (d, *J* = 8.5 Hz, 1H), 7.72 (d, *J* = 7.7 Hz, 2H), 7.69 – 7.60 (m, 2H), 7.58 (q, *J* = 3.0 Hz, 1H), 7.53 (d, *J* = 7.5 Hz, 1H), 7.47 (q, *J* = 8.2 Hz, 3H), 7.36 (d, *J* = 7.7 Hz, 1H), 7.27 (d, *J* = 3.9 Hz, 1H), 5.68 (s, 1H), 5.18 (s, 2H), 2.82 (d, *J* = 4.0 Hz, 3H), 2.15 (d, *J* = 27.1 Hz, 6H); ^13^C NMR (126 MHz, DMSO) δ 167.51, 154.66, 147.45, 144.07, 142.62, 141.47, 139.66, 137.08, 136.42, 134.97, 133.94, 132.14, 131.78, 130.55, 129.68, 129.40, 129.16, 128.30, 126.43, 125.80, 124.60, 122.29, 118.56, 111.42, 102.91, 42.55, 26.85, 18.60, 10.79; HRMS: calcd. for C_31_H_28_N_7_OS [M+H] m/z 518.2015, found 518.2025.

#### 2-(3-methoxy-5-(5-phenylthiophen-2-yl)pyridin-2-yl)-N-methyl-1-((5-methyl-1H-pyrazol-3-yl)methyl)-1H-benzo[d]imidazole-5-carboxamide (46)

Light brown solid; yield 52%; ^1^H NMR (500 MHz, DMSO) δ 12.33 (s, 1H), 8.74 (s, 1H), 8.44 (s, 1H), 8.22 (s, 1H), 7.94 (s, 2H), 7.87 – 7.60 (m, 5H), 7.49 (d, *J* = 7.9 Hz, 2H), 7.40 (d, *J* = 7.7 Hz, 1H), 5.66 (s, 1H), 5.35 (s, 2H), 3.94 (s, 3H), 2.84 (s, 3H), 2.10 (s, 3H); ^13^C NMR (126 MHz, DMSO) δ 167.47, 155.69, 150.59, 147.85, 145.17, 142.54, 139.55, 138.40, 137.86, 137.76, 137.12, 133.66, 132.29, 129.76, 129.18, 128.70, 128.36, 125.90, 125.73, 122.58, 118.85, 116.44, 111.33, 102.83, 56.61, 42.57, 26.85, 10.76; HRMS: calcd. for C_30_H_27_N_6_O_2_S [M+H] m/z 535.1916, found 535.1922.

#### 2-(2-(dimethylamino)-4-(5-phenylthiophen-2-yl)phenyl)-N-methyl-1-((5-methyl-1H-pyrazol-3-yl)methyl)-1H-benzo[d]imidazole-5-carboxamide (47)

white solid; yield 68%; ^1^H NMR (500 MHz, DMSO) δ 12.40 (s, 1H), 8.49 (d, *J* = 5.4 Hz, 1H), 8.28 (s, 1H), 7.85 (d, *J* = 7.9 Hz, 3H), 7.80 (d, *J* = 3.7 Hz, 1H), 7.71 (d, *J* = 4.1 Hz, 1H), 7.66 (d, *J* = 8.8 Hz, 1H), 7.57 (h, *J* = 3.3 Hz, 3H), 7.46 (d, *J* = 7.2 Hz, 3H), 5.58 (s, 1H), 5.33 (s, 2H), 2.92 (t, *J* = 3.3 Hz, 3H), 2.71 (s, 6H), 2.18 (s, 3H); ^13^C NMR (126 MHz, DMSO) δ 167.62, 155.19, 152.58, 147.81, 143.67, 143.23, 142.66, 139.55, 137.44, 136.30, 134.13, 133.91, 129.69, 128.98, 128.38, 126.39, 125.76, 125.53, 122.07, 121.08, 118.43, 117.63, 114.23, 111.30, 102.61, 42.81, 26.85, 10.75; HRMS: calcd. For C_32_H_31_N_6_OS [M+H] m/z 547.2280, found 547.2282.

#### N-methyl-1-((5-methyl-1H-pyrazol-3-yl)methyl)-2-(6-(5-phenylthiophen-2-yl)-1H-indol-2-yl)-1H-benzo[d]imidazole-5-carboxamide (48)

Light brown solid; yield 50%; ^1^H NMR (500 MHz, DMSO) δ 12.52 (s, 1H), 12.13 (s, 1H), 8.47 (s, 1H), 8.22 (s, 1H), 7.88 – 7.69 (m, 6H), 7.60 – 7.31 (m, 7H), 5.91 (s, 1H), 5.71 (s, 2H), 2.85 (s, 3H), 2.17 (s, 3H); ^13^C NMR (126 MHz, DMSO) δ 166.36, 146.88, 146.73, 143.49, 141.41, 141.04, 138.91, 137.52, 136.65, 133.03, 128.55, 128.18, 127.19, 127.11, 126.98, 124.51, 124.36, 123.52, 121.51, 121.14, 117.51, 117.12, 109.90, 107.58, 103.38, 101.83, 41.91, 25.77, 9.71; HRMS: calcd. for C_32_H_27_N_6_OS [M+H] m/z 543.1967, found 543.1966.

#### 2-(5-(isoquinolin-5-yl)pyridin-3-yl)-N-methyl-1-((5-methyl-1H-pyrazol-3-yl)methyl)-1H-benzo[d]imidazole-5-carboxamide (49)

white solid; yield 50%; ^1^H NMR (500 MHz, DMSO) δ 12.39 (s, 1H), 9.48 – 9.40 (m, 1H), 9.22 (s, 1H), 8.91 (d, *J* = 3.1 Hz, 1H), 8.52 (q, *J* = 3.3 Hz, 2H), 8.44 (t, *J* = 4.5 Hz, 1H), 8.27 (dd, *J* = 12.3, 5.0 Hz, 2H), 7.92 (t, *J* = 5.1 Hz, 1H), 7.87 – 7.80 (m, 2H), 7.71 (dt, *J* = 9.0, 3.9 Hz, 2H), 5.92 (s, 1H), 5.53 (s, 2H), 2.82 (t, *J* = 4.2 Hz, 3H), 2.10 (d, *J* = 3.4 Hz, 3H); ^13^C NMR (126 MHz, DMSO) δ 167.34, 153.49, 152.01, 151.34, 149.54, 147.84, 144.41, 142.65, 139.86, 138.37, 137.91, 134.49, 134.37, 133.56, 132.46, 129.55, 129.01, 128.94, 127.77, 126.57, 122.84, 118.80, 117.82, 111.44, 103.19, 43.14, 26.85, 10.75; HRMS: calcd. for C_28_H_24_N_7_O [M+H] m/z 474.2053, found 474.2042.

#### 2-(3-(isoquinolin-5-yl)-2-methylphenyl)-N-methyl-1-((5-methyl-1H-pyrazol-3-yl)methyl)-1H-benzo[d]imidazole-5-carboxamide (50)

white solid; yield 68%; ^1^H NMR (500 MHz, DMSO) δ 12.37 (s, 1H), 9.42 (d, *J* = 3.1 Hz, 1H), 8.51 (t, *J* = 4.8 Hz, 1H), 8.41 (t, *J* = 4.6 Hz, 1H), 8.20 (dd, *J* = 10.3, 5.6 Hz, 2H), 7.80 (q, *J* = 6.6 Hz, 2H), 7.74 (t, *J* = 4.9 Hz, 1H), 7.66 (dt, *J* = 8.4, 3.7 Hz, 2H), 7.54 (t, *J* = 8.0 Hz, 1H), 7.44 (d, *J* = 7.6 Hz, 1H), 7.21 (t, *J* = 4.9 Hz, 1H), 5.71 (s, 1H), 5.25 (s, 2H), 2.81 (d, *J* = 4.0 Hz, 3H), 2.10 (s, 3H), 1.68 (s, 3H); ^13^C NMR (126 MHz, DMSO) δ 167.51, 154.62, 153.37, 144.06, 142.58, 139.61, 138.08, 137.15, 136.84, 134.10, 132.25, 131.58, 130.49, 129.15, 128.73, 127.97, 127.71, 126.32, 122.28, 118.56, 118.35, 111.33, 102.86, 42.56, 26.84, 17.71, 10.80; HRMS: calcd. for C_30_H_27_N_6_O [M+H] m/z 487.2246, found 487.2258.

#### 2-(5-(isoquinolin-5-yl)-3-methoxypyridin-2-yl)-N-methyl-1-((5-methyl-1H-pyrazol-3-yl)methyl)-1H-benzo[d]imidazole-5-carboxamide (51)

white solid; yield 68%; ^1^H NMR (500 MHz, DMSO) δ 12.29 (s, 1H), 9.40 (d, *J* = 2.9 Hz, 1H), 8.56 – 8.48 (m, 1H), 8.43 (d, *J* = 2.8 Hz, 1H), 8.38 (s, 1H), 8.23 (d, *J* = 8.3 Hz, 1H), 8.16 (s, 1H), 7.90 (d, *J* = 6.3 Hz, 1H), 7.78 (ddd, *J* = 20.4, 9.1, 5.3 Hz, 4H), 7.61 (s, 1H), 5.61 (d, *J* = 2.9 Hz, 1H), 5.33 (s, 2H), 3.82 (d, *J* = 2.9 Hz, 3H), 2.76 (d, *J* = 3.9 Hz, 3H), 2.03 (s, 3H); ^13^C NMR (126 MHz, DMSO) δ 167.51, 155.37, 153.47, 150.78, 144.44, 142.58, 141.65, 138.27, 137.13, 137.05, 134.65, 133.67, 132.42, 129.20, 129.00, 128.97, 127.72, 122.60, 121.87, 118.92, 118.13, 111.35, 103.00, 56.59, 40.59, 26.86; HRMS: calcd. for C_29_H_26_N_7_O_2_ [M+H] m/z 504.2158, found 504.2159.

#### 2-(2-(dimethylamino)-4-(isoquinolin-5-yl)phenyl)-N-methyl-1-((5-methyl-1H-pyrazol-3-yl)methyl)-1H-benzo[d]imidazole-5-carboxamide (52)

white solid; yield 50%; ^1^H NMR (500 MHz, DMSO) δ 12.32 (s, 1H), 9.43 (t, *J* = 2.7 Hz, 1H), 8.58 – 8.51 (m, 1H), 8.40 (t, *J* = 4.6 Hz, 1H), 8.21 (d, *J* = 10.0 Hz, 2H), 7.91 – 7.71 (m, 4H), 7.56 (ddd, *J* = 10.2, 7.9, 2.8 Hz, 2H), 7.24 – 7.13 (m, 2H), 5.51 (s, 1H), 5.30 (s, 2H), 2.86 – 2.76 (m, 3H), 2.67 – 2.55 (m, 6H), 2.09 (s, 3H); HRMS: calcd. for C_31_H_30_N_7_O [M+H] m/z 516.2512, found 516.2521.

#### 2-(6-(isoquinolin-5-yl)-1H-indol-2-yl)-N-methyl-1-((5-methyl-1H-pyrazol-3-yl)methyl)-1H-benzo[d]imidazole-5-carboxamide (53)

white solid; yield 72%; ^1^H NMR (500 MHz, DMSO) δ 12.54 (s, 1H), 12.16 (s, 1H), 9.43 (d, *J* = 2.6 Hz, 1H), 8.54 – 8.44 (m, 2H), 8.24 – 8.16 (m, 2H), 7.94 – 7.73 (m, 6H), 7.63 (s, 1H), 7.46 (s, 1H), 7.23 (dd, *J* = 8.3, 2.4 Hz, 1H), 5.92 (s, 1H), 5.76 (s, 2H), 2.85 (t, *J* = 3.4 Hz, 3H), 2.18 (d, *J* = 2.5 Hz, 3H); ^13^C NMR (126 MHz, DMSO) δ 167.74, 153.33, 147.97, 143.66, 142.45, 139.78, 138.58, 137.49, 134.09, 133.94, 131.70, 129.56, 129.14, 127.99, 127.89, 127.78, 127.50, 122.61, 122.53, 121.75, 118.73, 118.28, 113.44, 111.04, 104.23, 102.96, 42.84, 26.89, 10.91; HRMS: calcd. for C_31_H_26_N_7_O [M+H] m/z 512.2199, found 512.2206.

#### 2-(2-(isoquinolin-5-yl)pyridin-4-yl)-N-methyl-1-((1-methyl-1H-pyrazol-3-yl)methyl)-1H-benzo[d]imidazole-5-carboxamide (76)

white solid; yield 48%; ^1^H NMR (500 MHz, DMSO) δ 9.45 (s, 1H), 9.20 (s, 1H), 8.92 (s, 1H), 8.62 (s, 1H), 8.49 (dt, *J* = 33.6, 4.6 Hz, 2H), 8.34 – 8.15 (m, 2H), 8.03 – 7.65 (m, 5H), 7.58 (s, 1H), 6.17 (s, 1H), 5.55 (s, 2H), 3.58 (d, *J* = 3.1 Hz, 3H), 2.82 (t, *J* = 3.9 Hz, 3H); ^13^C NMR (126 MHz, DMSO) δ 167.42, 153.42, 152.03, 151.27, 149.55, 147.21, 144.21, 142.56, 138.29, 138.04, 134.45, 134.33, 133.55, 132.56, 132.44, 129.60, 129.00, 127.83, 126.45, 122.89, 118.84, 117.93, 111.41, 104.97, 49.08, 42.84, 26.86; HRMS: calcd. For C_28_H_24_N_7_O [M+H] m/z 474.2042, found 474.2049.

#### 2-(2-(isoquinolin-5-yl)pyridin-4-yl)-N-methyl-1-(thiazol-2-ylmethyl)-1H-benzo[d]imidazole-5-carboxamide (77)

white solid; yield 50%; ^1^H NMR (500 MHz, DMSO) δ 9.44 (d, *J* = 3.0 Hz, 1H), 9.14 (d, *J* = 2.9 Hz, 1H), 8.90 (d, *J* = 2.8 Hz, 1H), 8.55 – 8.36 (m, 3H), 8.27 (d, *J* = 9.8 Hz, 2H), 7.96 – 7.75 (m, 4H), 7.66 (dt, *J* = 17.2, 3.0 Hz, 3H), 6.09 (d, *J* = 3.1 Hz, 2H), 2.82 (d, *J* = 4.1 Hz, 3H); ^13^C NMR (126 MHz, DMSO) δ 166.16, 164.15, 152.40, 151.01, 150.44, 148.29, 143.31, 142.11, 141.55, 137.10, 136.68, 133.34, 133.29, 132.45, 131.35, 128.89, 127.89, 126.66, 125.20, 122.12, 120.69, 117.93, 116.69, 110.18, 44.85, 25.77; HRMS: calcd. for C_27_H_21_N_6_OS [M+H] m/z 477.1498, found 477.1507.

#### 2-(2-(isoquinolin-5-yl)pyridin-4-yl)-N-methyl-1-(oxazol-4-ylmethyl)-1H-benzo[d]imidazole-5-carboxamide (78)

white solid; yield 54%; ^1^H NMR (500 MHz, DMSO) δ 9.45 (d, J = 2.8 Hz, 1H), 9.28 (d, J = 2.7 Hz, 1H), 8.94 (d, J = 2.8 Hz, 1H), 8.63 (d, J = 2.5 Hz, 1H), 8.53 (dd, J = 6.0, 2.8 Hz, 1H), 8.45 (t, J = 4.2 Hz, 1H), 8.36 – 8.18 (m, 4H), 7.95 (d, J = 6.5 Hz, 1H), 7.89 – 7.80 (m, 2H), 7.80 – 7.68 (m, 2H), 5.57 (s, 2H), 2.82 (t, J = 3.7 Hz, 3H); ^13^C NMR (126 MHz, DMSO) δ 167.27, 153.50, 153.35, 152.24, 151.45, 149.64, 144.36, 142.71, 138.16, 138.00, 137.96, 135.40, 134.48, 134.41, 133.57, 132.47, 129.71, 129.02, 128.96, 127.79, 126.50, 122.88, 118.90, 117.85, 111.34, 40.76, 26.84; HRMS: calcd. for C_27_H_21_N_6_O_2_ [M+H] m/z 461.1726, found 461.1735.

#### 1-(cyclopropylmethyl)-2-(2-(isoquinolin-5-yl)pyridin-4-yl)-N-methyl-1H-benzo[d]imidazole-5-carboxamide (79)

Brown sticky solid; yield 58%; ^1^H NMR (500 MHz, DMSO) δ 9.46 (d, *J* = 2.8 Hz, 1H), 9.15 (d, *J* = 2.8 Hz, 1H), 8.94 (q, *J* = 2.5 Hz, 1H), 8.57 (dt, *J* = 5.3, 2.3 Hz, 1H), 8.45 (dt, *J* = 14.5, 3.5 Hz, 2H), 8.33 – 8.22 (m, 2H), 7.98 – 7.91 (m, 1H), 7.85 (qd, *J* = 8.3, 4.6 Hz, 3H), 7.72 (dd, *J* = 6.0, 2.7 Hz, 1H), 4.41 – 4.32 (m, 2H), 2.83 (t, *J* = 3.8 Hz, 3H), 1.17 – 1.09 (m, 1H), 0.49 – 0.39 (m, 2H), 0.26 (t, *J* = 4.1 Hz, 2H); ^13^C NMR (126 MHz, DMSO) δ 167.32, 153.55, 151.84, 151.42, 149.43, 144.43, 142.69, 138.35, 137.89, 134.50, 134.44, 133.64, 132.52, 129.41, 129.03, 128.97, 127.81, 127.00, 122.81, 118.86, 117.80, 111.49, 48.97, 26.85, 11.83, 4.38; HRMS: calcd. for C_27_H_24_N_5_O [M+H] m/z 434.1981, found 434.2000.

#### 2-(2-(isoquinolin-5-yl)pyridin-4-yl)-N-methyl-1-(4-methylbenzyl)-1H-benzo[d]imidazole-5-carboxamide (80)

white solid; yield 72%; ^1^H NMR (500 MHz, CDCl_3_) δ 9.36 (s, 1H), 8.98 (d, *J* = 3.3 Hz, 1H), 8.85 (d, *J* = 3.2 Hz, 1H), 8.61 – 8.33 (m, 1H), 8.28 – 8.03 (m, 3H), 7.89 – 7.53 (m, 4H), 7.35 (dd, *J* = 8.3, 3.4 Hz, 1H), 7.08 (dd, *J* = 7.7, 3.4 Hz, 2H), 6.98 – 6.84 (m, 2H), 6.39 (s, 1H), 5.52 (d, *J* = 3.6 Hz, 2H), 3.04 (t, *J* = 4.4 Hz, 3H), 2.28 (d, *J* = 3.4 Hz, 3H); HRMS: calcd. for C_31_H_26_N_5_O_2_ [M+H] m/z 484.2137, found 484.2149.

#### 2-(2-(isoquinolin-5-yl)pyridin-4-yl)-N-methyl-1-(4-(methylthio)benzyl)-1H-benzo[d]imidazole-5-carboxamide (81)

Off-white solid; yield 65%; ^1^H NMR (500 MHz, DMSO) δ 9.47 – 9.41 (m, 1H), 9.06 (d, *J* = 2.6 Hz, 1H), 8.88 (q, *J* = 2.4 Hz, 1H), 8.54 – 8.40 (m, 2H), 8.34 – 8.14 (m, 3H), 7.83 (dd, *J* = 8.5, 3.5 Hz, 3H), 7.73 – 7.63 (m, 1H), 7.55 (t, *J* = 4.7 Hz, 1H), 7.14 (dt, *J* = 8.4, 2.2 Hz, 2H), 7.00 – 6.90 (m, 2H), 5.70 (d, *J* = 3.2 Hz, 2H), 2.82 (t, *J* = 3.8 Hz, 3H), 2.37 (t, *J* = 2.3 Hz, 3H); ^13^C NMR (126 MHz, DMSO) δ 167.30, 153.47, 152.22, 151.46, 149.16, 144.40, 142.71, 138.47, 138.06, 137.67, 134.43, 134.30, 133.65, 133.49, 132.43, 129.79, 128.95, 127.71, 127.31, 126.71, 126.52, 123.13, 119.03, 117.68, 111.32, 47.79, 26.86, 15.04; HRMS: calcd. for C_31_H_26_N_5_OS [M+H] m/z 516.1858, found 516.1866.

#### 2-(2-(isoquinolin-5-yl)pyridin-4-yl)-N-methyl-1-((5-methylpyridin-2-yl)methyl)-1H-benzo[d]imidazole-5-carboxamide (82)

Light brown solid; yield 40%; ^1^H NMR (500 MHz, DMSO) δ 9.43 (s, 1H), 9.15 (s, 1H), 8.88 (s, 1H), 8.60 – 8.36 (m, 3H), 8.36 – 8.14 (m, 3H), 7.83 (dt, *J* = 23.2, 7.4 Hz, 3H), 7.69 – 7.51 (m, 3H), 7.24 (d, *J* = 7.9 Hz, 1H), 5.74 (s, 2H), 2.82 (d, *J* = 4.6 Hz, 3H), 2.18 (s, 3H); ^13^C NMR (126 MHz, DMSO) δ 167.34, 153.48, 153.04, 152.49, 151.30, 150.18, 149.40, 144.34, 142.69, 138.57, 137.97, 137.81, 134.38, 134.26, 133.45, 132.69, 132.37, 129.63, 128.97, 128.93, 127.74, 126.62, 122.92, 121.88, 118.90, 117.68, 111.22, 49.49, 26.85, 17.96; HRMS: calcd. for C_30_H_25_N_6_O [M+H] m/z 485.2090, found 485.2098.

#### MST assay

MST analysis was performed as previously reported.^11–13^ Briefly, compounds showing an apparent interaction in the initial single-concentration screen were advanced to MSTfor dose-dependent analysis. Each compound was tested in a 12-point dilution series, with concentrations extending from 250 μM to the low-nanomolar range, in the presence of fluorescently labeled NOS1AP-His protein. The final DMSO concentration was 2.5%. Following 30 minutes of equilibration at room temperature, samples were transferred to standard MST capillaries and analyzed on a Monolith X instrument. Measurements used medium-to-high MST power and 60-80% LED excitation with red-channel detection.

#### SPR Analysis of NOSTRA Binding to NOS1AP

Direct binding of NOSTRA to recombinant human NOS1AP was characterized SPR using a Biacore 8K instrument (Cytiva). NOS1AP was immobilized on a Series S CM5 sensor chip by standard amine coupling, with a reference flow cell subjected to identical activation and blocking procedures without protein immobilization. Binding experiments were conducted at 25 °C in HBS-P+ running buffer (10 mM HEPES, 150 mM NaCl, 0.05% surfactant P20, pH 7.4) supplemented with 1% DMSO. NOSTRA was serially diluted to concentrations of 0.125-8 µM and injected at 30 µL/min, with association and dissociation phases of 120 and 300 s, respectively. Sensorgrams were corrected by reference-channel subtraction, blank-buffer subtraction, and DMSO solvent correction. Binding responses were globally fitted to a 1:1 Langmuir interaction model to determine association (k_on_) and dissociation (k_off_) rate constants and calculate the equilibrium dissociation constant (*K*_D_ = k_off/_k_on_). Independent steady-state affinity analysis was performed using equilibrium binding responses. Three independent experiments were conducted, and affinity estimates were expressed as mean ± SD.

#### NOS1AP Protein-Protein Interaction Selectivity Assay

The selectivity of NOSTRA toward NOS1AP-associated protein complexes was evaluated using a homogeneous time-resolved fluorescence (HTRF) proximity assay. Six established NOS1AP interaction partners were examined individually: SCRIB, RASD1 (Dexras1), SYN1 (synapsin I), NOS1 (nNOS), CPE (carboxypeptidase E), and MAPT (tau). Recombinant His-tagged NOS1AP (50 nM) and FLAG-tagged interaction partners (100 nM) were combined in assay buffer containing 25 mM HEPES (pH 7.4), 150 mM NaCl, 0.01% Tween-20, and 0.1% bovine serum albumin. Protein mixtures were incubated with NOSTRA (0, 0.3, 1, 3, or 10 µM) for 60 min at room temperature in white 384-well plates, maintaining a constant final DMSO concentration of 0.5%. Terbium-labeled anti-His donor and d2-labeled anti-FLAG acceptor reagents were subsequently added, followed by an additional 60-min incubation protected from light. Time-resolved fluorescence was measured at 620 and 665 nm following excitation at 337 nm on Tecan Spark. HTRF ratios were calculated from acceptor and donor emission intensities, corrected for background fluorescence, and normalized to the corresponding vehicle-treated protein pair. Single-protein, compound-only, and single-label controls were included to assess nonspecific proximity and fluorescence interference. Each condition was evaluated in triplicate.

#### SPR Analysis of NOS1AP–SCRIB Cooperativity

The effect of NOSTRA on NOS1AP-SCRIB binding affinity and kinetics was investigated using a Biacore 8K SPR system (Cytiva). Recombinant human NOS1AP was immobilized on a Series S CM5 sensor chip by amine coupling. Increasing concentrations of recombinant SCRIB (0.39-50 µM) were injected in the absence or presence of NOSTRA at 0.3, 1, 3, or 10 µM. Binding measurements were conducted at 25 °C in HBS-P+ buffer supplemented with 0.5% DMSO. To maintain constant compound exposure during the binding measurements, NOSTRA was included at the corresponding concentration in both analyte and running buffers. SCRIB and NOSTRA were preincubated for 30 min before injection, and association and dissociation phases were monitored for 180 and 600 s, respectively, at a flow rate of 30 µL/min. Reference-subtracted sensorgrams were analyzed to determine apparent equilibrium dissociation constants and, where supported by the data, kinetic association and dissociation rate constants. Compound-dependent enhancement of binding affinity was quantified using the apparent cooperativity factor α = *K*_D_,vehicle/ *K*_D_,NOSTRA. Six independent experiments were performed, and results were expressed as mean ± SD.

#### NOS1AP-SCRIB TR-FRET Proximity Assay

NOSTRA-dependent enhancement of NOS1AP-SCRIB association was independently assessed using a time-resolved fluorescence resonance energy transfer (TR-FRET) assay. Recombinant His-tagged NOS1AP (25 nM) and FLAG-tagged SCRIB (50 nM) were combined in assay buffer containing 25 mM HEPES (pH 7.4), 150 mM NaCl, 0.01% Tween-20, and 0.1% bovine serum albumin. Reactions were assembled in white 384-well low-volume plates at a final volume of 20 µL per well. NOSTRA was tested over a concentration range of 0.03-30 µM, with DMSO maintained at 0.5% across all conditions. Following a 60-min incubation at room temperature, terbium-labeled anti-His donor and d2-labeled anti-FLAG acceptor reagents were added, and the mixtures were incubated for an additional 60 min protected from light. Fluorescence was recorded on Tecan Spark using excitation at 337 nm and emission detection at 620 and 665 nm with time-resolved acquisition. TR-FRET ratios were calculated from acceptor-to-donor emission intensities, corrected for background fluorescence, and normalized to vehicle-treated controls. Single-protein, single-label, and compound-only controls were included to evaluate nonspecific signals and fluorescence interference. Concentration-dependent changes in proximity signals were analyzed to assess NOSTRA-mediated enhancement of NOS1AP-SCRIB association.

#### Native Mass Spectrometry of Ternary Complex Formation

Native mass spectrometry was employed to characterize NOSTRA-dependent assembly of the NOS1AP-SCRIB complex under nondenaturing conditions. Recombinant human NOS1AP and SCRIB were buffer-exchanged into 150 mM ammonium acetate (pH 7.4) using centrifugal ultrafiltration to remove nonvolatile salts and detergents. The proteins were combined at equimolar concentrations (5 µM each) and incubated with NOSTRA at 0, 0.3, 1, 3, 10, or 30 µM for 30 min at room temperature, maintaining a constant DMSO concentration not exceeding 0.5%. Samples were analyzed by native nanoelectrospray ionization mass spectrometry. Mass spectra were acquired to detect free NOS1AP, free SCRIB, NOS1AP-SCRIB binary complexes, NOS1AP-NOSTRA complexes, and the NOS1AP-NOSTRA-SCRIB ternary assembly. Molecular species were assigned based on their expected intact masses and corresponding charge-state distributions. Relative abundances were determined by integrating the signals associated with each assigned species and normalizing to the summed intensity of all assigned species within each spectrum. Compound-dependent changes in ternary-complex abundance were evaluated across the NOSTRA concentration series.

#### Structure preparation for computational study

The AlphaFold model of human NOS1AP (UniProt O75052; model AF-O75052-F1) and its predicted aligned error matrix were retrieved from the AlphaFold Protein Structure Database. Confidently folded segments were defined as contiguous runs of residues with pLDDT above 70, merging gaps of up to four residues and discarding runs shorter than eight residues; secondary-structure content was estimated from Cα(i)–Cα(i+4) distances. Structures were prepared in Maestro (Schrödinger Suite 2026-3) with the Protein Preparation Workflow: hydrogen addition, assignment of protonation states at pH 7.4, and restrained minimisation. The isolated PTB domain (residues 15-176) was capped with acetyl and N-methylamide groups and used for all subsequent calculations.

#### Homolog superposition and the composite peptide-binding groove

Experimental PTB-domain structures of homologous proteins (Supplementary Table S1) were retrieved from the Protein Data Bank and superposed onto the model with the structalign utility of Schrödinger Suite 2026-3, taking the first deposited model for NMR entries. Sequence identity to NOS1AP was computed by local pairwise alignment (BLOSUM62, gap penalties −11 and −1) between the NOS1AP PTB sequence and the longest chain of each superposed structure, over the aligned region. In each peptide complex, chains of 40 residues or fewer were treated as the bound peptide, which excludes additional copies of the PTB domain present in some crystals. Peptides were transferred into the model coordinate frame and the model residues with any heavy atom within 5.0 Å of a transferred peptide were recorded. Residues meeting this criterion in at least two of the five peptide complexes were retained as the composite peptide-binding groove (24 residues); 15 residues met it in all five.

#### Pocket identification and matched control patches

Candidate pockets were identified with SiteMap on three receptor definitions: the full-length model, the isolated PTB domain and the folded helical segment. Pockets whose residues had a mean pLDDT below 70, or in which more than 30% of residues fell below pLDDT 70, were not considered further. Control patches were generated on the prepared PTB domain with a fixed random seed. A solvent-exposed residue (side-chain SASA at least 15 Å²) was chosen as a centre at least 10 Å from the centre of any candidate pocket, the composite groove or another control, and the patch was grown over search radii of 5.0–16.0 Å until its residue count matched the paired pocket to within three residues. Patches were required to share no residue with the pocket they controlled, to overlap other controls by at most 25% of residues, to have a mean solvent-accessible surface area within 0.5-to 2.0-fold of the pocket, and to satisfy the same pLDDT criterion applied to pockets. Where no patch satisfied the surface-area criterion it was relaxed to 3.0-fold and the relaxation recorded.

#### Mixed-solvent molecular dynamics

Mixed-solvent molecular dynamics was performed on the prepared PTB domain using the MixMD implementation of Schrödinger Suite 2026-3 with the OPLS4 force field. Three heteroaromatic cosolvent probes representing chemical features of the compound series, isoquinoline, benzothiazole and pyridine, were simulated independently. Ten simulations were run per probe, each comprising 15 ns of equilibration followed by 5 ns of unrestrained production, giving 600 ns simulated in total of which the 150 ns of production sampling was analyzed, on a single NVIDIA RTX 4070 Ti SUPER. Hotspots and their constituent residues were taken from the standard MixMD occupancy analysis. Occupancy is reported for each hotspot as its score expressed as a percentage of the sum over all hotspots, both for the pooled data and for each probe separately. Probe selection was based on the chemical structures of the series and made no use of the measured affinities.

#### Structural characterization of the hotspot

Relative solvent accessibility was computed on the prepared PTB domain with an in-house implementation of the Shrake–Rupley algorithm (200 sphere points, 1.4 Å probe radius) and normalised to the theoretical maximum accessible surface areas of Tien et al. (2013); residues above 0.25 were classified as solvent-exposed. Side-chain hydrogen-bond acceptors (Asp, Glu, Asn, Gln, Ser, Thr, Tyr, His) and donors (Arg, Lys, His, Ser, Thr, Tyr, Asn, Gln, Trp) were assigned by residue type, so that His, Ser, Thr, Tyr, Asn and Gln count as both. Distances between exposed acceptors were measured between side-chain acceptor atoms. Overlap between the hotspot, the composite groove, the SiteMap pockets and the control patches was computed at the residue level and is reported in both directions where the sets differ in size.

#### Human iPSC-Derived Cortical Neuron Studies

Human induced pluripotent stem cell (iPSC)-derived cortical neurons representing three independent healthy donors were obtained from FUJIFILM Cellular Dynamics International (Madison, WI, USA) and maintained in neuronal culture medium according to the manufacturer’s recommendations. Neurons were plated on poly-D-lysine/laminin-coated culture plates and matured for approximately 4 weeks before treatment. NOSTRA was administered at final concentrations of 0, 0.1, 0.3, 1, 3, and 10 µM, with DMSO maintained at 0.1% (v/v). Following 48 h of treatment, endogenous NOS1AP-SCRIB association was assessed using HTRF proximity assay with antibodies recognizing the respective endogenous proteins. Rac1 activation was quantified using a Rac1-GTP G-LISA activation assay (Cytoskeleton, Inc., Denver, CO, USA), with signals normalized to total protein content. Cell viability was assessed using CellTiter-Glo Luminescent Cell Viability Assay (Promega), and membrane damage was quantified using a lactate dehydrogenase (LDH) release assay, expressed relative to maximum LDH release from lysed cells. Spontaneous neuronal activity was recorded using a multielectrode array (MEA) system (Axion BioSystems, Atlanta, GA, USA) after treatment. Mean firing rate, network burst frequency, and synchrony were calculated using the manufacturer’s analysis software. Experiments were performed using three independent donor-derived neuronal preparations, with technical measurements averaged within each donor before statistical analysis.

#### AD Patient-Derived iPSC Cortical Neuron Studies

Human iPSC-derived cortical neurons from three independent AD donors were obtained from FUJIFILM Cellular Dynamics (International, Madison, WI, USA) and cultured on poly-D-lysine/laminin-coated plates in neuronal maintenance medium. The samples were obtained from three sporadic AD donors aged 68, 74, and 79 years (female, male, and female, respectively). Neurons were allowed to mature for approximately 4 weeks before treatment. AD-derived neurons were exposed to NOSTRA (0, 0.1, 0.3, 1, 3, or 10 µM) for 48 h. All comparisons are against vehicle-treated neurons. Endogenous NOS1AP-SCRIB association was quantified using an HTRF proximity assay, and Rac1 activation was determined by measuring GTP-bound Rac1 using a G-LISA assay (Cytoskeleton, Inc.). Amyloid-β40 and amyloid-β42 concentrations in conditioned medium were measured using commercially available ultrasensitive immunoassays, and the Aβ42/Aβ40 ratio was calculated. Phosphorylated tau (pTau217 and pTau181), total tau, and neurofilament light chain (NfL) were quantified using analyte-specific immunoassays (Quanterix, Billerica, MA, USA). Cell viability and membrane integrity were evaluated using CellTiter-Glo (Promega) and LDH release assays, respectively. Spontaneous neuronal activity was assessed using an Axion BioSystems MEA platform, with mean firing rate, burst frequency, network burst frequency, synchrony, and percentage of active electrodes extracted from recorded activity. Measurements were performed across three independent AD donor lines, with donor-level means used as biological replicates for statistical analysis.

## In Vitro Physicochemical, Metabolic, Permeability, and Safety Profiling

The physicochemical properties, metabolic stability, permeability, and preliminary safety profile of NOSTRA were characterized using established in vitro ADME assays. Kinetic and thermodynamic aqueous solubility were determined at pH 7.4 by LC-MS/MS quantification following compound equilibration and removal of insoluble material. The distribution coefficient (logD_7.4_) was measured using a shake-flask partitioning assay, and the apparent pKa was determined by potentiometric titration. Metabolic stability was evaluated in pooled human and mouse liver microsomes (0.5 mg/mL) supplemented with 1 mM NADPH at 37 °C, with aliquots collected over 60 min for LC-MS/MS analysis. Intrinsic clearance was additionally determined in cryopreserved human and mouse hepatocytes. Plasma stability was evaluated in human plasma over 120 min, while plasma protein binding and brain tissue binding were determined by equilibrium dialysis. Passive blood–brain barrier permeability was assessed using a PAMPA-BBB assay, and bidirectional permeability was measured in MDCK-MDR1 and BCRP transporter-expressing cell monolayers. Apparent permeability coefficients (Papp) and efflux ratios were calculated from donor-to-receiver transport measurements. Cytochrome P450 inhibition was evaluated against CYP1A2, CYP2C9, CYP2C19, CYP2D6, and CYP3A4 using isoform-selective probe substrates, with time-dependent CYP3A4 inhibition assessed following NADPH-dependent preincubation. hERG channel inhibition was assessed using an automated patch-clamp assay, and cellular cytotoxicity was determined in HepG2 cells and primary human hepatocytes. Potential reactive metabolite formation was evaluated by glutathione trapping followed by LC-MS/MS analysis. Assays were conducted in triplicate, and results were expressed as mean ± SD.

## Software and data availability

Schrödinger Suite 2026-3 (Maestro, SiteMap, MixMD), RDKit 2023.09.6 and Biopython were used. Coordinates of the prepared NOS1AP PTB domain model and of the model annotated with HS1 probe-contact frequency are provided as Supporting Information (PDB).

## Supporting Information

Supporting Information containing chemical characterization of key compounds including NMR spectrum, HRMS and HPLC data (PDF)

## Author Information

**Corresponding Author: Moustafa Gabr** - Department of Radiology, Columbia University, New York, NY 10032, USA;

## Author Contributions

B.K. designed, synthesized, and characterized the compounds. A.A. performed the biophysical studies. S.C. performed computational studies and cellular assays. M.G. conceptualized and supervised the project and secured funding. B.K., A.A., S.C., and M.G. contributed to writing and editing the manuscript.

## Notes

The author declare no conflict of interests.

