## Supporting Information for "DNA-Encoded Library-Guided Discovery of a First-in-Class Molecular Glue of the NOS1AP-SCRIB Complex in Alzheimer’s Disease Patient-Derived Neurons"

**Table of Contents**

| 1. ^1^H NMR, ^13^C NMR of compounds | S2-S29 |
| --- | --- |
| 2. HRMS of compounds | S30-S39 |
| 3. HPLC purity Chromatograms of **10**, **13**, **29**, **31**, and **49** | S39-S40 |
| 4. MST binding curve for compound 10 | S40 |
| 5. Per-probe hotspot occupancy | S41 |
| 6. Experimental PTB-domain structures | S42 |

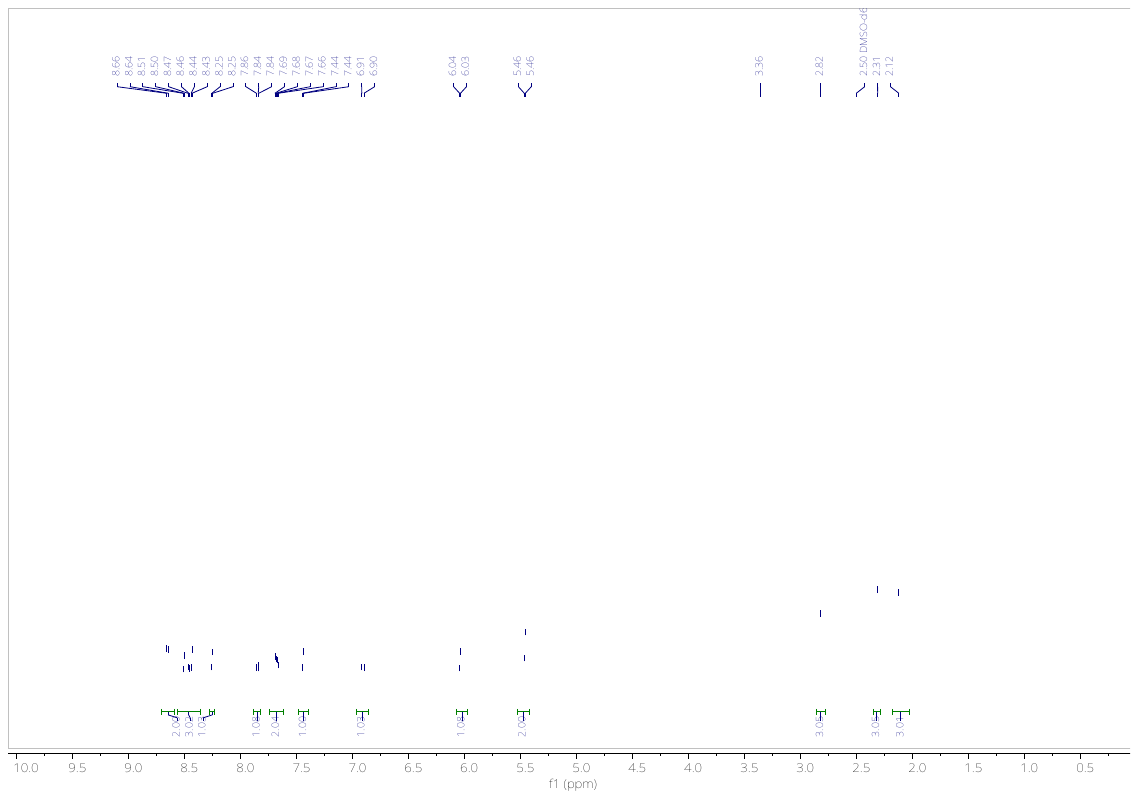

**Fig. 1**. 1H NMR of **10** in DMSO-*d*_6_.

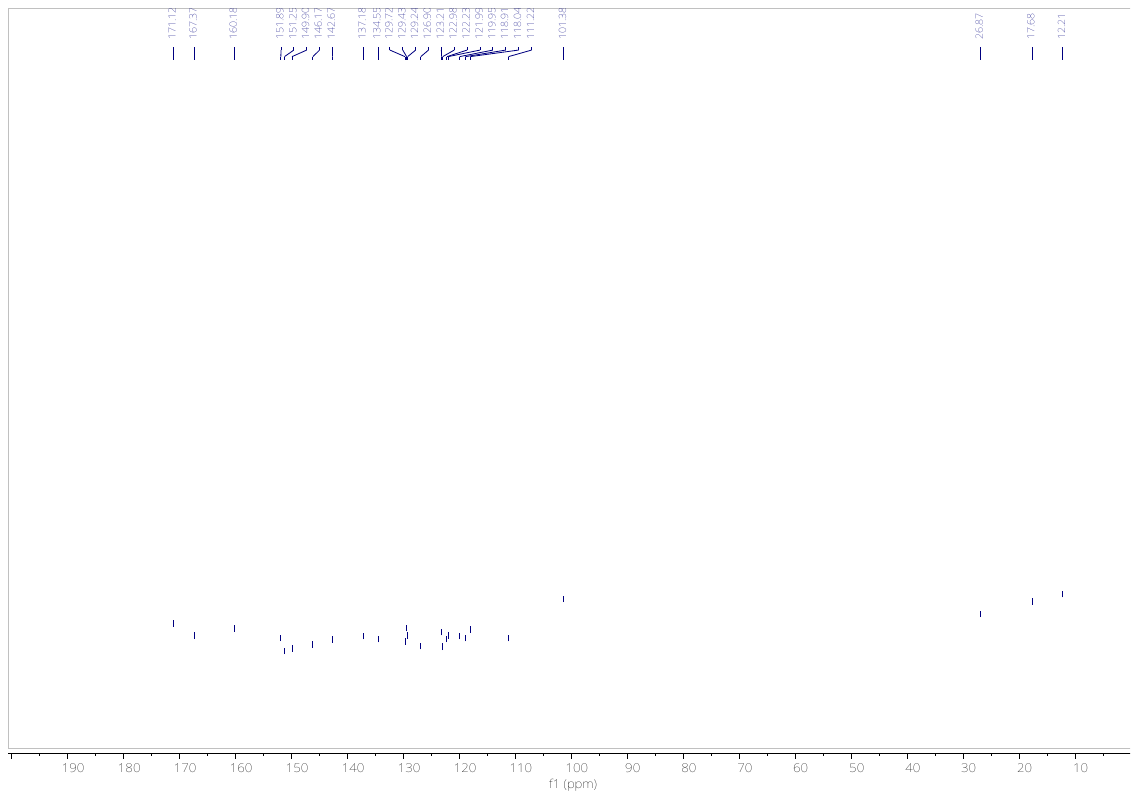

**Fig. 2**. ^13^C NMR of compound **10** in DMSO-*d*_6_.

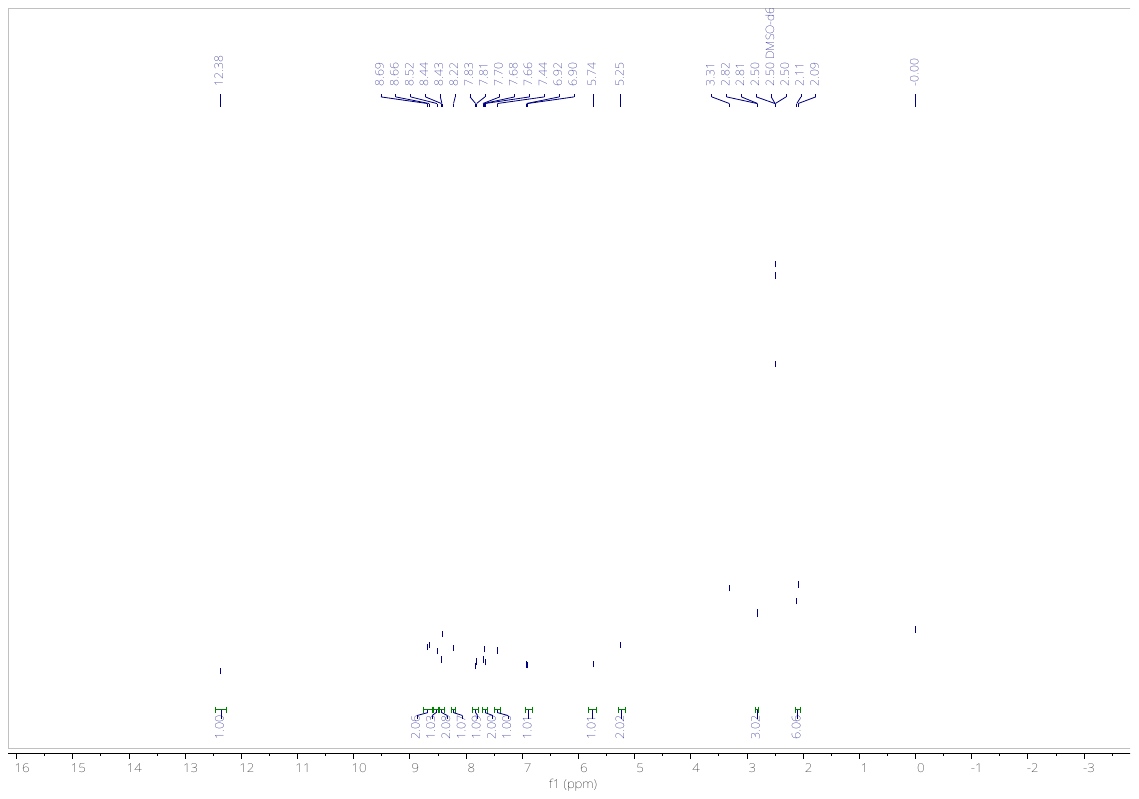

**Fig. 3**. ^1^H NMR of compound **13** in DMSO-*d*_6_.

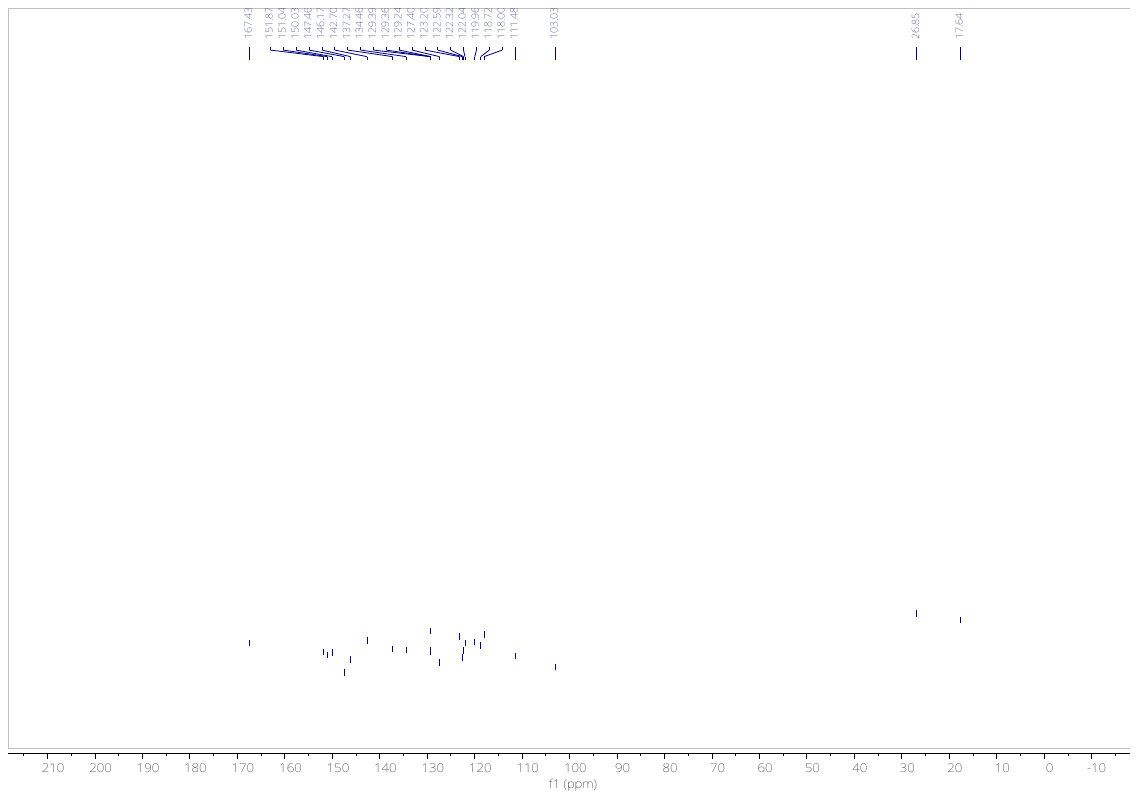

**Fig. 4**. ^13^C NMR of compound **13** in DMSO-*d*_6_.

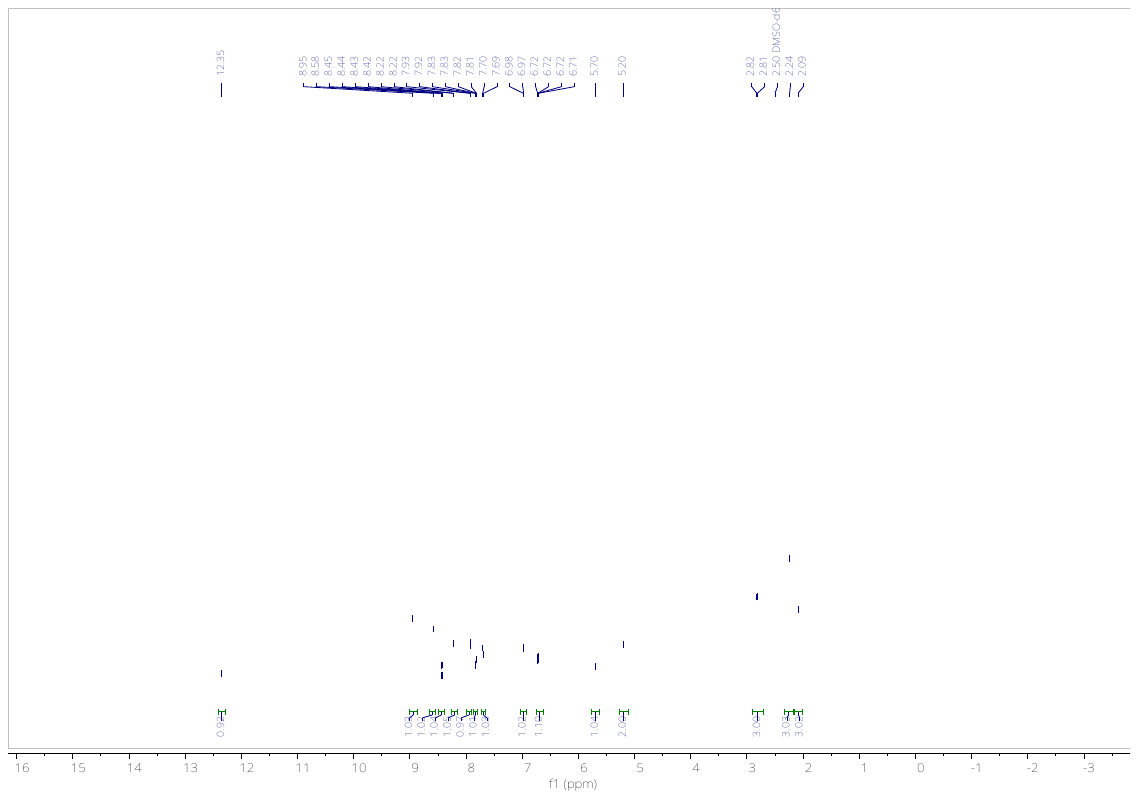

**Fig. 5**. ^1^H NMR of compound **24** in DMSO-*d*_6_.

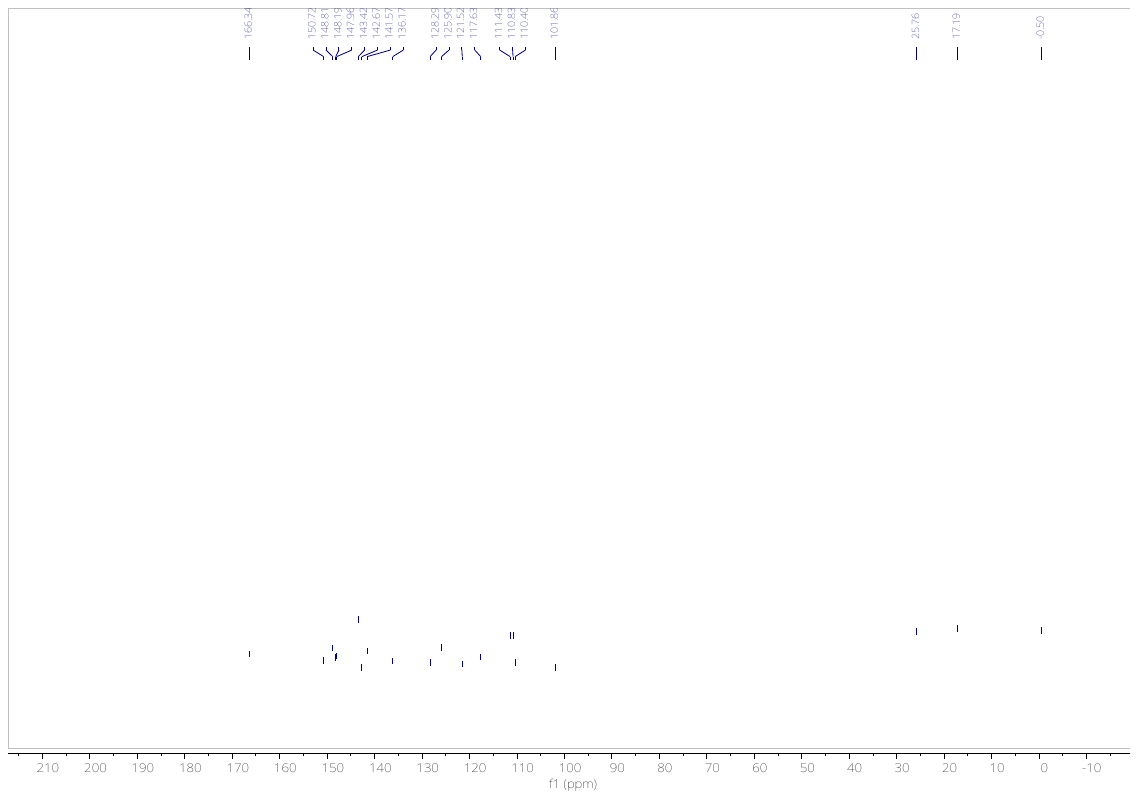

**Fig. 6**. ^13^C NMR of compound **24** in DMSO-*d*_6_.

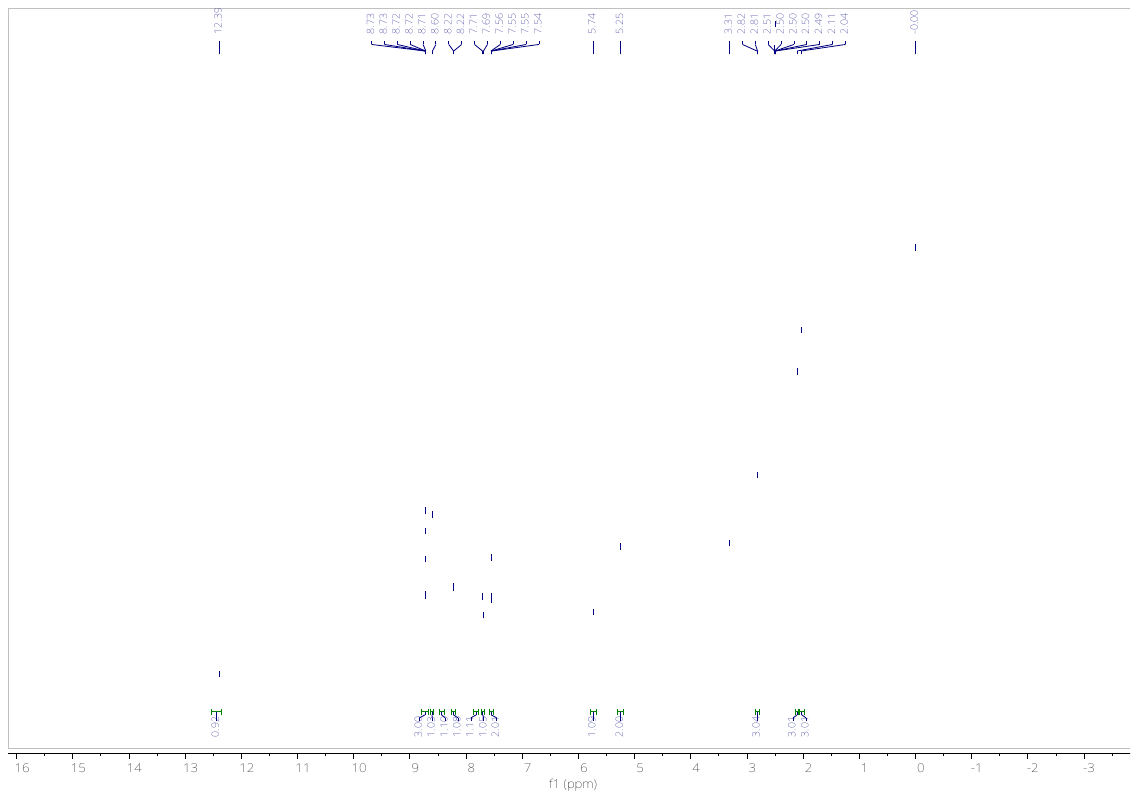

**Fig. 7**. ^1^H NMR of compound **25** in DMSO-*d*_6_.

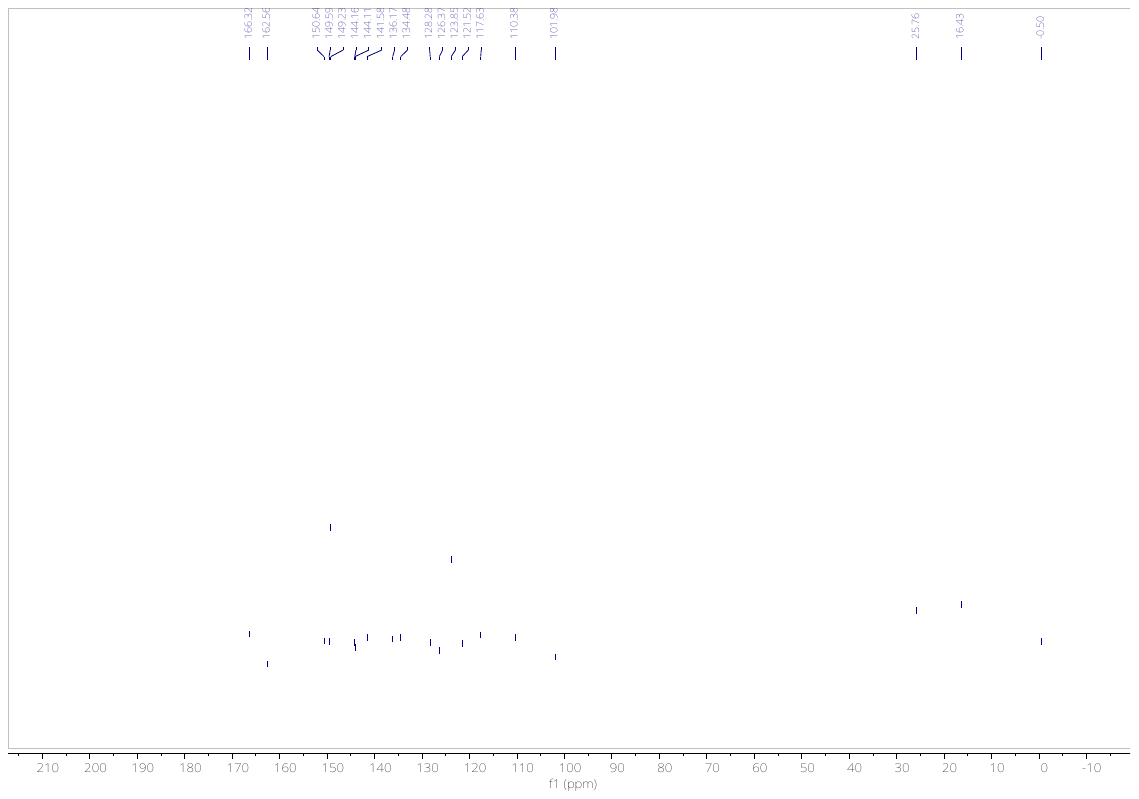

**Fig. 8**. ^13^C NMR of compound **25** in DMSO-*d*_6_.

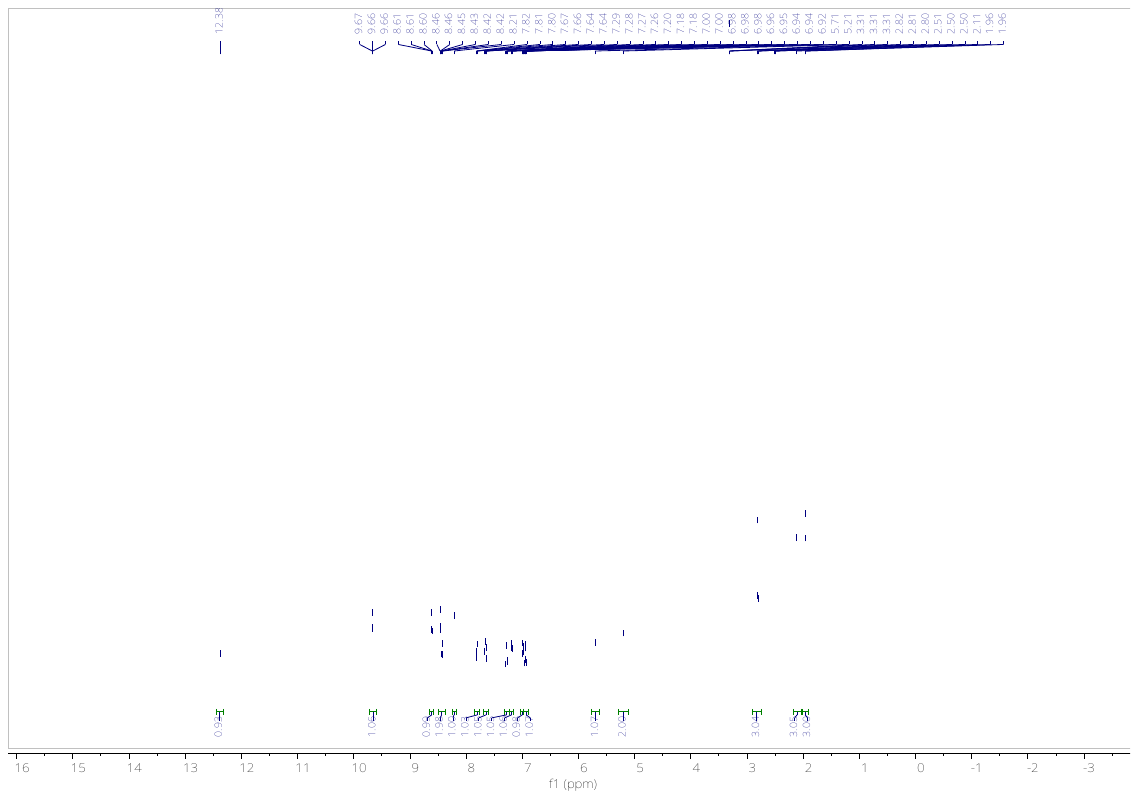

**Fig. 9**. ^1^H NMR of compound **26** in DMSO-*d*_6_.

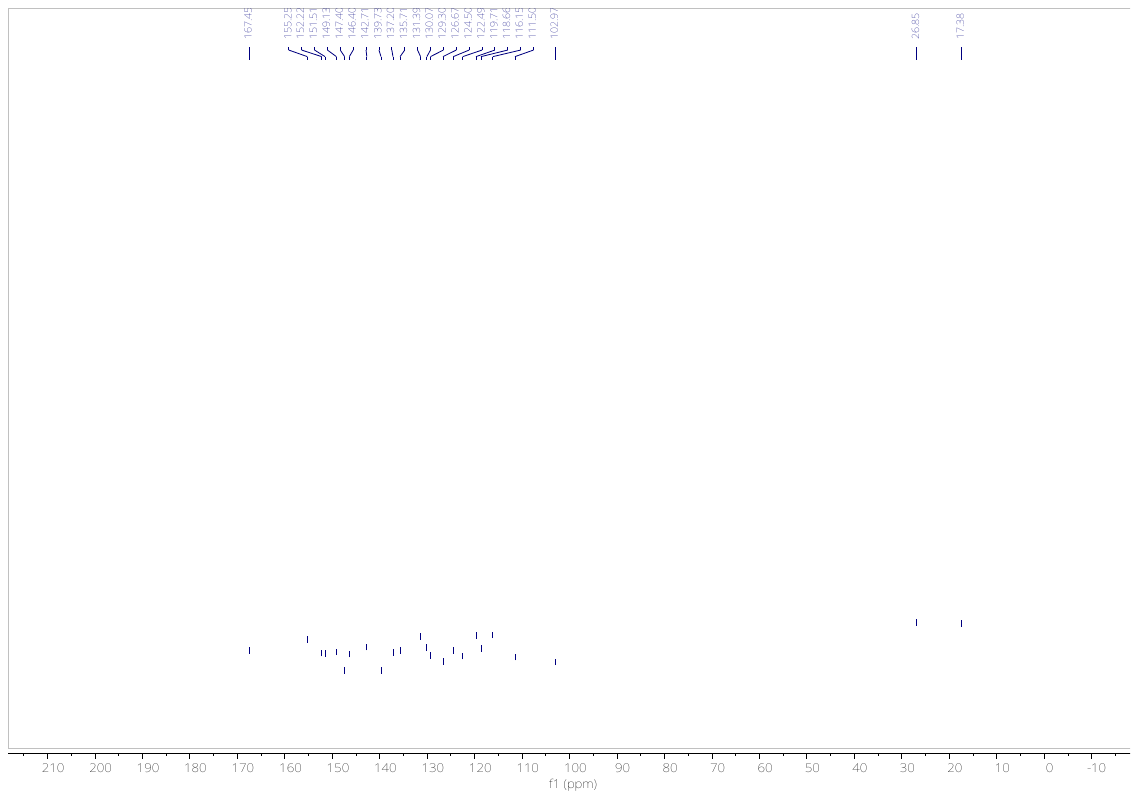

**Fig. 10**. ^13^C NMR of compound **26** in DMSO-*d*_6_.

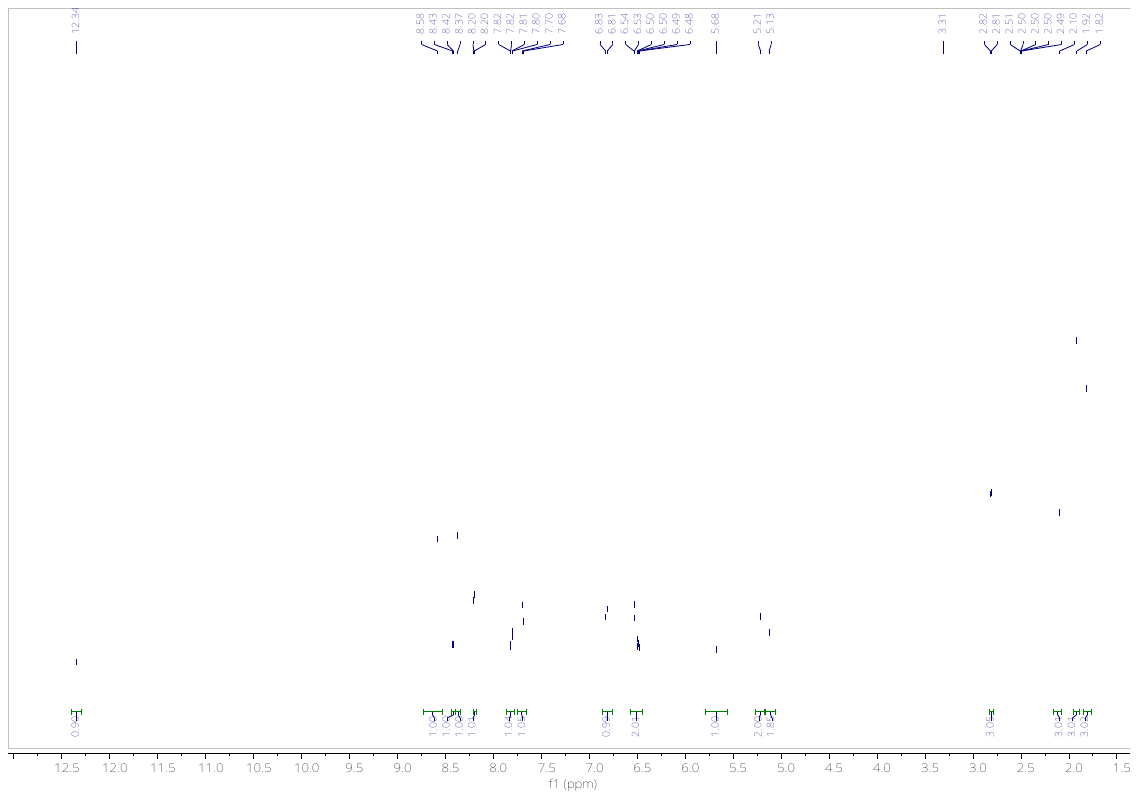

**Fig. 11**. ^1^H NMR of compound **27** in DMSO-*d*_6_.

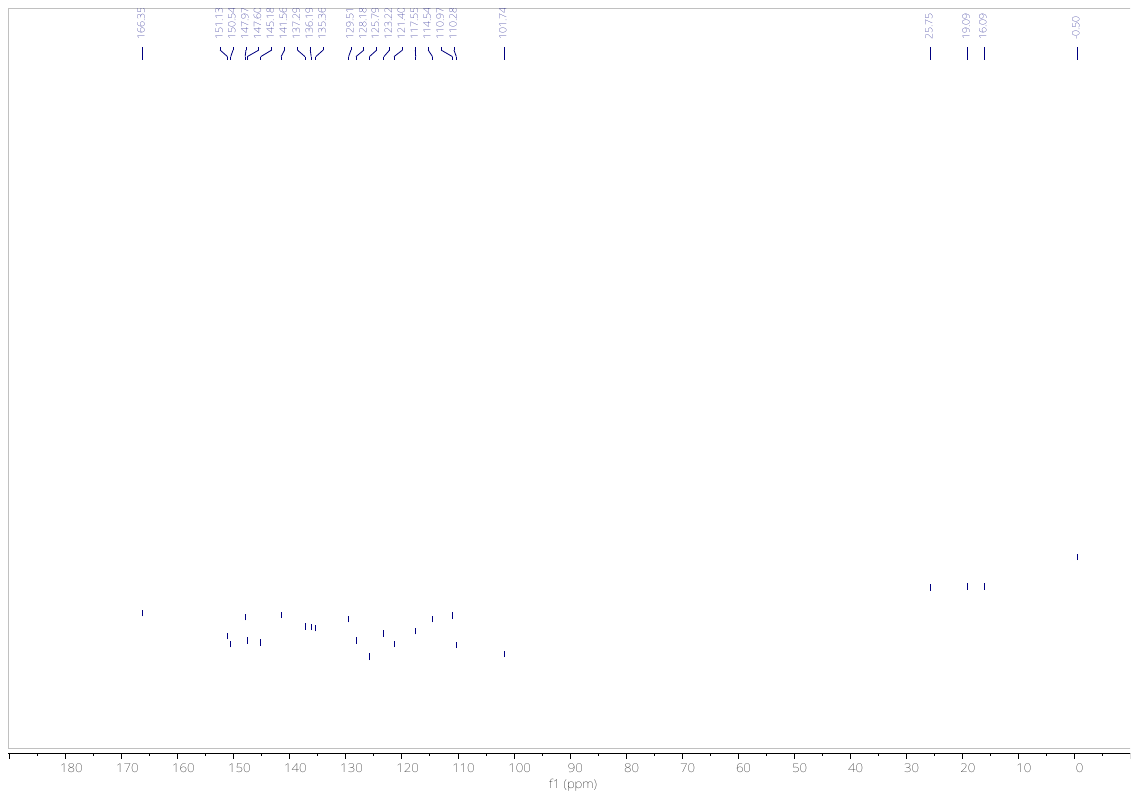

**Fig. 12**. ^13^C NMR of compound **27** in DMSO-*d*_6_.

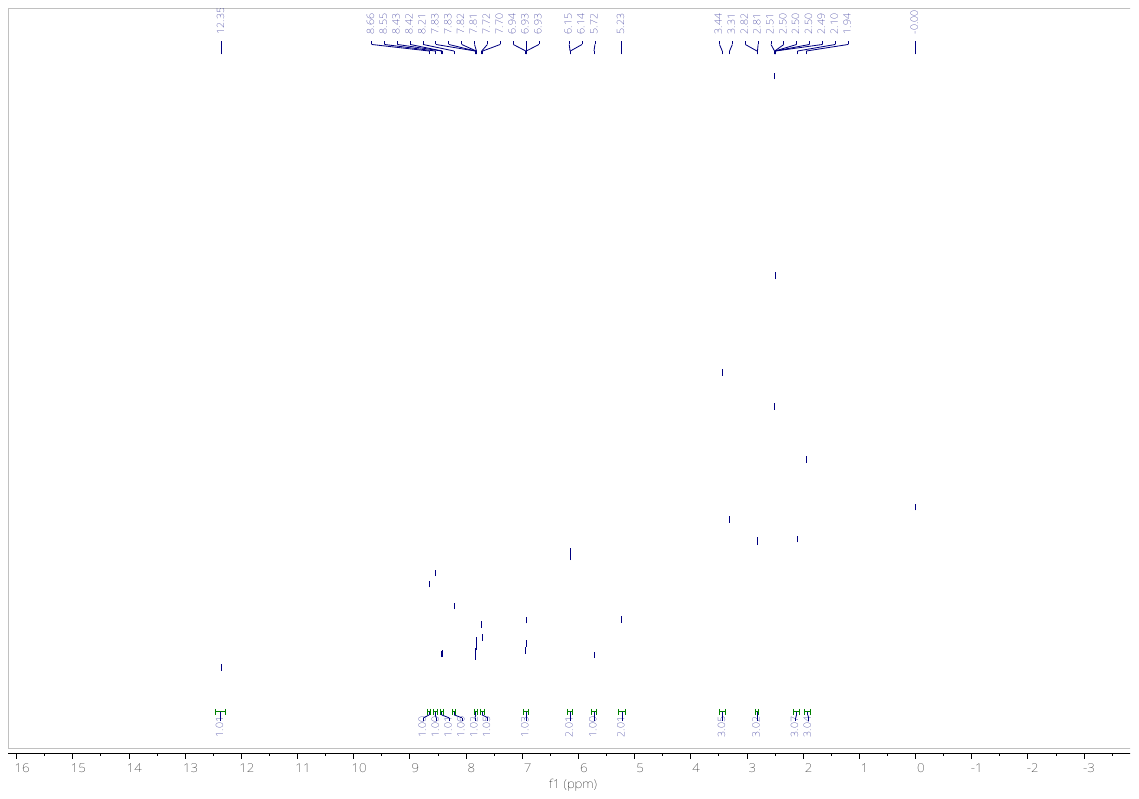

**Fig. 13**. ^1^H NMR of compound **28** in DMSO-*d*_6_.

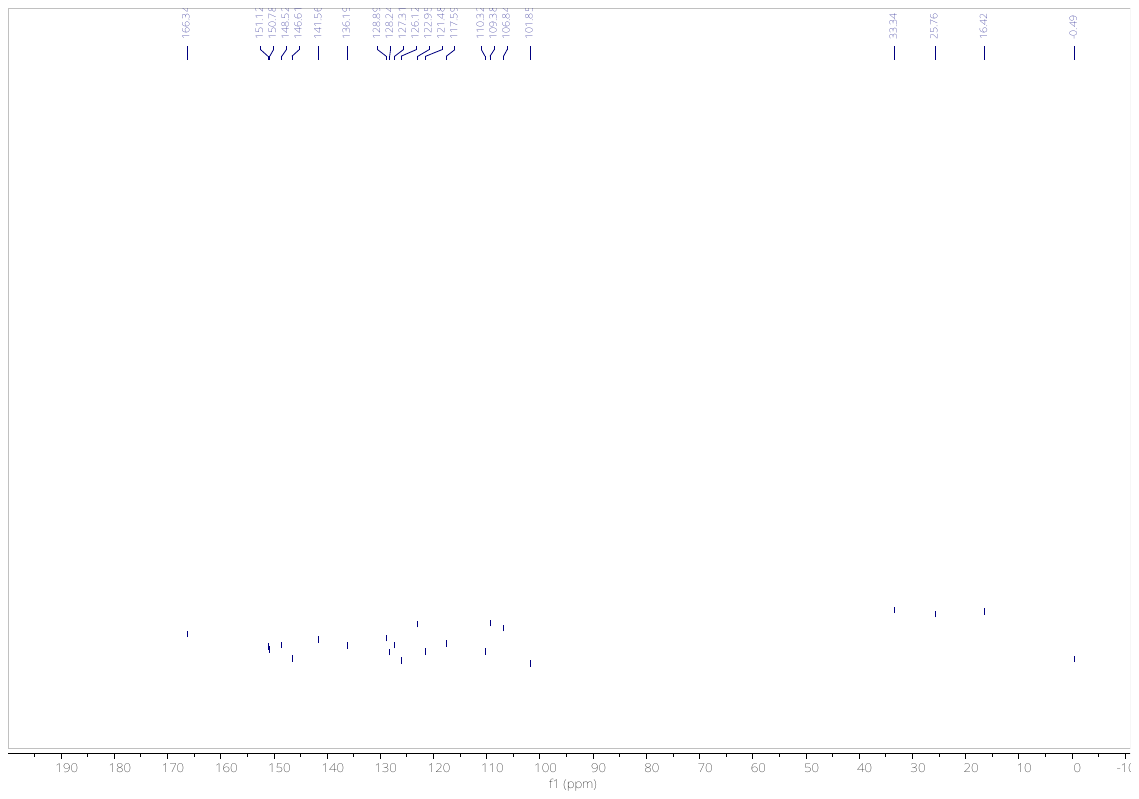

**Fig. 14**. ^13^C NMR of compound **28** in DMSO-*d*_6_.

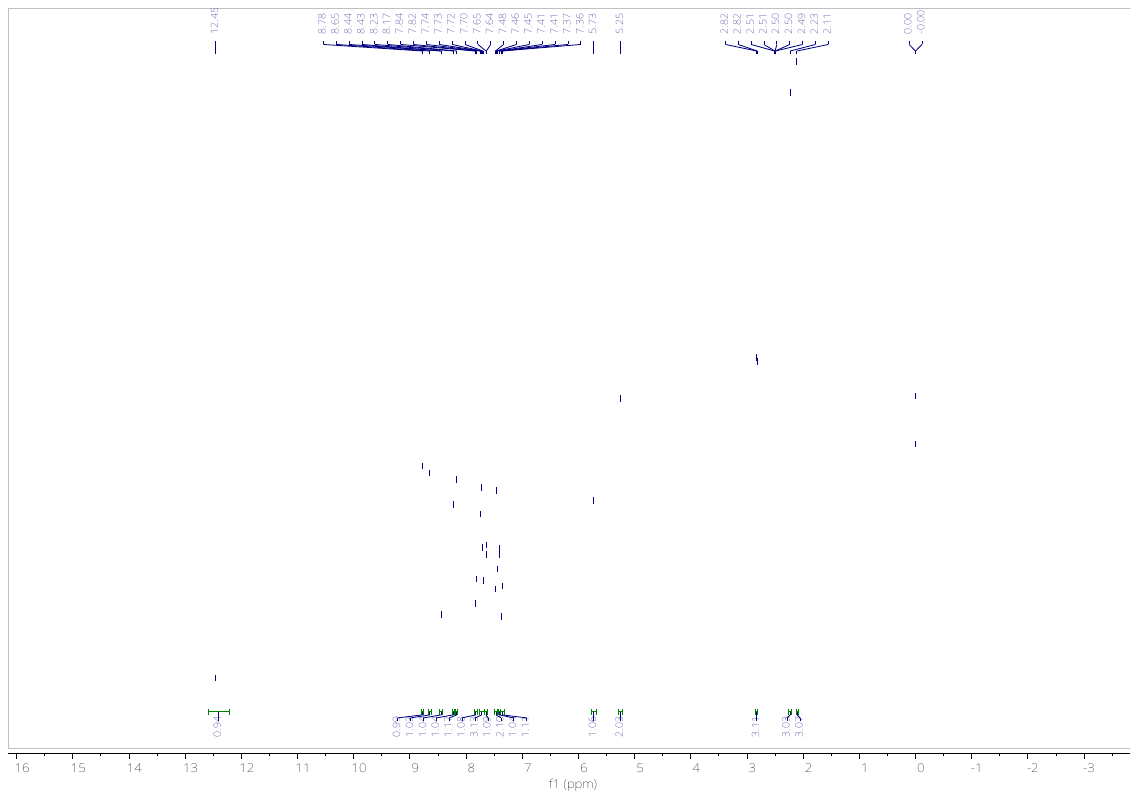

**Fig. 15**. ^1^H NMR of compound **29** in DMSO-*d*_6_.

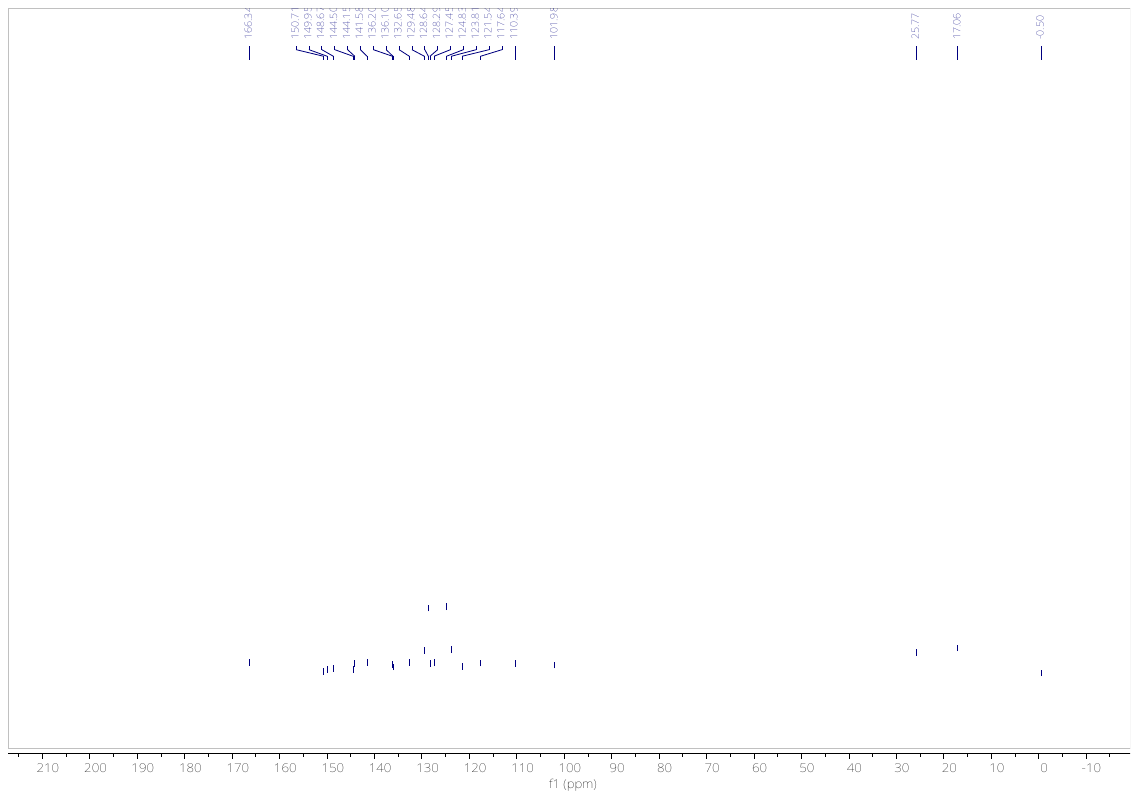

**Fig. 16**. ^13^C NMR of compound **29** in DMSO-*d*_6_.

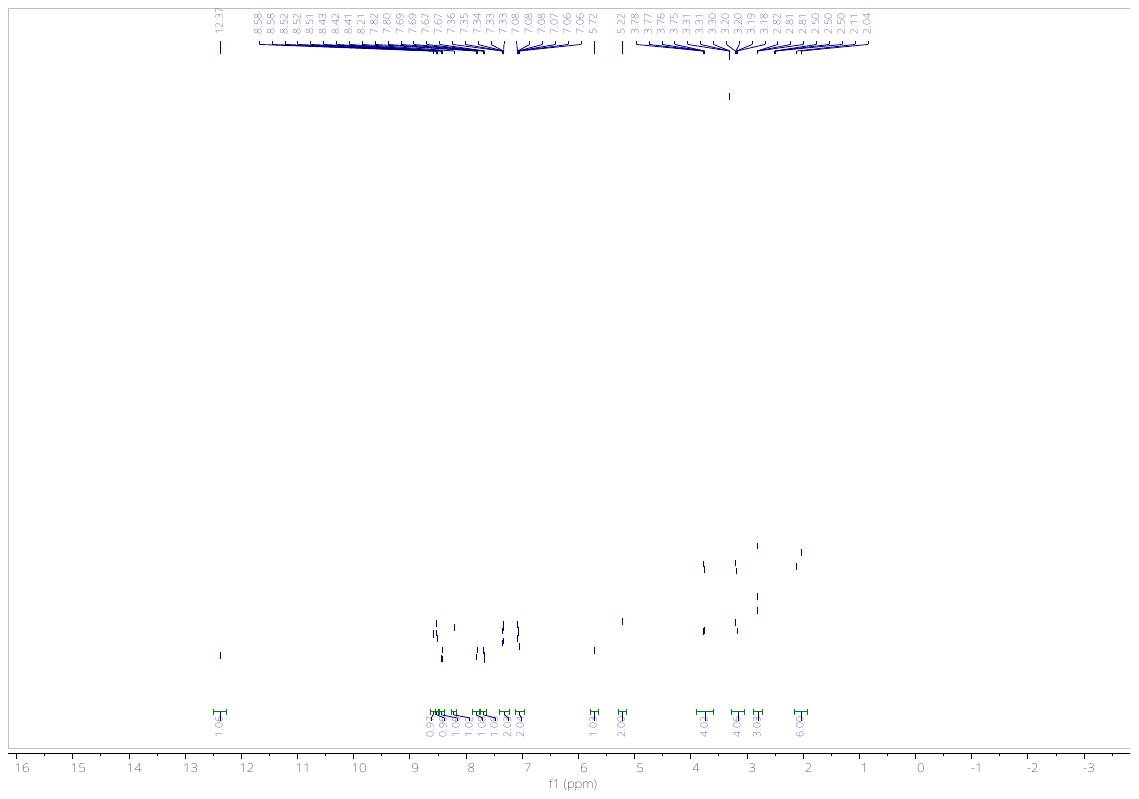

**Fig. 17**. ^1^H NMR of compound **30** in DMSO-*d*_6_.

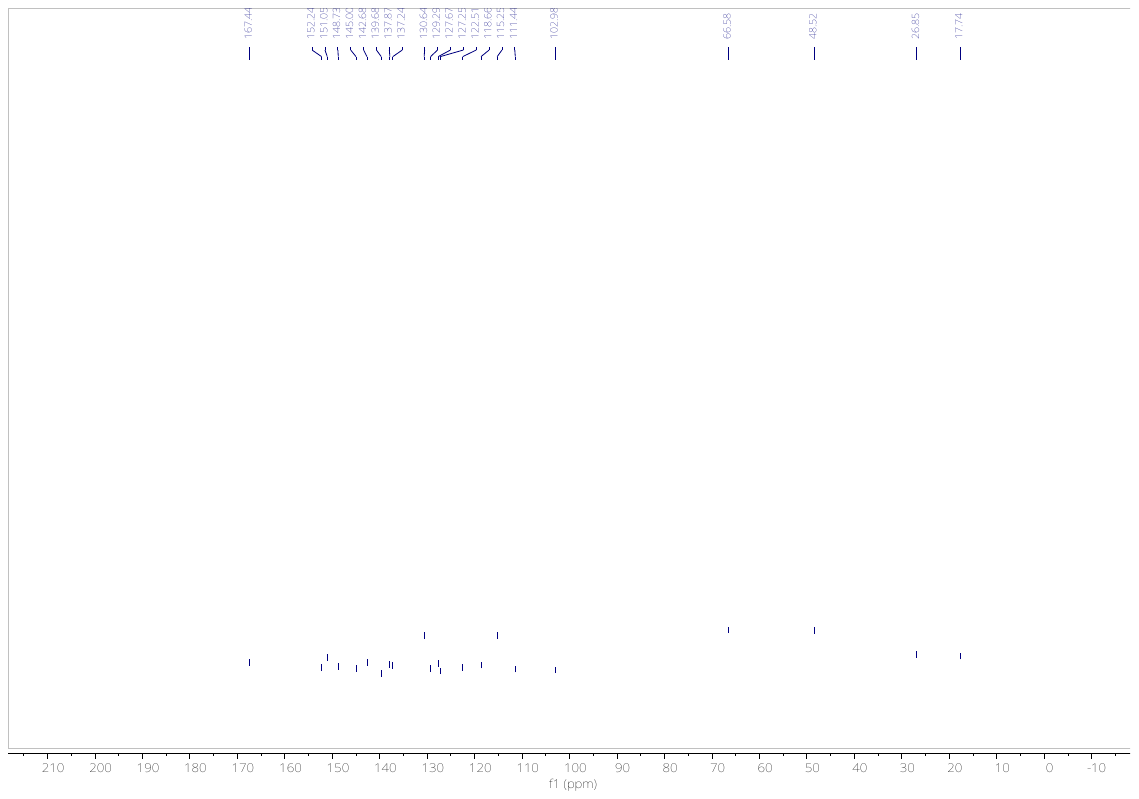

**Fig. 18**. ^13^C NMR of compound **30** in DMSO-*d*_6_.

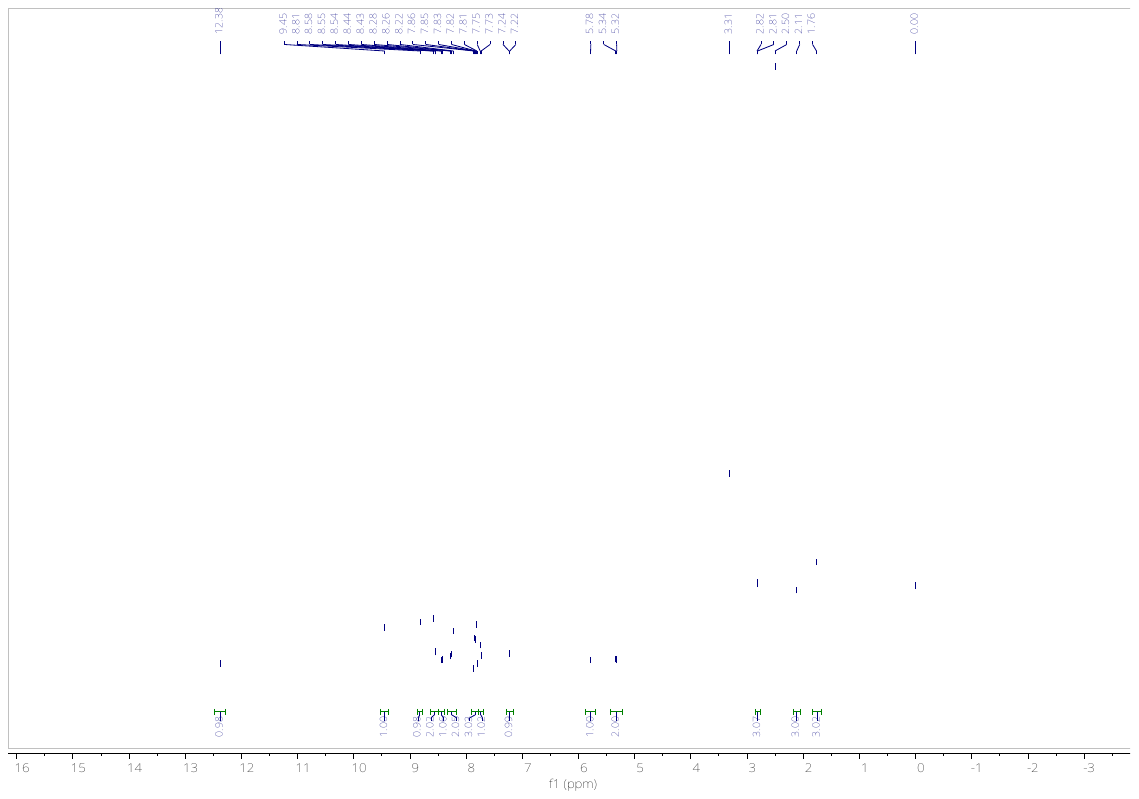

**Fig. 19**. ^1^H NMR of compound **31** in DMSO-*d*_6_.

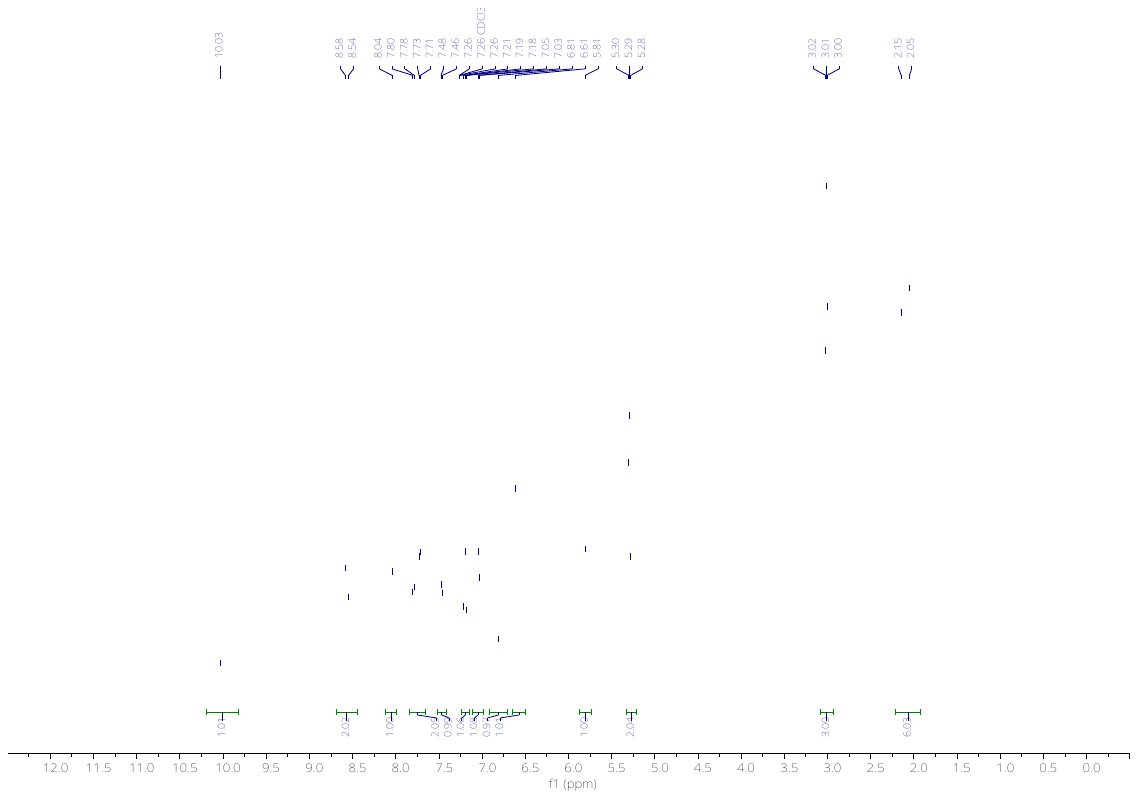

**Fig. 20**. ^1^H NMR of compound **32** in CDCl_3_.

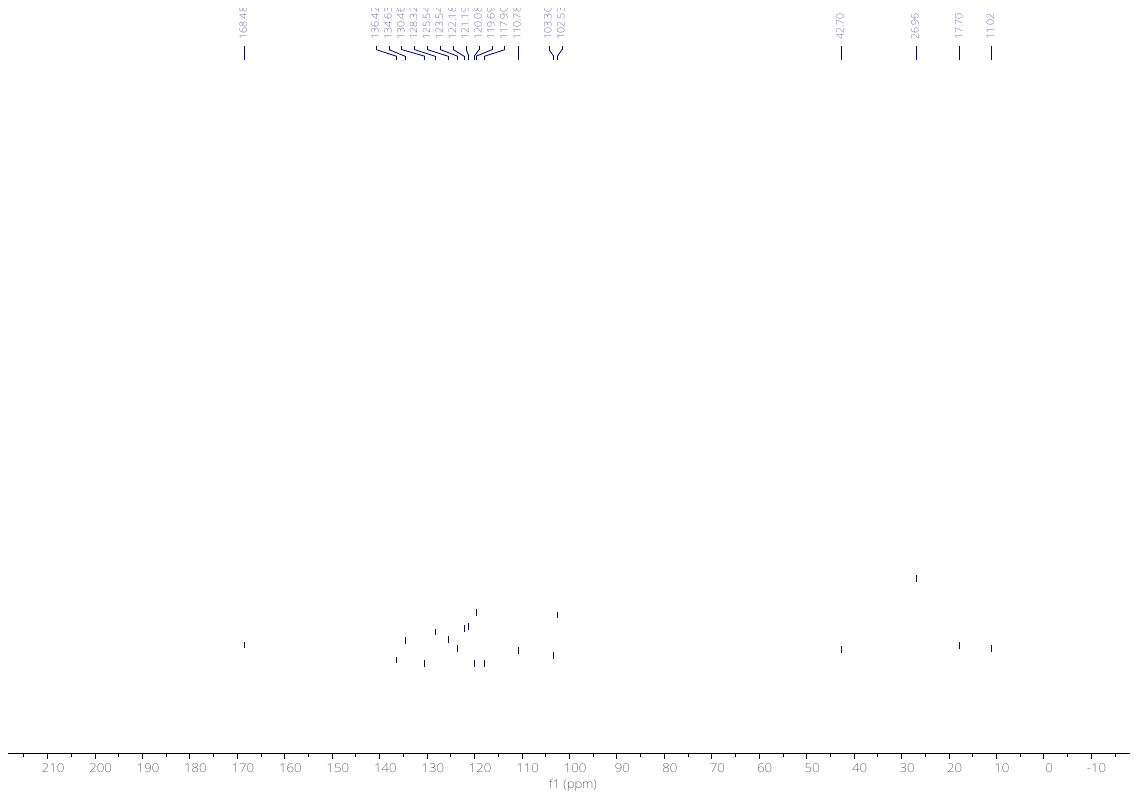

**Fig. 21**. ^13^C NMR of compound **32** in CDCl_3_.

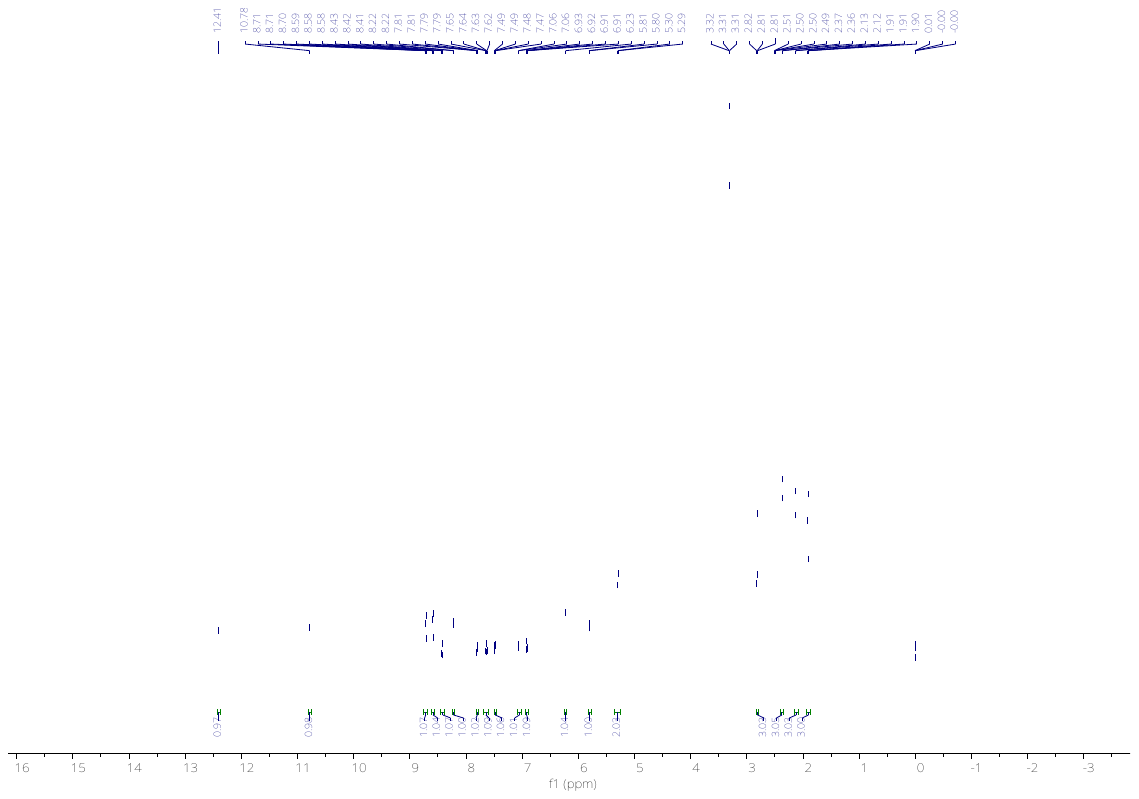

**Fig. 22**. ^1^H NMR of compound **33** in DMSO-*d*_6_.

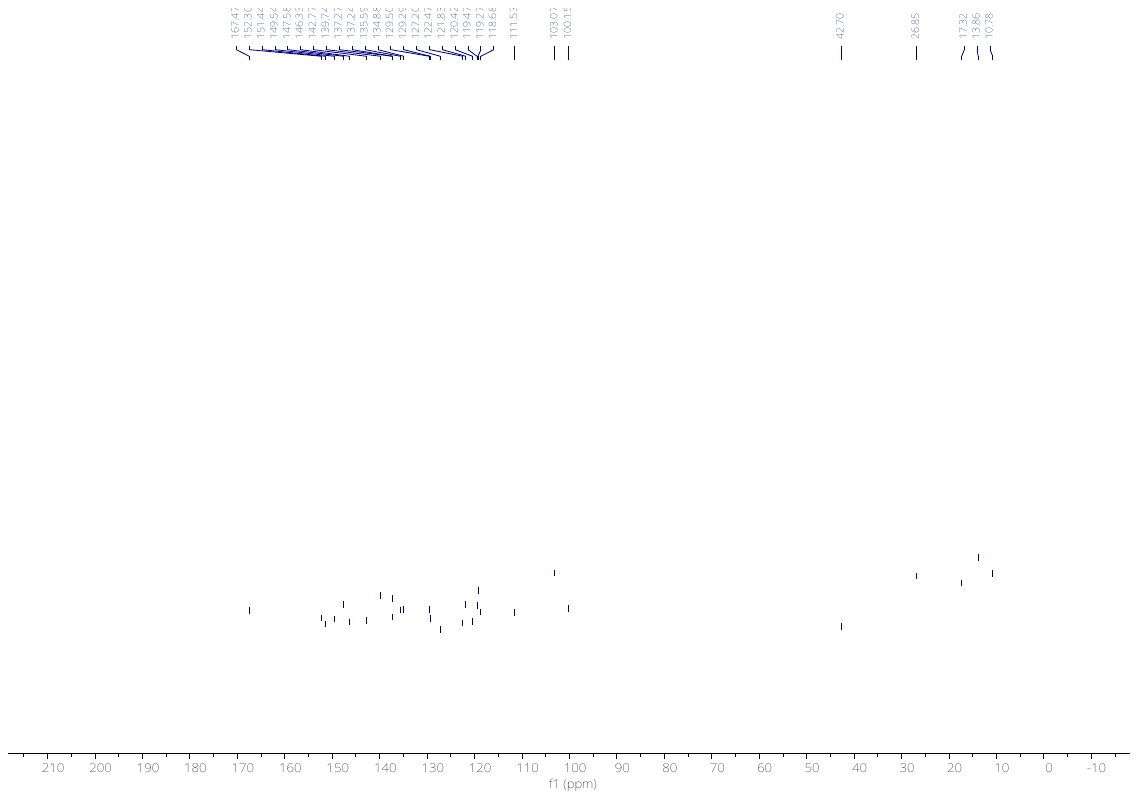

**Fig. 23**. ^13^C NMR of compound **33** in DMSO-*d*_6_.

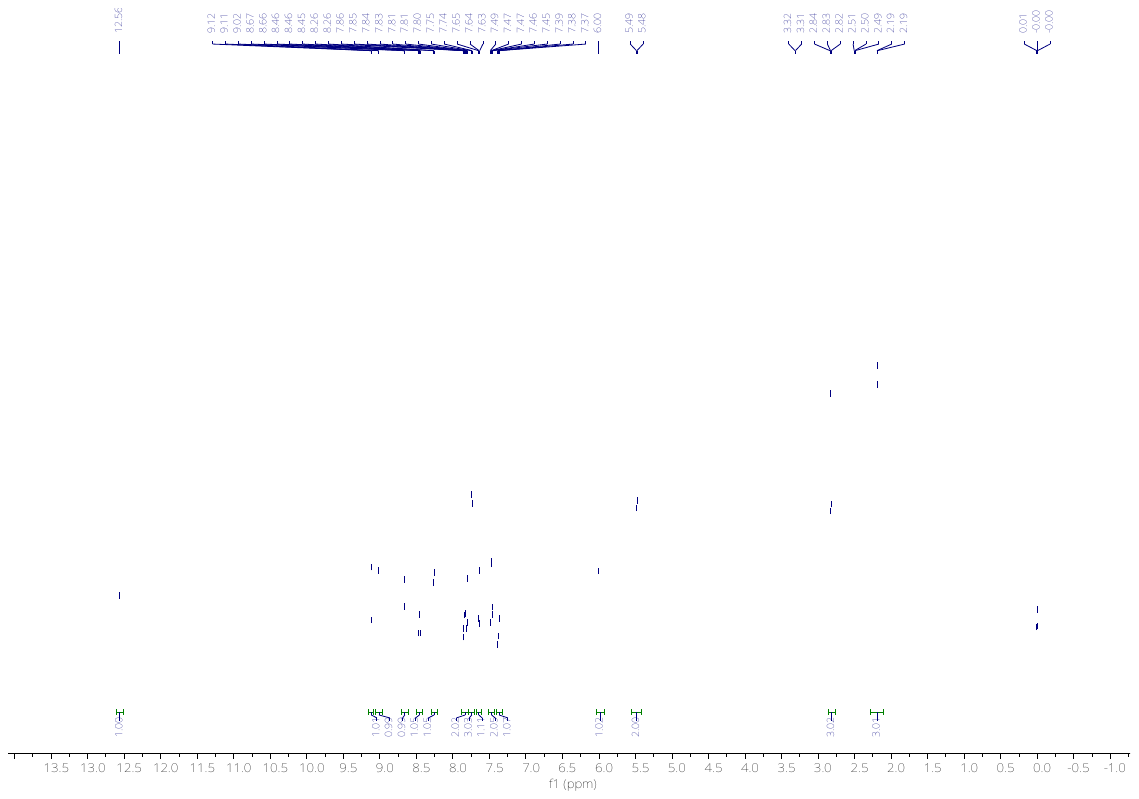

**Fig. 24**. ^1^H NMR of compound **44** in DMSO-*d*_6_.

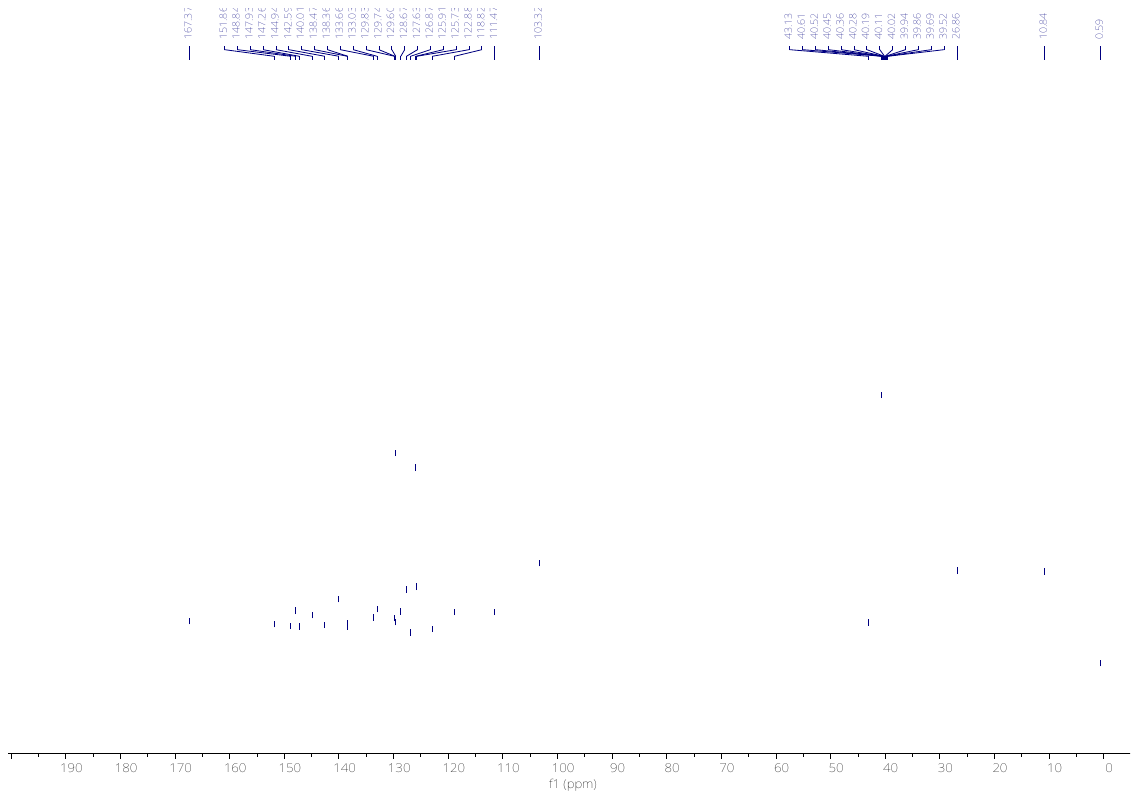

**Fig. 25**. ^13^C NMR of compound **44** in DMSO-*d*_6_.

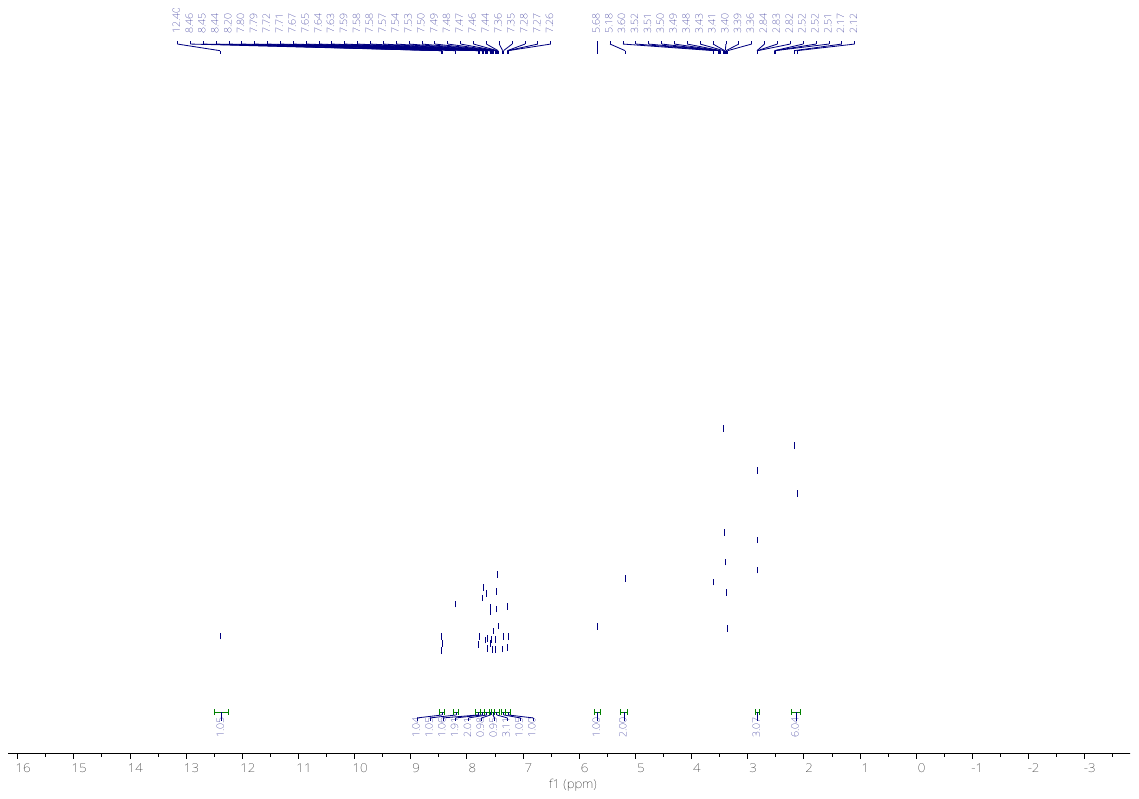

**Fig. 26**. ^1^H NMR of compound **45** in DMSO-*d*_6_.

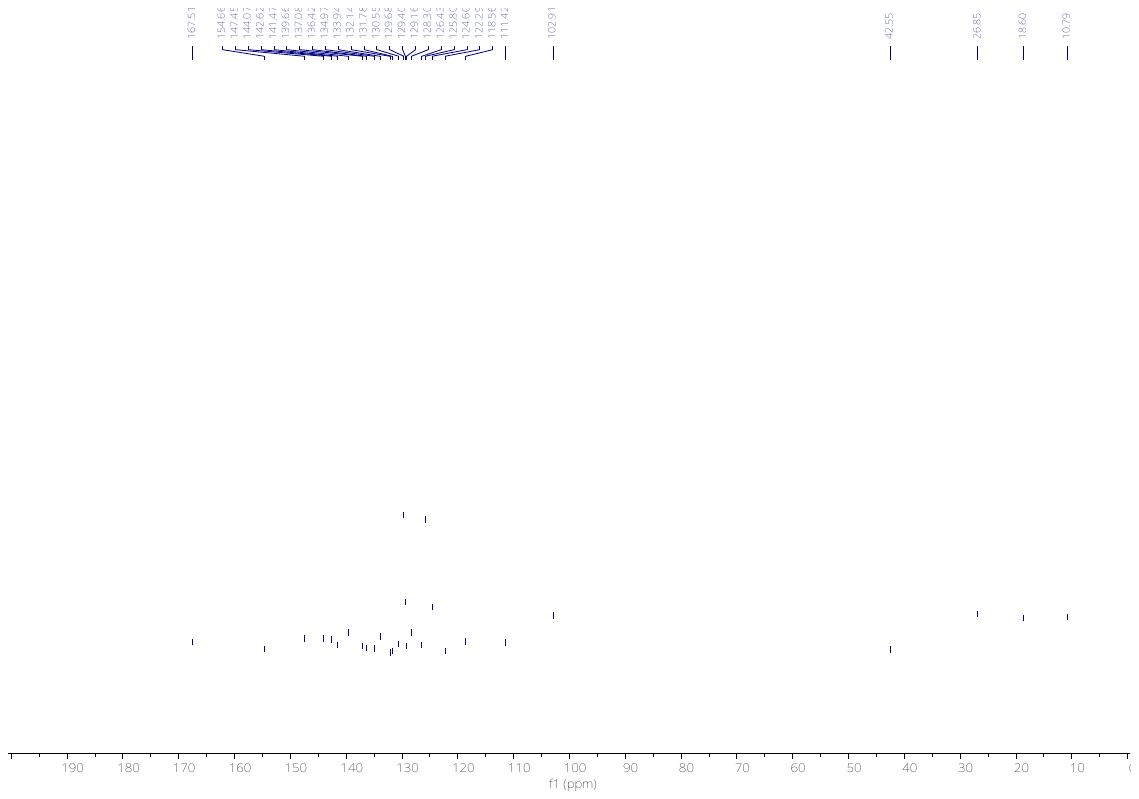

**Fig. 27**. ^13^C NMR of compound **45** in DMSO-*d*_6_.

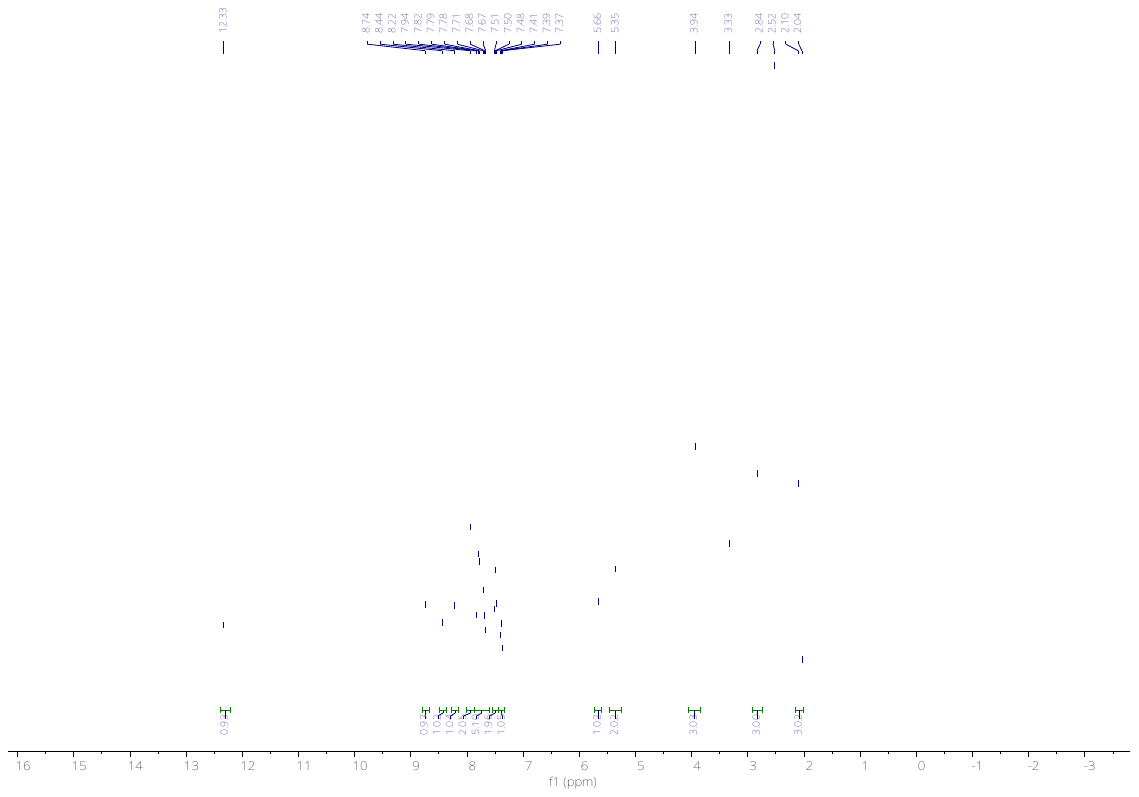

**Fig. 28**. ^1^H NMR of compound **46** in DMSO-*d*_6_.

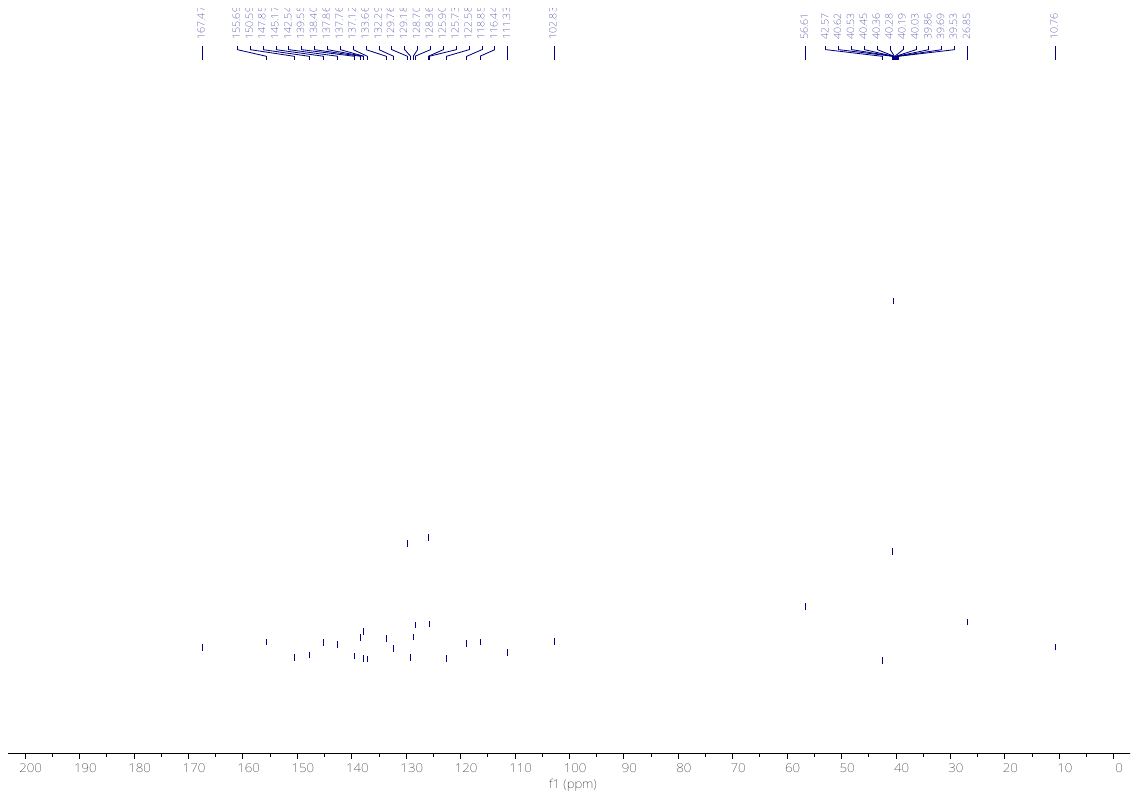

**Fig. 29**. ^13^C NMR of compound **46** in DMSO-*d*_6_.

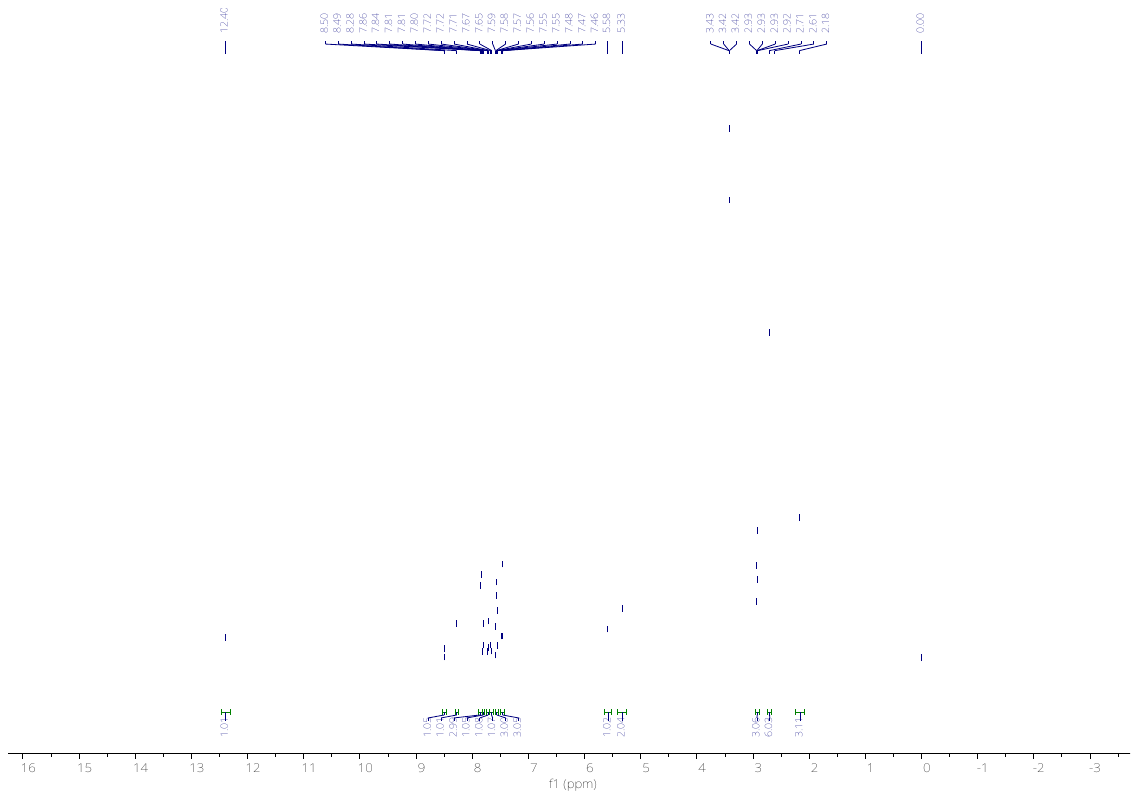

**Fig. 30**. ^1^H NMR of compound **47** in DMSO-*d*_6_.

**Fig. 31**. ^13^C NMR of compound **47** in DMSO-*d*_6_.

**Fig. 32**. ^1^H NMR of compound **48** in DMSO-*d*_6_.

**Fig. 33**. ^13^C NMR of compound **48** in DMSO-*d*_6_.

**Fig. 34**. ^1^H NMR of compound **49** in DMSO-*d*_6_.

**Fig. 35**. ^13^C NMR of compound **49** in DMSO-*d*_6_.

**Fig. 36**. ^1^H NMR of compound **50** in DMSO-*d*_6_.

**Fig. 37**. ^13^C NMR of compound **50** in DMSO-*d*_6_.

**Fig. 38**. ^1^H NMR of compound **51** in DMSO-*d*_6_.

**Fig. 39**. ^13^C NMR of compound **51** in DMSO-*d*_6_.

**Fig. 40**. ^1^H NMR of compound **52** in DMSO-*d*_6_.

**Fig. 41**. ^1^H NMR of compound **53** in DMSO-*d*_6_.

**Fig. 42**. ^13^C NMR of compound **53** in DMSO-*d*_6_.

**Fig. 43**. ^1^H NMR of compound **76** in DMSO-*d*_6_.

**Fig. 44**. ^13^C NMR of compound **76** in DMSO-*d*_6_.

**Fig. 45**. ^1^H NMR of compound **77** in DMSO-*d*_6_.

**Fig. 46**. ^13^C NMR of compound **77** in DMSO-*d*_6_.

**Fig. 47**. ^1^H NMR of compound **77** in CDCl_3_.

**Fig. 48**. ^1^H NMR of compound **78** in DMSO-*d*_6_.

**Fig. 49**. ^13^C NMR of compound **78** in DMSO-*d*_6_.

**Fig. 50**. ^1^H NMR of compound **79** in DMSO-*d*_6_.

**Fig. 51**. ^13^C NMR of compound **79** in DMSO-*d*_6_.

**Fig. 52**. ^1^H NMR of compound **81** in DMSO-*d*_6_.

**Fig. 53**. ^13^C NMR of compound **81** in DMSO-*d*_6_.

**Fig. 54**. ^1^H NMR of compound **82** in DMSO-*d*_6_.

**Fig. 55**. ^13^C NMR of compound **82** in DMSO-*d*_6_.

**HRMS:**

**Fig. 56.** HRMS of compound **10**.

**Fig. 57.** HRMS of compound **13**.

**Fig. 58.** HRMS of compound **24**.

**Fig. 59.** HRMS of compound **25**.

**Fig. 60.** HRMS of compound **26**.

**Fig. 61.** HRMS of compound **27**.

**Fig. 62.** HRMS of compound **28**.

**Fig. 63.** HRMS of compound **29**.

**Fig. 64.** HRMS of compound **30**.

**Fig. 65.** HRMS of compound **31**.

**Fig. 66.** HRMS of compound **32**.

**Fig. 67.** HRMS of compound **33**.

**Fig. 68.** HRMS of compound **44**.

**Fig. 69.** HRMS of compound **45**.

**Fig. 70.** HRMS of compound **46**.

**Fig. 71.** HRMS of compound **47**.

**Fig. 72.** HRMS of compound **48**.

**Fig. 73.** HRMS of compound **49**.

**Fig. 74.** HRMS of compound **50**.

**Fig. 75.** HRMS of compound **51**.

**Fig. 76.** HRMS of compound **52**.

**Fig. 77.** HRMS of compound **53**.

**Fig. 78.** HRMS of compound **76**.

**Fig. 79.** HRMS of compound **77**.

**Fig. 80.** HRMS of compound **78**.

**Fig. 81.** HRMS of compound **79**.

**Fig. 82.** HRMS of compound **80**.

**Fig. 83.** HRMS of compound **81**.

**Fig. 84.** HRMS of compound **82**.

**HPLC Chrmatograms of compounds:**

**Fig. 85.** HPLC Chromatogram of compound **10**.

**Fig. 86.** HPLC Chromatogram of compound **13**.

**Fig. 87.** HPLC Chromatogram of compound **29**.

**Fig. 88.** HPLC Chromatogram of compound **31**.

**Fig. 89.** HPLC Chromatogram of compound **49**.

**MST binding curve for the DEL hit:**

**Fig. 90. MST dose-response for the DEL-derived hit 10 against recombinant human NOS1AP over 15.3 nM-250 µM (n = 3).** No binding isotherm could therefore be fitted, and K_D_ is reported as >50 µM.

**Fig. 91. HS1 ranks first for each probe tested.**

Occupancy of each of the twelve hotspots (HS) by each probe separately, expressed as a percentage of that probe’s total occupancy. Isoquinoline, benzothiazole and pyridine each place HS1 first and HS2 second, so the ranking is not driven by the chemistry of any single probe.

**Table S1. Experimental PTB-domain structures used to validate the NOS1AP model and to define the composite peptide-binding groove**

Structures were identified by a sequence search over the NOS1AP PTB domain (residues 15–176) and superposed onto the model with structalign. Sequence identity was computed by local pairwise alignment (BLOSUM62, gap penalties −11/−1) between the NOS1AP PTB sequence and the longest chain of the superposed structure, over the aligned region. Cα RMSD is to the model. For peptide complexes, "model residues contacted" counts model residues with a heavy atom within 5.0 Å of the transferred peptide; residues meeting this criterion in at least two of the five complexes define the composite groove.

| **PDB** | **Protein and complex** | **Type** | **Sequence identity (%)** | **Cα RMSD (Å)** | **Model residues contacted** |
| --- | --- | --- | --- | --- | --- |
| 6ITU | GULP1 PTB domain with APP peptide (X-ray, human) | peptide complex | 35 | 1.68 | 20 |
| 3SO6 | ARH/LDLRAP1 PTB domain with LDLR tail (X-ray) | peptide complex | 37 | 1.71 | 23 |
| 8YM2 | AIDA-1 PTB domain with SynGAP NPxF motif (X-ray) | peptide complex | 32 | 1.78 | 25 |
| 5YI7 | Numb PTB domain with Pon peptide (X-ray, Drosophila) | peptide complex | 31 | 2.29 | 18 |
| 1DDM | Numb PTB domain with NAK peptide (NMR) | peptide complex | 31 | 2.70 | 21 |
| 5NJJ | Human Numb isoform-1 PTB domain, apo (X-ray) | apo | 32 | 2.37 | – |
| 3F0W | Numb-like PTB domain, apo (X-ray, human) | apo | 33 | 2.48 | – |
| 2M38 | AIDA-1 PTB domain, apo (NMR) | apo | 33 | 2.73 | – |
